# Coding agents author interpretable single-cell embedding models from the literature

**DOI:** 10.64898/2026.07.07.737048

**Authors:** Niklas Brunn, Sonia Maria Krißmer, Maximilian Frosch, Markus Frick, Marco Prinz, Harald Binder

## Abstract

The single-cell literature catalogs cell states as validated marker-gene programs — a sparse, compositional prior. Conventional embedding methods do not leverage this prior and learn cell-state structure de novo from the expression matrix, producing dense dimensions needing post-hoc interpretation and batch correction. Here we show coding agents can author single-cell embedding models directly from the literature. Given a scenario that focuses this literature lens on a chosen biological subdomain, the agent edits a structured Python template, curating named, literature-cited gene programs and composing them into axes, without a gene-set database, training, or sight of the data. Across mouse and human tissues these zero-shot embeddings are competitive in biological quality with conventional, foundation-model, and program-informed baselines, batch-robust by construction and reproducible across runs, complementing data-driven embeddings. Because each dimension is a named, cited gene program, the embedding is interpretable and auditable, and its composable axes can be steered into a developmental tree.

## Introduction

Reducing each cell’s high-dimensional expression profile to a compact, low-dimensional embedding is a fundamental step in single-cell RNA-sequencing (scRNA-seq) analysis, underlying how cells are visualized, clustered, annotated, integrated, and ordered along trajectories (Luecken and Theis, 2019; Heumos et al., 2023). Each cell is measured across thousands of genes, but these counts are sparse and noisy and the biological signal spans far fewer dimensions, so a good embedding suppresses technical noise and exposes the underlying cell-state structure. Most methods learn this representation directly from the expression data, viewing cells through the lens of the data itself (Lopez et al., 2018). In parallel, the scientific literature already encodes a second, complementary representation of the same biology, a literature lens through which the same cells can be viewed. It describes each cell state primarily as a marker-gene program, a small set of genes enriched in that state (Pasquini et al., 2021), alongside related gene-set forms such as signatures (Liberzon et al., 2015), co-expression modules (Langfelder and Horvath, 2008; Kotliar et al., 2019), and curated atlases and ontologies that pair cell types with their markers (Diehl et al., 2016; Franzén et al., 2019). Because these programs come from experimentally validated studies, each reflects a biologically defined identity rather than a technical artifact of measurement (Hu et al., 2023). This literature representation is therefore sparse, compositional, and human-readable, yet it is scattered across many papers and databases and cannot be used directly as an embedding. Here, we show that a coding agent can assemble it into one.

Conventional scRNA-seq embedding methods set this prior aside and re-learn cell-state structure from the expression data alone, producing a dense latent space whose dimensions are complex functions of many genes and require post-hoc interpretation (Lopez et al., 2018). Learning purely from data also leaves them sensitive to technical variation such as batch effects, so a dedicated correction step is needed when pooling samples (Luecken et al., 2022). Single-cell foundation models instead bring prior knowledge through large-scale pre-training on cell atlas corpora (Abdulla et al., 2025), promising transferable representations (Rosen et al., 2026). Yet used without fine-tuning they often fail to outperform simpler baselines (Kedzierska et al., 2025), and enlarging their pre-training corpora yields little additional gain (DenAdel et al., 2026).

A different strategy keeps the literature prior explicit, building the embedding directly from curated gene programs. Gene-set scoring (UCell (Andreatta and Carmona, 2021)) and program-anchored embedding models (expiMap (Lotfollahi et al., 2023)) describe cells through named, interpretable gene programs, but draw those programs from fixed, human-curated databases (PanglaoDB, Reactome (Franzén et al., 2019; Gillespie et al., 2022)). None authors its own programs from the literature for the scenario at hand, and these human-supplied programs form a flat representation that cannot be composed into structured axes or steered into a chosen topology. A related approach instead brings language-model knowledge of genes and cells to single-cell tasks (GenePT (Chen and Zou, 2025), scELMo (Liu et al., 2026)), but encodes it as dense embeddings rather than an interpretable, editable embedding whose dimensions are named gene programs.

Here we show that coding agents can author interpretable single-cell embedding models directly from the literature, given only a description of the biological scenario. Because that scenario selects which biological subdomain the agent draws on, the user chooses where to focus this literature lens, and can steer and inspect what knowledge enters the embedding. Coding agents extend LLMs with reasoning, tool use, and code execution, so they can iteratively write, run, and revise programs to resolve complex, multi-step tasks (Yang et al., 2024). They have begun to accelerate scientific discovery, from algorithm design (Romera-Paredes et al., 2024) to the discovery of single-cell analysis methods (Aygün et al., 2026). These capabilities fit the task of authoring an embedding model, which requires retrieving and reasoning over the literature, editing a structured specification, and self-checking the result. Conceptually, we treat the literature as a latent model of cell states, from which the agent curates the part relevant to a given scenario. Working from this scenario, the agent edits a declarative Python template, the blueprint, to curate named, literature-cited gene programs and compose them into embedding axes, with no human-supplied gene-set database, no training, and without ever seeing the data. Constructing the embedding is then a deterministic, CPU-only readout over the curated programs. Because each dimension is a named axis composed of cited gene programs, the embedding is interpretable and auditable by construction. Unlike a fixed database, the programs are curated dynamically. Unlike a trained model, the result is a short, editable specification whose axes can be composed and steered into structured topologies.

We demonstrate the approach across mouse and human tissues. A single authoring run produces an embedding whose named axes faithfully discriminate the cell types they name. The resulting zero-shot embeddings are competitive in biological quality with conventional, foundation-model, and program-informed baselines, complementing the data-driven view rather than replacing it, while being batch-robust by construction, since they score cells only on curated marker genes. Across repeated authoring runs the embedding is reproducible, even though the agent selects partly different genes each time. The authored specification is a Python script, auditable down to individual genes, programs, and citations, and produced in minutes at kilobyte scale. A single natural-language instruction can steer the embedding’s topology, for example into a developmental tree whose depth tracks embryonic stage. In practice, applying the approach to a new dataset requires only a description of its biological scenario. We package this workflow into an interactive web application that runs locally and keeps the user’s data on their machine. There, a user authors a blueprint, audits and edits it directly in the browser, maps their data, and explores the named axes and their steered topologies. Our contribution is an autonomous, auditable way to produce interpretable embedding models, with capabilities that fixed-database or trained interpretable methods currently do not provide.

## Results

### Coding agents author interpretable, literature-grounded embedding models whose named axes are faithful

A coding agent authors an embedding model directly from a description of the biological scenario. Given only a scenario configuration and the declarative Python template, it edits a single function to compose the blueprint, and it never sees the expression data (Figure 1a). Editing only this one function constrains the agent’s output to the blueprint API rather than arbitrary code. This fixes the structural form of the model and confines run-to-run variation to its biological content, the genes and programs the agent selects, leaving every blueprint named, standardized, and auditable. The blueprint it returns is a short specification rather than a trained model. Every output dimension is a named axis built from gene modules, and each module is a small set of marker genes the agent curated from the literature and cited to its source. Constructing the embedding for a dataset is then a deterministic, CPU-only readout in which each cell is scored on the marker genes of each axis, without training and without a batch variable (Figure 1b). The full agentic pipeline is laid out in Supp. Fig. S1.

**Figure 1.**
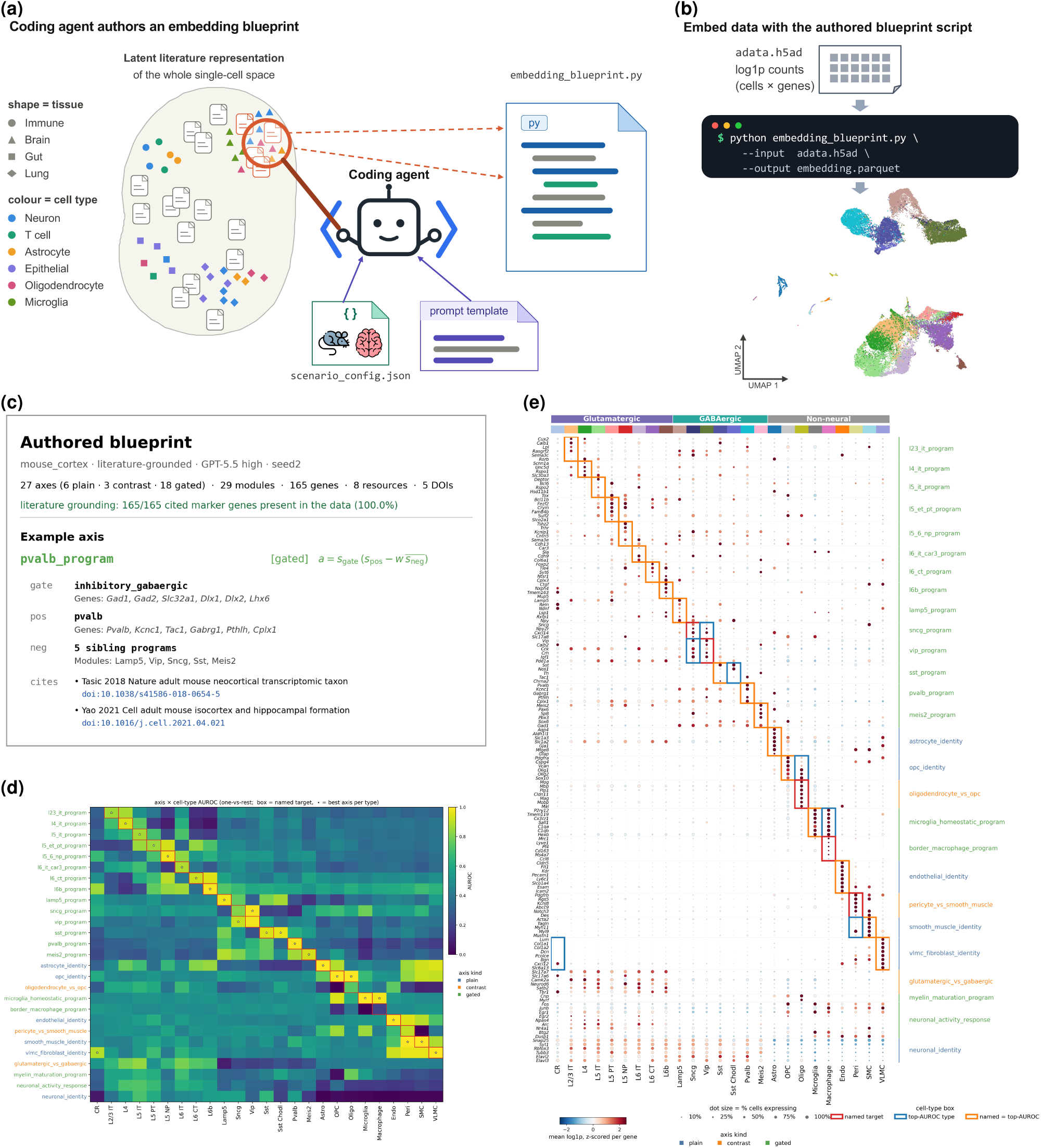
A coding agent authors an interpretable, literature-grounded embedding model. **(a)** Schematic of the agentic authoring step: from a latent literature representation of single-cell biology, the scenario configuration selects one biological subdomain (shape = tissue), and with a prompt template, the coding agent edits a single function of the declarative Python template to compose the blueprint from literature-cited gene programs, without seeing the data. (**b**) Applying the blueprint: a deterministic, CPU-only pass embeds a dataset’s log1p counts (.h5ad) into the cell-by-axis table, here represented by a UMAP plot of the mouse_cortex embedding colored by cell type. (**c**) The authored blueprint (mouse_cortex, literature-grounded, GPT-5.5 high): headline counts, the literature-grounding coverage, and one fully expanded example axis, showing its gated composition and positive/gate/negative gene modules with the cited gene symbols and DOIs. (**d**) Axis × cell-type AUROC matrix (one-vs-rest discrimination of each axis’s activation): axes are rows (colored by kind), cell types columns (shared with panel e). The red box marks the type each axis is named for and the white star the best axis per type. (**e**) Marker-gene dot plot: agent-curated marker genes (rows, grouped at right by their declaring axis) against cell types (columns, grouped on top by broad class). Dot color is the per-type mean log1p expression, *z*-scored across types per gene. Dot size is the fraction of cells expressing it. Outlines mark each axis’s named type (red) and each type’s top-AUROC axis (blue), orange where they coincide.

We instantiate this on an integrated mouse isocortex atlas we assembled from three studies (mouse_cortex (Tasic et al., 2016, 2018; Zeisel et al., 2018)). A single authoring run (Codex, GPT-5.5 at high reasoning effort) produced a 27-dimensional model, built from 29 named gene modules spanning 165 marker genes, each module citing peer-reviewed cortical atlases and ontology resources (8 resources, 5 DOIs, Figure 1c). The agent’s three inputs and the blueprint it authored for this run are reproduced in full in Supp. Code S1–S4. The blueprint API provides three kinds of axis (Methods). A plain axis reads out a single program, a contrast axis opposes two programs to separate populations at opposite ends of one distinction, and a gated axis restricts a program to the cells of a parent class, so chained gates encode hierarchical structure. The showcase model uses all three, with 6 plain identity axes, 3 contrasts, and 18 gated axes. Together they lay out the canonical organization of the cortex (Tasic et al., 2018). A pan-neuronal axis and a glutamatergic-versus-GABAergic contrast set the coarse split, gated programs resolve the glutamatergic layers (L2/3, L4, L5 IT/ET/NP, L6 CT/IT/b) and GABAergic interneuron classes (Lamp5, Vip, Sncg, Sst, Pvalb, Meis2), and further axes name the non-neuronal types (astrocyte, OPC, oligodendrocyte, microglia, border macrophage, endothelial, pericyte, smooth muscle, VLMC). The agent represents two of these non-neuronal types with a contrast axis rather than a plain identity axis, opposing oligodendrocyte to OPC and pericyte to smooth muscle. Every marker the agent cited is present in the data (165/165 genes), so the grounding is not merely nominal.

Because each axis is an explicit, named composition of cited gene modules, reading a dimension requires no post-hoc attribution, only its declared gene set. The embedding is therefore interpretable by construction, and it is also faithful to the data. By faithful we mean that each named axis discriminates the cell type it names. We quantify this faithfulness directly. For every axis we score how well its activation separates each annotated cell type (one-vs-rest area under the receiver operating characteristic curve, AUROC), and read this against the type the axis is named for (Methods). The named axes are sharply diagonal (Figure 1d). Almost all are brightest for the type they claim and dim elsewhere, and across the 23 axes that name a subclass the activation discriminates that subclass with mean AUROC 0.90 (median 0.95, with 16*/*23 above 0.90 and 21*/*23 above 0.80). The model names an axis for 23 of the 25 annotated subclasses. The two it does not target, Cajal–Retzius and the rare Sst Chodl interneuron, surface as the columns with no boxed axis, an explicit and inspectable gap rather than a silent one. The lone failing named axis (border macrophage, AUROC 0.20, below chance) shows as a dim boxed cell that a domain expert could audit. As a complementary view, assigning each cell to its most-activated axis recovers the annotated types along a clear diagonal (Supp. Fig. S2). The same faithfulness is visible gene by gene. Grouping the cited markers by the axis that declares them, and ordering cell types by broad class, yields a clean diagonal (Figure 1e). Each named axis’s genes are elevated, in both mean level and the fraction of cells expressing them, precisely in the cells of the type it names. The non-neuronal programs light up sharply and almost exclusively in their populations, and the layered glutamatergic and interneuron programs track their cortical subclasses. Each axis correspondingly localizes to a coherent region of the expression manifold (Supp. Fig. S3). This per-axis faithfulness is not an artifact of the single showcased run. Across all ten re-authoring seeds the per-type discrimination remains consistently high (Supp. Fig. S4).

### Blueprints reach competitive embedding quality and are batch-invariant by construction

We benchmark the blueprint against established embedding methods on three independent tissues across two species, an integrated mouse isocortex atlas (mouse_cortex) and the scIB human pancreas and human immune compendia. The three carry heterogeneous batch structure, spanning multiple studies, protocols, sequencing technologies, and donors, so batch robustness is tested across batch types rather than one technical source. Every method is scored with the same scIB suite of bio-conservation and batch-correction metrics (Luecken et al., 2022) (Methods). The blueprint carries biological prior knowledge that the unsupervised embeddings lack. Given only the species and tissue, the agent draws on the literature to decide which cell types occur in it and which markers define them, whereas the unsupervised methods learn cell structure de novo from the data. We benchmark the blueprint under both of its grounding variants, literature-grounded (every gene module must cite a literature resource) and resource-unconstrained (noref), where citations are not required. We tier the comparators by the prior knowledge they access (Methods). A prior-free floor (PCA and scVI (Lopez et al., 2018) without its batch key) accesses none, and scVI adds only the batch covariate. A zero-shot foundation-model tier adds generic pretraining on large cell atlases (scVI-census (Abdulla et al., 2025) and UCE (Rosen et al., 2026)). The most direct comparators are two program- and marker-based methods that, like the blueprint, consume explicit gene programs but take them from a human-curated database rather than authoring them from the literature. These are UCell (Andreatta and Carmona, 2021), training-free gene-set scoring on PanglaoDB (Franzén et al., 2019) markers, and expiMap (Lotfollahi et al., 2023), a trained interpretable VAE that corrects batch by conditioning on the batch key. The purpose of this benchmark is to assess the blueprint’s biological quality and batch robustness against these comparators.

The aggregated results across all datasets indicate that competitive bio-conservation and batch robustness are a general property of program- and marker-based models. Specifically, the blueprint (both grounding variants), UCell and expiMap form one tight cluster high on the batch axis, well above PCA, scVI-census, UCE and scVI without its batch key. Only scVI lies further to the top-right (Figure 2a). This clustering holds on each dataset individually, not only on the average (Supp. Fig. S5). On biological quality the blueprint matches these comparators. Averaged over the benchmark the three sit essentially on top of one another, and per dataset the blueprint ties them on the human pancreas (scIB Total 0.64–0.67), is overtaken on the human immune compendium (expiMap 0.66, UCell 0.64 versus the blueprint’s 0.57–0.58), and pulls ahead on the fine-grained mouse cortex.

**Figure 2.**
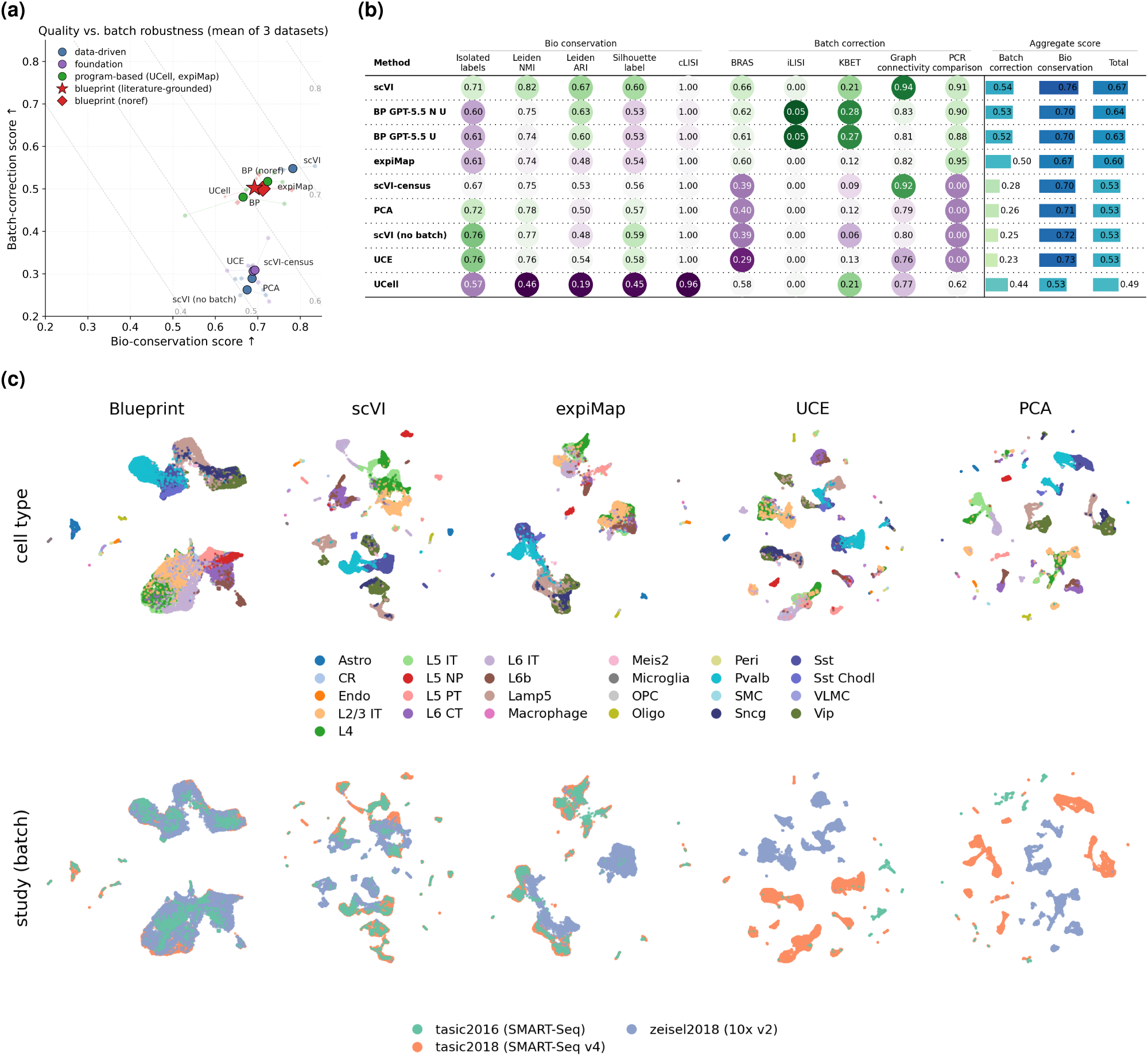
Blueprints reach competitive quality and are batch-invariant by construction. (**a**) Averaged trade-off of scIB bio-conservation against batch-correction, one marker per method, taken as the mean over the three benchmark datasets (mouse_cortex at the fine subclass resolution, human_pancreas, and human_immune). The three faint same-color points are the per-dataset values. Markers are colored by method class. The two blueprint grounding variants are shown in red, a star for the literature-grounded variant and a diamond for the resource-unconstrained (noref) variant. The program- and marker-based comparators UCell (training-free gene-set scoring) and expiMap (trained interpretable VAE) share one method class, shown in green. Dashed lines connect points of equal composite score (0.6 · bio + 0.4 · batch). (**b**) mouse_cortex scIB leaderboard at the fine subclass resolution, one row per method, mean across seeds, with per-metric cells and Bio/Batch/Total bars. The blueprint rows BP GPT-5.5 U and BP GPT-5.5 N U are the literature-grounded and resource-unconstrained (noref) variants. (**c**) mouse_cortex UMAPs at the best seed per method (based on scIB composite score), the top row colored by cell type and the bottom row by study (the batch).

On the mouse_cortex subclass labels (L2/3 IT, L5 IT/ET/NP, Pvalb, Sst, Vip and so on) the blueprint reaches scIB Total 0.63–0.64 (ARI 0.60–0.64), ahead of the trained expiMap (0.60, ARI 0.48) and UCell (0.49, ARI 0.19), while the prior-free floor and foundation tier sit near 0.53 and only scVI is higher (0.67, ARI 0.67). Figure 2b shows the mouse-cortex leaderboard, and the full numbers with error bars for every dataset and resolution are in Supp. Table 2. The mechanism is the program vocabulary. Neither fixed database resolves cortical subclasses. PanglaoDB, UCell’s source, offers only generic neurotransmitter and morphology classes such as glutamatergic, GABAergic, pyramidal, and interneuron, and expiMap’s Reactome and PanglaoDB sets add molecular pathways but still no subclass identities. UCell therefore cannot separate the fine subclasses, and training lets expiMap recover only partway. The agent is not bound to a fixed database, instead curating from the literature whatever programs fit the scenario and authoring roughly 23 named subclass axes that no database supplies. At the coarser three-class cortical resolution, where the database does carry adequate programs, the gap vanishes (UCell 0.61, blueprint 0.61, expiMap 0.58, scVI 0.56) and the blueprint separates the three coarse classes at least as cleanly as scVI while remaining fully interpretable (Supp. Table 2). The blueprint reaches this without training, a batch variable, or sight of the data.

The blueprint, UCell and expiMap reach the batch axis by different routes. All three mix batches well, with scIB batch-correction sub-scores of 0.44–0.54 across the three datasets, close to scVI (0.55) and well above the prior-free floor and foundation tier (0.24–0.38, Figure 2a). For the blueprint and UCell this invariance is by construction, whereas expiMap, like scVI, achieves it by correction, conditioning its trained latent on the batch key. For the blueprint the invariance follows from how the agent builds the model. It selects only genes the literature validates as markers of specific cell types and programs and groups them into named modules intended to track biological rather than technical variation. Because the embedding reads out only these curated modules, the genes and library-size shifts that carry batch and protocol effects have little dimension to load onto, so the blueprint cannot over- or under-correct a batch effect it never sees. scVI without its batch key drops to the prior-free models (batch ≈ 0.25), confirming that a conventional method depends on the batch variable for this mixing. The effect is visible directly in the embeddings (Figure 2c). On mouse_cortex, which concatenates three studies spanning full-length and droplet protocols, the blueprint, scVI and expiMap interleave the three studies within shared cell-type structure, whereas UCE and PCA separate cells by study, the technical split the blueprint structurally cannot represent. The same contrast holds in the human pancreas and immune embeddings (Supp. Fig. S6). The literature-grounded and data-driven embeddings recover the same shared cell-type structure while differing in how they reach it and in what they make explicit, so the two are complementary rather than interchangeable.

To simulate generalization to unseen data, we hold out an entire study, technology, or donor as a query on each benchmark dataset, the SMART-seq Tasic 2016 study for the cortex, the SMART-seq2 technology for the pancreas, and one bone-marrow donor with one PBMC sample for the immune compendium. scVI and expiMap are fit on the reference and mapped onto the query by a frozen pass or a query-side scArches fine-tuning step (Lotfollahi et al., 2022). Scoring reference-to-query label transfer with a *k*-nearest-neighbor classifier and the joint-embedding scIB suite (Supp. Fig. S7, S8), the zero-shot blueprint is competitive across all three tissues. It attains the highest accuracy on human_pancreas (0.97 versus 0.95 for scVI and 0.92 for expiMap), ties scVI on mouse_cortex (0.79, with the best macro-*F*_1_, 0.53 versus 0.46, recovering rare subclasses the others blur), and on the immune compendium trails only scVI (0.84 versus 0.88) while staying ahead of expiMap (0.78), both variants transferring indistinguishably. Consistent with the above benchmarking results, scVI retains the highest joint-embedding scIB Total throughout (0.69 versus the blueprint’s 0.63 on the cortex).

### A structured agentic framework yields reproducible, auditable embeddings

The blueprint forces the agent to build the embedding through a fixed API rather than arbitrary code, composing cited gene modules into typed axes. This confines run-to-run variation to biological content and keeps every run comparable and auditable. We ask whether the structure costs biological quality, and whether agent authoring is reproducible. On a representative dataset with clean, canonical annotations (human pancreas), we compare the structured blueprint against free-form prompting, an arbitrary script bound only to the same cell-by-axis output contract, at matched content (literature-grounded, naïve, and resource-unconstrained variants; blueprints in both Stage A and refined states; Codex, GPT-5.5, high reasoning effort, 10 runs each; Figure 3). The results demonstrate that structure is not a quality cost. Blueprint cohorts match or exceed the matched free-form cohorts on scIB (Luecken et al., 2022) bio-conservation (Figure 3a) and stay at least comparable on batch correction (Figure 3b), whereas naïve free-form reaches competitive cell-type scores only by collapsing batch mixing. The structure also supplies auditable handles that free-form lacks. Each blueprint emits a bounded set of named gene modules (Figure 3d), axes (Figure 3e), and a citation trail (Figure 3f), with gene content at least as reproducible as free-form (Figure 3c) and a far tighter dimensionality (naïve free-form spans ∼26 to 99 axes versus the blueprint’s 16–45).

**Figure 3.**
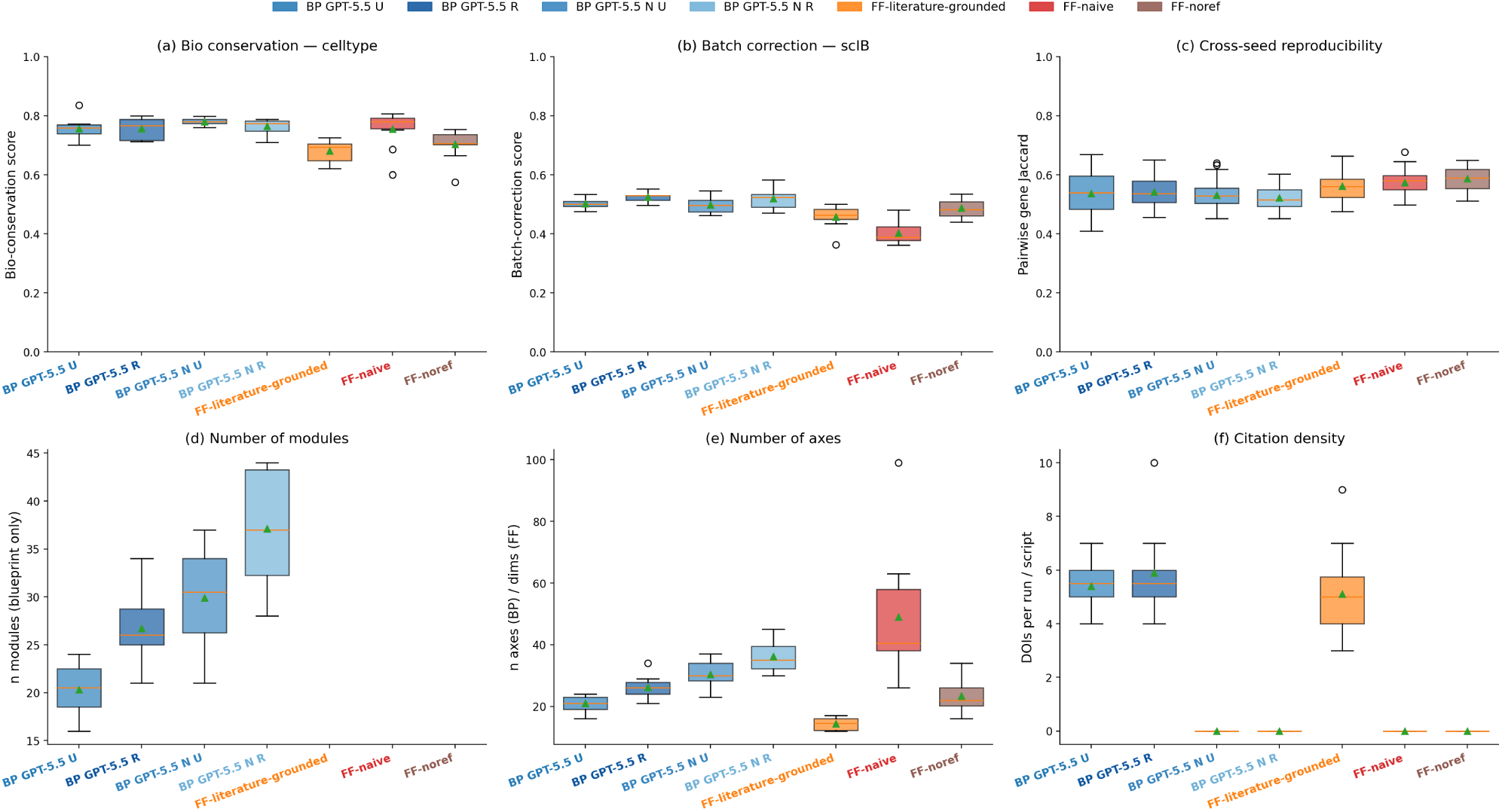
What the structured blueprint framework provides over free-form prompting. Blueprint variants (BP) versus free-form variants (FF) of the agentic pipeline at matched content (human pancreas, Codex, GPT-5.5 at high reasoning effort, 10 runs each). In the blueprint labels, U denotes unrefined (Stage A), R refined (Stage A+B), and N resource-unconstrained (noref). (**a**) scIB bio-conservation sub-score (mean of NMI, ARI, ASW-label, cLISI, isolated-label score) and (**b**) scIB batch-correction sub-score (mean of graph connectivity, ASW-batch, iLISI, kBET, PCR). (**c**) Cross-seed reproducibility as the per-pair gene-Jaccard distribution. (**d**) Number of gene modules (blueprint-only structural property). (**e**) Number of embedding axes (blueprint) / dimensions (free-form). (**f**) Citation density (DOIs per run/script). Boxes show the median and interquartile range, whiskers 1.5×IQR, and the triangle the mean. All distributions are over the 10 runs.

The embedding is reproducible across runs even though the agent picks partly different genes each time. We quantify this over 10 re-authoring runs at three levels, the specification, the embedding, and the biological conclusions, all on the same cells. The specifications show a moderate overlap in their genes (mean pairwise Jaccard 0.53). Yet, the embedding geometry is highly reproducible, with mean pairwise linear centered-kernel-alignment (CKA; (Kornblith et al., 2019)) of 0.85 (Figure 4a), so runs choose different genes for the same programs but arrive at essentially the same embedding. Benchmarked against a retrained neural network method, blueprint embeddings are at least as reproducible as scVI across seeds (CKA 0.82–0.85 versus 0.74, non-overlapping 95% bootstrap CIs) and far more reproducible than free-form (0.52–0.59; Figure 4b). At the conclusion level their cell-type quality is likewise consistent (scIB ARI s.d. 0.06–0.12 across seeds versus 0.10–0.20 for free-form). This does not depend on the most capable engine. A smaller model (GPT-5.4-mini) reaches comparable quality and still-high reproducibility (CKA 0.77–0.83). Reproducibility alone is not enough. In contrast, an unintegrated baseline (scVI without its batch key) is highly reproducible (CKA 0.82) yet recovers little structure (ARI 0.35).

**Figure 4.**
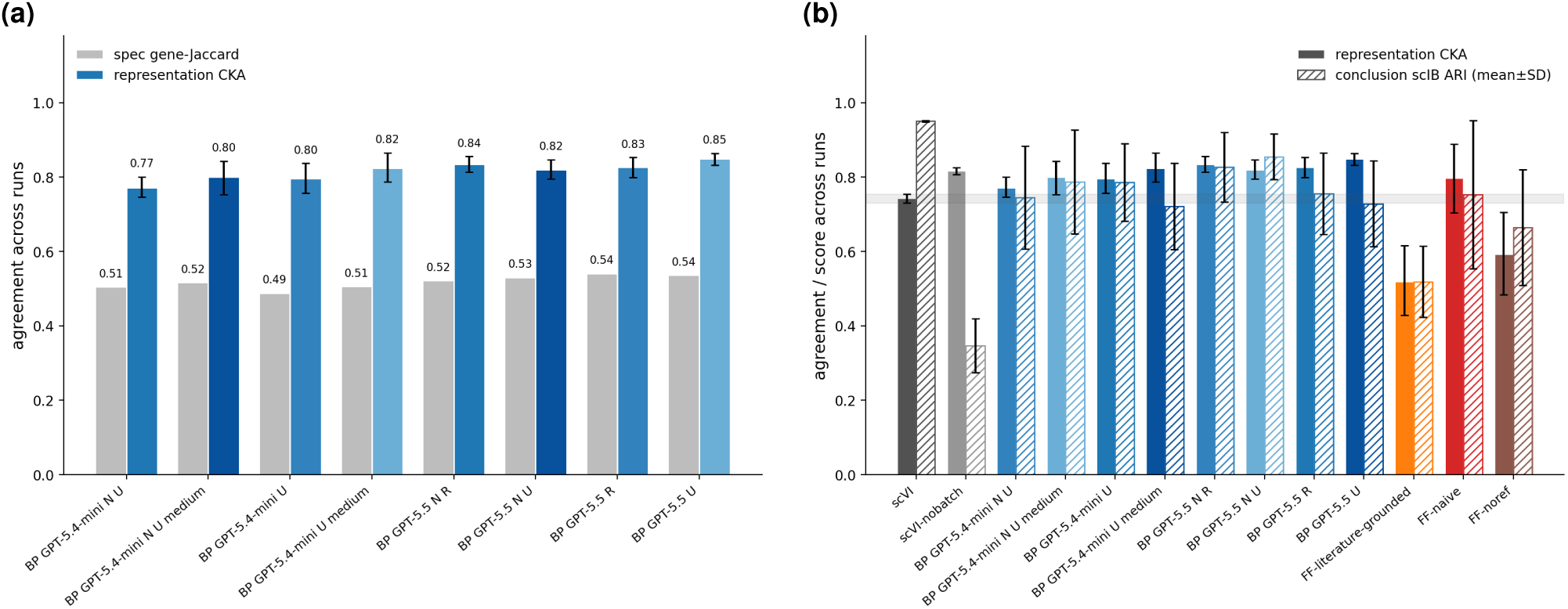
Variation in the agent-authored specification does not propagate to the embedding. **(a)** For each blueprint cohort (9–10 runs), agreement across runs at the specification level (mean pairwise gene Jaccard) versus the representation level (mean pairwise linear CKA). In the blueprint labels, U denotes unrefined (Stage A), R refined (Stage A+B), N resource-unconstrained (noref), and a trailing effort token (e.g. medium) the reasoning effort when it is not high. (**b**) Reproducibility across all multi-seed cohorts: scVI baselines, blueprint variants from two authoring engines (GPT-5.5 at high reasoning effort and the smaller GPT-5.4-mini at high and medium reasoning effort), and free-form variants. Solid bars are the representation CKA with run-level bootstrap 95% CIs, and the grey band marks scVI’s CKA interval. Hatched bars are the conclusion-level scIB ARI-vs-truth (mean ± s.d. across seeds), the s.d. whisker being the quality-reproducibility signal.

To see what composes reproducibility, and what makes a blueprint auditable where a free-form script is not, we ask which elements of the specification recur across the 10 runs (Supp. Figs. S9, S10). The agent reuses a stable canonical vocabulary while rarer elements vary from run to run. A core of 92 marker genes appears in every run, out of 462 seen at least once in the resource-unconstrained variant, a sharply bimodal recurrence spectrum that explains why the gene-Jaccard sits near 0.53. The same stable core holds at the module and axis levels, with a consistent axis-kind composition (plain, contrast, or gated), so most run-to-run disagreement is naming, not biology. Literature grounding adds a citation layer with the same stable core. A small set of anchor references (for example Segerstolpe 2016 (Segerstolpe et al., 2016) and the Tabula Sapiens atlas (The Tabula Sapiens Consortium, 2022)) recurs in nearly every run. This audit is possible only because the blueprint exposes named genes, modules, axes, and citations. Free-form scripts expose no such structured handles, so the structure is what turns run-to-run variation into something an expert can inspect and correct, and a foundation that further models can build on.

The approach is also not expensive to author, store, and run. A blueprint is a compact, literature-derived script, not a trained network, so authoring costs a single LLM session (Stage A: ∼30–125k tokens, 2–7 minutes) and yields a 10–40 KB script, and building the embedding is a CPU-only pass of minutes with no GPU (Supp. Fig. S11). The optional Stage B refinement loop is the dominant expense, and it substantially expands the specification’s literature coverage and citations while leaving predictive scIB quality unchanged within run-to-run s.d. in both grounding modes (Supp. Fig. S12), so it acts as audit and coverage completion rather than a quality knob and is best reserved for a final pass. Against conventional methods, the agentic pipelines sit in the low-cost, small-footprint end of the cost–quality trade-off (Supp. Fig. S11), reaching competitive overall scIB quality while storing a kilobyte-scale script rather than a 5 MB (scVI) to ∼5.7 GB (UCE) checkpoint and running on CPU in minutes rather than ∼12 (scVI) to ∼76 (UCE) minutes. This is not specific to the most capable engine, with GPT-5.4-mini authoring comparable blueprints at similar cost.

### Axis topology can be steered to encode an interpretable developmental tree

The blueprint’s axis algebra makes the embedding’s structure a controllable design target. Because each output dimension is an explicit, typed composition of named gene programs, the agent can be asked in natural language, through the scenario’s free-text notes, to arrange those dimensions into a chosen topology, the parent–child tree induced by the axes’ gate relations. We demonstrate this on the Pijuan-Sala mouse gastrulation atlas (mouse_gastrulation, E6.5–E8.5 (Pijuan-Sala et al., 2019)). A single instruction directs the agent to organize its axes, each labeled by a developmental cell state, as a time-forward gated tree, in which tree depth reflects developmental time and sibling states are separated by passing the sibling modules as negatives. The agent returns a 40-axis tree (five plain roots and 35 gated axes, citing 12 literature resources; the refined showcase blueprint, Supp. Fig. S13). Its topology recovers the canonical lineage hierarchy of the gastrulating embryo (Pijuan-Sala et al., 2019) (Figure 5a). The epiblast branches into primitive streak, neural and surface ectoderm, and primordial germ cells. The streak in turn gives rise to nascent mesoderm, definitive endoderm, and the axial node/notochord. Mesoderm splits into cardiac, haemato-endothelial, and paraxial fates, and the haemato-endothelial axis continues into a blood progenitor and onward into erythroid, myeloid, and megakaryocyte/EMP leaves. Gut tube endoderm divides into foregut and hindgut, and the neural tube into anterior and posterior neural states. This structure is designed in, not discovered post hoc, as trajectory-inference tools such as PAGA (Wolf et al., 2019) instead recover the branching lineage graph from transcriptional similarity in a data representation, which need not reflect the underlying developmental process (Tritschler et al., 2019).

**Figure 5.**
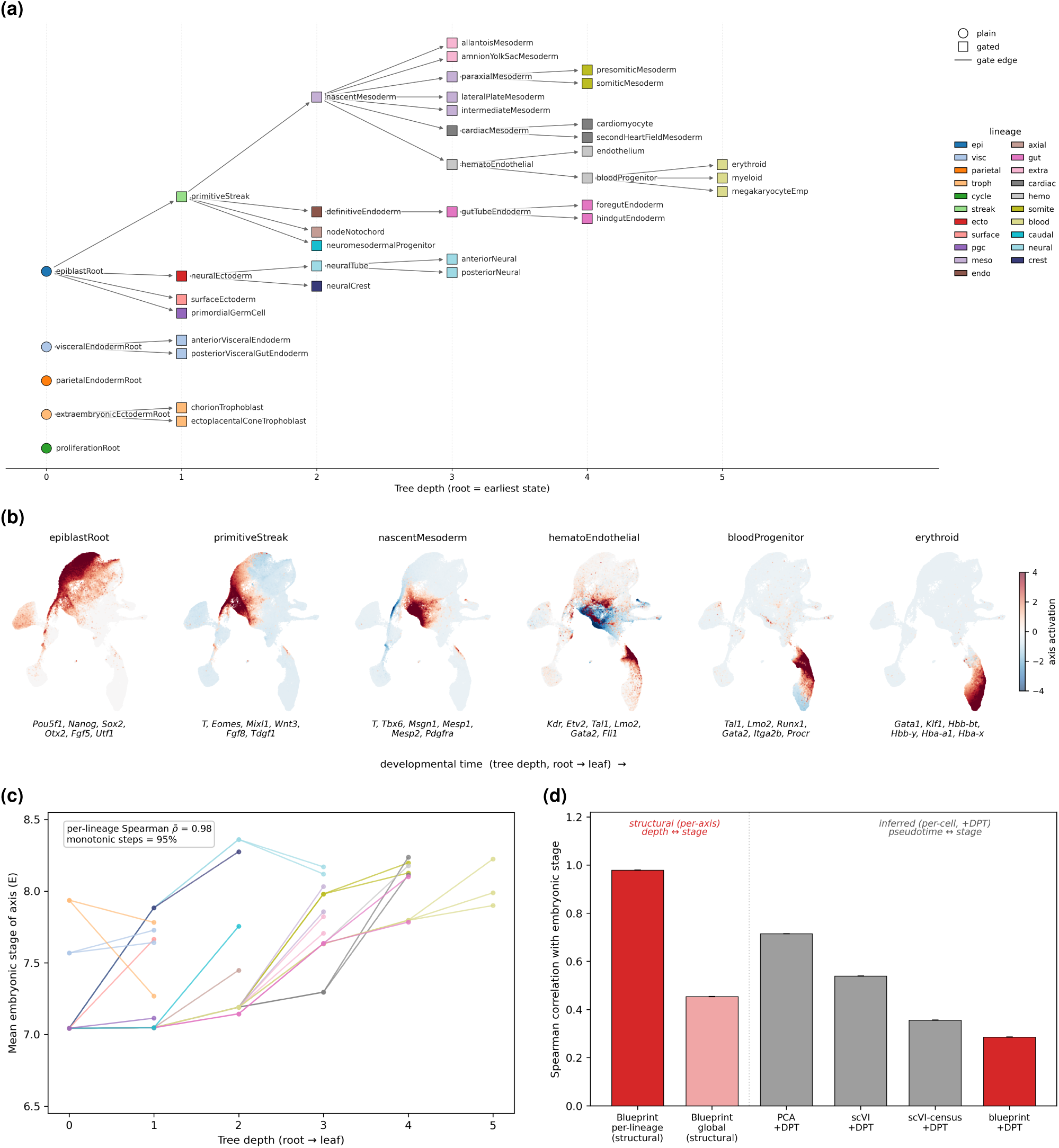
Axis topology steered into a time-ordered developmental tree. Refined blueprint for the mouse_gastrulation embryo atlas (E6.5–E8.5), authored with Codex, GPT-5.5 at high reasoning effort. (**a**) The axis tree of the authored blueprint. Nodes are embedding axes, the *x* position is tree depth (root to leaf), edges are parent-to-child gate relations, color encodes lineage, and marker shape encodes axis kind (plain root or gated). (**b**) One root-to-leaf branch, ordered left to right by increasing tree depth. Each panel is one shared UMAP of the embedding, colored by that axis’s activation. Beneath each panel are the first six marker genes of the axis’s positive gene module, which the agent curated from the literature. (**c**) For each root-to-leaf lineage (one line), the mean annotated embryonic stage of the cells assigned to each axis against tree depth. The per-lineage Spearman correlation and the percentage of non-decreasing parent-to-child steps are shown at the top left. (**d**) Absolute Spearman correlation with annotated embryonic stage. The structural bars are computed per axis, relating each axis’s tree depth to the mean stage of its assigned cells, averaged within each lineage (per-lineage) or pooled over all axes (global). The inferred “+DPT” bars are computed per cell, relating each cell’s diffusion pseudotime (DPT (Haghverdi et al., 2016), rooted at an Epiblast/E6.5 cell) to its stage (*n* ≈ 109k cells with an annotated stage). The two groups are not a head-to-head comparison. They measure different quantities, a by-construction ordering of named states against an inferred per-cell pseudotime.

The encoded depth reflects real developmental time rather than an arbitrary drawing. We assign each cell to the axis it scores most strongly and read off the mean annotated embryonic time of the cells assigned to it (Methods). Within each lineage of the tree, axis depth orders embryonic time almost perfectly (per-lineage Spearman *ρ* = 0.98, with 95% of parent-to-child steps non-decreasing in time, Figure 5c), and this ordering reproduces across three independent runs (*ρ* = 0.89–1.00, Supp. Fig. S14). The ordering is, however, per-lineage. Pooled across all lineages the single global depth-versus-time correlation is only moderate (*ρ* ≈ 0.45), because distinct lineages mature at different absolute rates, so depth indexes developmental progression within each lineage rather than against a single absolute time scale common to all lineages. The branch points recover known lineage splits, with each named axis dominated by its matching annotated population. The progression is directly visible in a two-dimensional UMAP of the embedding. Walking a single branch from the epiblast root down to the erythroid leaf, the axis activation sweeps continuously across the embedding as depth increases (Figure 5b), and three further lineages spanning endoderm, mesoderm, and ectoderm show a similar continuous sweep (Supp. Fig. S15). The embedding as a whole lays cells out along the E6.5→E8.5 developmental gradient (Supp. Fig. S16).

The temporal ordering is carried by the axis topology, not by the geometry of the point cloud. Each axis is still an expression-similarity readout, the mean expression of its marker genes, so the embedding is grounded in gene expression just as conventional methods are. The two encodings are thus complementary: the agent arranges those similarity axes into a tree whose depth is developmental time. A conventional embedding instead carries time only implicitly, recoverable through a separate trajectory-inference step whose similarity-based ordering must be validated against an external time label rather than taken as ground truth (Tritschler et al., 2019). We therefore score every pseudotime against the annotated embryonic stage. Running diffusion pseudotime (DPT (Haghverdi et al., 2016)) on each embedding’s geometry, a global pseudotime recovers embryonic stage moderately for PCA (*ρ* = 0.72), scVI (*ρ* = 0.54), and scVI-census (*ρ* = 0.36), but only after a root is chosen and only as a single unnamed scalar.

The blueprint is by design a weak diffusion substrate (*ρ* ≈ 0.29, Figure 5d). It thus supplies by construction what a black-box embedding yields only after extra inference and without interpretability, a named, literature-grounded, time-ordered lineage tree that a domain expert can read, audit, and edit. The same branches can be traversed interactively in the web application released with the code, which recolors the embedding along each root-to-leaf path.

## Discussion

We have shown that a coding agent can author an interpretable single-cell embedding model directly from the literature, given only a description of the biological scenario and with no gene-set database, no training, and without ever seeing the data. Its named axes faithfully discriminate the cell types they name, and the zero-shot embeddings are competitive in biological quality with conventional, foundation-model, and program-informed baselines while remaining batch-robust by construction. Authoring is reproducible across runs, produced in minutes at kilobyte scale, and auditable down to individual genes, programs, and citations, and a single natural-language instruction can steer the embedding’s topology into a structured form such as a developmental tree. No existing approach delivers these properties together. Data-driven embeddings need post-hoc interpretation (Lopez et al., 2018; Lotfollahi et al., 2023) and a dedicated batch-correction step (Luecken et al., 2022) that the blueprint does not. Interpretable program methods are readable too, but draw their programs from a fixed database (Andreatta and Carmona, 2021; Lotfollahi et al., 2023) rather than curating them for the scenario, and offer no way to compose them into typed axes. Zero-shot foundation models, despite broad pretraining, did not outperform simpler baselines in our benchmark (Kedzierska et al., 2025), while the blueprint stays competitive without it. We also deliver the approach as a usable tool, an interactive web application that runs locally so a practitioner can author, edit, and apply a blueprint to their own data without writing code.

Two of the blueprint’s properties, interpretability and batch robustness, follow from a single design choice. Each axis reads out only genes the literature validates as markers of a cell state (Pasquini et al., 2021; Hu et al., 2023). Reading a dimension therefore requires nothing beyond its declared gene set, and the genes and library-size shifts that carry batch and protocol effects have little dimension to load onto. The embedding is interpretable and batch-invariant at once, with no post-hoc attribution step and without ever seeing a batch label. This is a different route to batch robustness than correction-based integration takes. scVI and expiMap reach the batch axis by conditioning a trained latent on the batch key (Lopez et al., 2018; Lotfollahi et al., 2023), whereas the blueprint and UCell, used here as an embedding model (Andreatta and Carmona, 2021), reach it by construction, simply by leaving no channel in which a technical signal can be encoded. The same marker-only design explains where the agent pulls ahead of its fixed-database comparators. When the relevant programs are absent from a database, as cortical subclasses are from PanglaoDB (Franzén et al., 2019) and Reactome (Gillespie et al., 2022), gene-set scoring cannot recover them, whereas the agent curates roughly 23 subclass axes. We read this as evidence that, in the shown examples, the dynamically curated, literature-grounded programs are the decisive factor, not the scoring function itself.

Because the blueprint is an explicit, editable specification rather than a trained black box, the biology it encodes can be steered in ways a learned embedding does not afford. Adding or removing a named program changes the embedding by construction. As one example, the showcase cortex model did not give the rare Sst Chodl interneuron its own axis, an explicit gap that surfaces as a cell type with no named axis (Figure 1d). A user could close it by appending a gated axis that isolates this subtype through its defining marker *Chodl* within the Sst class (Tasic et al., 2018). The embedding would then gain a dedicated, named Sst Chodl axis by construction, a targeted edit that a black-box embedding does not afford. The same control extends from content to axis topology. A single natural-language instruction arranged the axes of the gastrulation blueprint into a time-forward tree whose depth tracked embryonic stage within each lineage, a structure designed in rather than recovered post hoc from transcriptional similarity (Tritschler et al., 2019). Because each axis is itself a named developmental state at a known tree depth, assigning each cell to its most-activated axis also yields a marker-grounded state label that carries a temporal rank, a coarse per-cell staging obtained without any trajectory inference.

Several limitations bound these claims. First, our benchmark datasets are published and open access, so the agent can draw on the very studies that describe them, together with their annotated types and markers, whether by retrieving them while authoring or from knowledge already in its pretraining. The curated programs are therefore not strictly independent of the benchmark’s source, even though the agent never sees the expression data or the labels and the scenario does not name the dataset. Our quality comparisons are thus best read as evidence that the approach translates known biology into a faithful embedding rather than as a blind test of generalization to unseen biology. Second, the approach is purely prior-driven, so it captures only what the literature already describes. It therefore cannot represent cell states the literature characterizes poorly, correct mismatches between curated markers and a given dataset, or discover structure the literature has not named, whereas a data-driven method retains these abilities by learning de novo (Lopez et al., 2018). The approach is therefore expected to be strongest in well-characterized domains, where the literature richly describes the relevant cell states. Where that coverage is sparse, we expect the blueprint to serve as a refinable prior rather than a standalone embedding. Third, the authored programs are not guaranteed correct, because the agent’s knowledge comes from language-model pretraining over heterogeneous, variable-quality text. The agent can curate inaccurate markers and individual axes can fail, such as the below-chance border-macrophage axis in our cortex showcase (Figure 1d), but the optional Stage B loop, expert review, the enforced literature grounding, and the citation audit let a user correct and verify which sources enter each program. Fourth, we established reproducibility, auditability, and cost in depth on only one representative dataset (the human pancreas) and one coding-agent harness, the Codex command-line interface driving OpenAI GPT models. The GPT-5.4-mini engine ablation indicates this does not hinge on the most capable engine, and confirming generality across tissues, biological conditions, and agentic frameworks remains future work. Finally, because authoring depends on an external language model whose weights may change, we treat model-version drift as a reproducibility limitation and mitigate it by pinning model identifiers and archiving every authored blueprint. Two directions follow naturally. The first is to combine the literature and data-driven lenses, which is also how we expect the approach to reach biology the literature describes only sparsely. Because they are complementary, capturing different and partly non-overlapping aspects of the same biology, combining them is more promising than choosing between them. Coupling the blueprint to a lightweight learned model that fits only the residual variation the blueprint leaves unexplained, in the spirit of masked-decoder approaches (Lotfollahi et al., 2023) or sparse-encoder approaches such as the boosting autoencoder (Hackenberg et al., 2025), could turn the auditable prior into a starting point for discovering structure the literature does not yet describe. A domain expert or an agent could then judge which learned residual patterns reflect genuine biology rather than technical noise and fold the validated ones back into the blueprint. This keeps the combined model interpretable and auditable and lets it surface populations the literature has not yet named. The second direction is local, dependency-free authoring. Running the agent on open, on-device language models would let the authoring step also run on the user’s machine, removing the one remaining external dependency. More broadly, our results suggest that coding agents can convert the scattered, qualitative knowledge of a field into quantitative, auditable models, provided their output is disciplined by structure. Constraining the agent to populate a declarative template through a fixed API, rather than to write arbitrary code, is what confines run-to-run variation to biological content and keeps every model standardized, auditable, and composable, a design principle we expect to transfer beyond single-cell genomics.

## Methods

### Overview of the agentic blueprint framework

A coding agent authors an embedding model from three inputs, without ever being shown single-cell RNA-sequencing (scRNA-seq) data and without training any computational model. The first, the only input the user provides, is a scenario configuration, a JSON file with comments that declares the biological context through a set of fields (species, tissue, celltype, age, sex, condition, and free-text notes). Only species is required, and null marks an unspecified field that is never filled with a default (Supp. Code S1). The free-text notes field is an optional design directive the user can supply. It steers the agent’s emphasis or the embedding’s structure, for example to resolve specific subtypes or to arrange the axes into a developmental tree. The second is a fixed Python template (the embedder_blueprint.py module) that exposes the embedding application programming interface (API) described below (Supp. Code S3). Its only agent-editable region is the build_blueprint() function. The third is a fixed prompt template that unifies the task, the scenario, and the literature-grounding rules (Supp. Code S2). We drive the agent with the codex command-line interface in non-interactive mode (codex exec), with web search and code execution enabled throughout. Unless stated otherwise, we run GPT-5.5 at high reasoning effort. For the reproducibility and cost analyses on human_pancreas we additionally author the literature-grounded and resource-unconstrained cohorts with a smaller, lower-cost model (GPT-5.4-mini at high and medium reasoning effort) as an engine ablation. The full pipeline, from these inputs to a comparable embedding, is laid out in Supp. Fig. S1.

Authoring proceeds in one mandatory stage and one optional stage. In the first stage (Stage A), the agent reasons over its pretrained knowledge and the literature it retrieves, edits build_blueprint() to instantiate a set of named, literature-cited gene programs and embedding axes (Supp. Code S4), and self-verifies the result (below). The authored specification (the *blueprint*) is a short Python script rather than a trained model. Constructing the embedding for a dataset is then a deterministic, CPU-only pass over the count matrix (next subsection).

An optional second stage (Stage B) refinement loop iteratively critiques and revises the Stage A blueprint. A *discriminator* agent receives the scenario and the blueprint’s structural summary (its --describe output, without the citation information). It rates the blueprint’s biological coverage and plausibility as fine, minor, or substantial, and returns a structured list of findings (Supp. Code S5). If the rating is not fine, a *generator* agent edits the blueprint to address each finding (Supp. Code S6). A regression guard then compares the element sets before and after the edit and reverts any deletion of a resource, gene module, or axis that the discriminator did not explicitly authorize. The loop runs for at most a pre-specified number of iterations, three in all runs reported here. Each iteration runs in isolation. The harness hides the prior reports, snapshots, and transcripts, so every judgment is made afresh. The Stage A blueprint is archived before refinement. Each refined run therefore also yields its unrefined snapshot as a separate cohort, which enables paired Stage A versus Stage A+B comparisons without re-running the pipeline. Under the resource-unconstrained variant (below) the discriminator and generator are replaced by citation-free counterparts.

### Blueprint and embedding construction

The blueprint API comprises three core structural components. A *resource* is a citable literature source (add_resource) of kind paper, atlas, or database. A *gene module* (add_gene_module) is a named set of gene symbols citing one or more resources by key. An *axis* (add_axis, add_contrast_axis, add_gated_axis) is one output dimension built from modules. The number of registered axes sets the embedding dimensionality, which the agent is instructed to let the biology determine rather than fix *a priori*. Axes are written in declaration order, which fixes the order of the output dimensions.

Data to be embedded must be provided as an AnnData object (an .h5ad file) whose expression matrix holds log-normalized counts. For a cell, each module *m* is scored as the mean expression *s_m_* of its member genes present in this matrix. Genes are matched case-insensitively as whole tokens, without alias resolution. A module with no present genes scores zero. Axes then combine these module scores with fixed algebra, where *s_p_*, *s_n_*, and *s_g_* are the scores of an axis’s positive, negative, and gate modules and *a* is the axis activation. A *plain* axis takes its positive module’s score, *a* = *s_p_*, reading out a single program such as a cell-type identity. A *contrast* axis subtracts a negative module from the positive, *a* = *s_p_* − *ws_n_*, with weight *w* = 1, separating two opposing populations along one axis. A *gated* axis multiplies a contrast by its gate module, 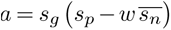, where 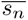 is the mean over one or more negative modules and the default weight is *w* = 0.5 (with no negatives, *a* = *s_g_ s_p_*). The gate restricts a program to a parent class and the negatives separate sibling subtypes, so a gated axis isolates a specific subtype within a broad class, and chained gates encode a hierarchy such as a developmental tree. Each axis is finally standardized across cells by column-wise *z*-scoring and clipped symmetrically to [−4, 4] (a configurable threshold). A constant axis maps to a zero column. A command-line minimum-gene threshold that can drop axes with too few present genes is left at its default of zero, so no axis is dropped. The result is written as a Parquet table with a cell_id column and dimensions dim_0. . . dim_{N-1} in axis order, alongside a JSON file recording, per module, how many declared genes were found in the dataset, which were missing, and the fraction of cells with a non-zero score. The template that implements this API and scoring engine is reproduced in Supp. Code S3, and the blueprint the agent authored for the mouse_cortex showcase in Supp. Code S4.

### Literature grounding and verification

Under the literature-grounded variant, each gene module must cite at least one resource. Papers must be peer-reviewed, carry a journal impact factor above 10, and have a digital object identifier (DOI). The impact-factor cutoff is a coarse heuristic to focus the agent on well-established findings in widely read journals, not a strict quality criterion. Reference atlases are cited through their companion paper and likewise carry a DOI. Curated pathway or ontology databases (KEGG, MSigDB, Reactome, GO, WikiPathways) may omit the DOI, because the gene-set identifier in the citation already identifies the gene set unambiguously. These rules are stated to the agent in the edit prompt (Supp. Code S2).

The agent edits only build_blueprint() and may not add helpers, classes, or imports. All API arguments are inline string and list literals, so the file remains statically parseable. The agent is also forbidden from training models or running principal component analysis (PCA), uniform manifold approximation and projection (UMAP), or pretrained-model inference, so the embedding derives solely from the declared gene programs. The scaffold enforces the machine-checkable part of this contract at run time. The API validators reject malformed DOIs (checked against a Crossref-style pattern), unknown resource kinds, duplicate keys, and any module that cites no resource. A placeholder check additionally rejects any skeleton text the agent left unfilled.

Impact factor and DOI authenticity cannot be verified from the file alone, so they are enforced instead by the prompt and by the agent’s own self-verification. After authoring, the agent self-verifies in two passes. A structural pass introspects the blueprint (python embedder_blueprint.py --describe), and a batched Crossref query confirms the title and first author of every paper and atlas DOI. A single consolidated correction pass then fixes any mismatch. We study two grounding variants that differ only in whether citations are required. The *literature-grounded* variant requires every module to cite resources. The *resource-unconstrained* variant (--noref) makes citations optional, a softer control on the role of forced literature grounding. The biology is still literature-derived, but the agent is not required to cite it.

### Free-form prompting baselines

To isolate the contribution of the structured blueprint, we compare against *free-form* prompting, in which the agent writes an arbitrary Python script (embed_freeform.py) that reads the same AnnData input and emits the same cell_id+dim_* Parquet output, with no blueprint API. Three variants match the blueprint’s grounding variants at the level of content rules. A *naïve* variant has no grounding rules, a *literature-grounded* variant inherits the impact-factor and DOI rules, and a *resource-unconstrained* variant drops them (Supp. Code S7–S9). Because free-form output is unconstrained, we additionally record its failure modes. We log scripts that fail to execute, scripts that violate the output contract (for example a malformed schema), the recovered gene tokens intersected with the cohort vocabulary, and the citation density. None of the 30 free-form runs (the three variants over ten seeds each) hard-failed, and all produced schema-valid output.

### Datasets and preprocessing

We use four datasets spanning mouse and human tissues (Table 1). They are an integrated mouse isocortex atlas (mouse_cortex), the human pancreas compendium (human_pancreas), the human immune compendium (human_immune), and the Pijuan-Sala mouse gastrulation atlas (mouse_gastrulation, E6.5–E8.5 (Pijuan-Sala et al., 2019)). The first three form the quantitative benchmark. The pancreas and immune compendia are the pancreas and “Immune (human)” integration tasks of the single-cell integration benchmarking (scIB) study (Luecken et al., 2022). We establish the process-level reproducibility, auditability, and cost analyses in depth on human_pancreas (Baron et al., 2016; Grün et al., 2016; Muraro et al., 2016; Segerstolpe et al., 2016; Xin et al., 2016; Lawlor et al., 2017). The gastrulation atlas is reserved for the structured developmental-tree showcase.

**Table 1.**
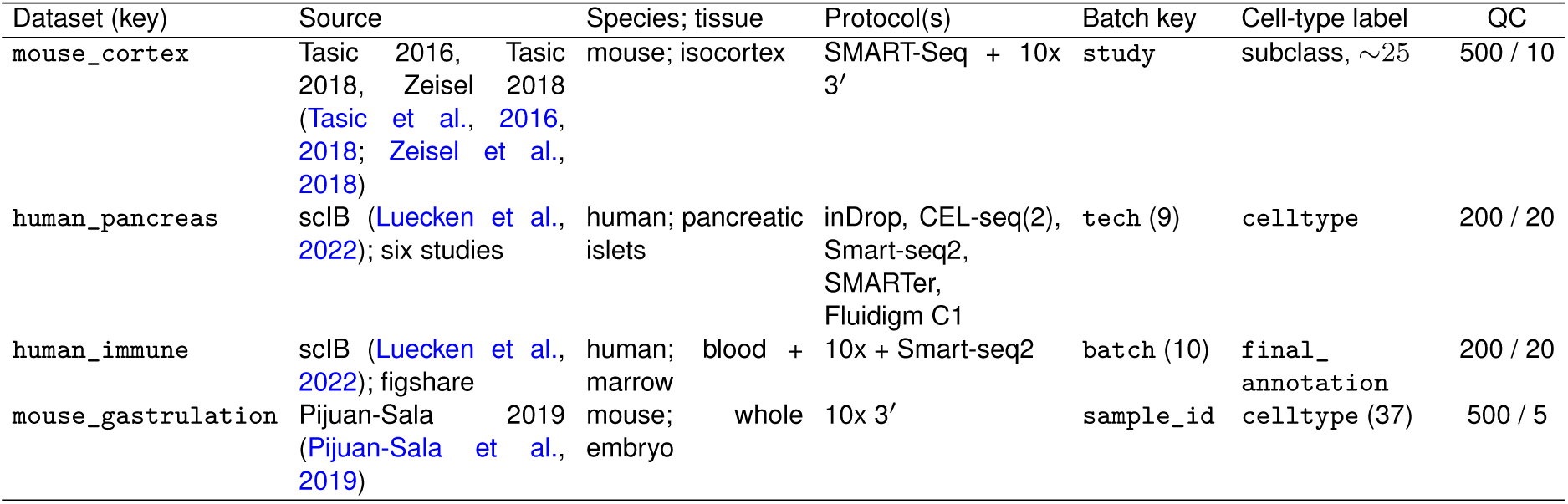
Datasets and preprocessing. The QC column gives quality-control thresholds as minimum detected genes per cell / maximum percent mitochondrial counts. Genes detected in fewer than three cells are removed in every dataset. The six human pancreas studies are Baron 2016, Grün 2016, Muraro 2016, Segerstolpe 2016, Xin 2016, and Lawlor 2017 (Baron et al., 2016; Grün et al., 2016; Muraro et al., 2016; Segerstolpe et al., 2016; Xin et al., 2016; Lawlor et al., 2017). The mouse_cortex cell-type labels are a ∼25-type subclass vocabulary we curated, harmonizing the three studies onto the Tasic 2018 scheme (Methods).

All datasets are processed identically up to per-dataset thresholds, with raw counts retained throughout. In a quality control step (scanpy (Wolf et al., 2018)), we remove cells with too few detected genes or too high a mitochondrial fraction, and genes detected in fewer than three cells. Thresholds are looser for droplet and multi-protocol data and stricter for the embryonic mouse_gastrulation (Table 1). We then library-size normalize to 10^4^ counts per cell, log(1 + *x*)-transform, and flag the top 2,000 highly variable genes (Seurat v3 flavor) ranked within the dataset’s batch covariate, without subsetting the gene set. Raw, normalized, and log-normalized layers are all preserved at this stage.

Two of the benchmark datasets are deliberately heterogeneous. mouse_cortex concatenates three studies spanning full-length SMART-Seq and 3*^′^* droplet chemistries (Tasic et al., 2016, 2018; Zeisel et al., 2018) on overlapping cortical biology, so that cross-protocol integratability can be assessed without explicit batch correction.

The three studies do not share a common annotation, so we curated the cell-type label ourselves. We mapped each study’s native annotations onto a common ∼25-type subclass vocabulary that we defined, using manually authored per-study mapping tables. The vocabulary is anchored on the Tasic 2018 cortical subclass scheme. We retained and audited any native label that falls outside it. A coarse three-class grouping (glutamatergic, GABAergic, non-neuronal) derived from it provides the root level for the gating-tree showcase. To keep the evaluation ground truth clean, we remove at integration any cell carrying no usable harmonized label (unannotated cells and the droplet atlas’s generic “Neurons” call, which resolves to no subclass). mouse_gastrulation carries, beyond its 37 author-annotated cell types, a developmental-stage label (E6.5–E8.5) and a per-embryo batch (sample_id). The stage label anchors the temporal validation of the gating-tree showcase.

### Conventional and foundation-model baselines

We compare against established embedding methods, tiered by the prior knowledge they access, all producing the same cell_id+dim_* embedding Parquet so the downstream evaluation is identical. The unsupervised tier comprises principal component analysis (PCA, scikit-learn, 50 components on the highly variable genes, deterministic) and scVI (scvi-tools (Lopez et al., 2018), a variational autoencoder with a 10-dimensional latent space trained *de novo* on raw counts with the batch covariate as a conditioning input, library defaults otherwise). As a prior-free control we also run scVI without its batch key, which removes the batch conditioning. A foundation-model tier comprises scVI Census and the Universal Cell Embedding (UCE). scVI Census is the scVI model pretrained on the CZ CELLxGENE Census (version 2025-11-08 (Abdulla et al., 2025)), applied by mapping input gene symbols to the Census Ensembl reference and reading out the 50-dimensional embedding. Its human and mouse checkpoints were pretrained on roughly 94.5 and 18.3 million cells, respectively. UCE is the 33-layer, 1,280-dimensional checkpoint (Rosen et al., 2026), applied as a frozen forward pass. Stochastic methods are run across 10 random seeds. Deterministic methods (PCA, scVI Census, UCE) use a fixed seed and are scored once.

### Gene-set and interpretable-program baselines

We additionally compare against two methods that, like the blueprint, build their representation from explicit, named gene programs but require a human to supply those programs. Both emit the same cell_id+dim_* embedding Parquet, and we run each on gene sets sourced off the shelf, reflecting a database a practitioner would reach for rather than a hand-tuned panel. Cell-type marker sets are drawn from PanglaoDB (Franzén et al., 2019), taking every cell type it lists for the dataset’s organs (Brain for mouse_cortex, Pancreas for human_pancreas, immune system plus blood for human_immune) with no hand-mapping to the dataset’s annotation. Human symbols are used as listed, title-cased to the mouse convention for mouse_cortex.

UCell (Andreatta and Carmona, 2021) (a training-free, gene-set-scoring baseline, run via the Python port pyUCell (Andreatta and Carmona, 2026)) scores each cell against every signature with the within-cell Mann–Whitney rank statistic (maximum rank at the package default of 1,500). We take the resulting cell-by-signature matrix as the embedding, one named axis per signature, with constant signatures dropped. Scoring is deterministic and rank-based, so it is invariant to normalization and, like the blueprint, reads cells only off marker expression with no channel for batch. expiMap (Lotfollahi et al., 2023) (a trained, interpretable-program baseline) is a masked-decoder variational autoencoder (scArches (Lotfollahi et al., 2022)) that corrects batch by conditioning, as scVI does. A binary gene-by-program membership matrix masks its linear decoder so each latent node reconstructs only the genes of its program, fixing the named axes before training. Mirroring the original expiMap, the program collection combines

Reactome pathways (Gillespie et al., 2022) (the MSigDB (Liberzon et al., 2015) c2.cp.reactome collection for human and m2.cp.reactome for mouse) with PanglaoDB cell-type markers (above). We train the model on the 2,000 highly variable genes, keeping only programs with at least twelve member genes among them (min_genes = 12) and letting a group-lasso penalty deactivate uninformative programs so the surviving active programs form the latent axes. Training runs up to 400 epochs with early stopping, reconstruction weight *α* = 0.7, Kullback–Leibler weight 0.5, hidden layers [256, 256, 256], and no unconstrained extension nodes, on raw counts with the dataset’s batch covariate as the conditioning key. We train it over 10 random seeds.

### Embedding evaluation metrics

Biological quality and batch robustness are quantified with the scIB metric suite (Luecken et al., 2022) using the scib-metrics Benchmarker (Virshup et al., 2023), a Python/JAX reimplementation that builds an approximate nearest-neighbor graph per embedding (pynndescent, *k* = 15). The clustering metrics, adjusted Rand index (ARI) and normalized mutual information (NMI), use Leiden clustering (Traag et al., 2019) with the resolution optimized against the reference labels (a sweep over ten resolutions, as in scIB), rather than the Benchmarker’s single-fixed-*k k*-means default. The bio-conservation sub-score is the unweighted mean of NMI, ARI, label-based average silhouette width (ASW-label), the cell-type local inverse Simpson’s index (cLISI (Korsunsky et al., 2019)), and the isolated-label score. The batch-correction sub-score is the unweighted mean of graph connectivity, batch-based silhouette width (ASW-batch), the integration local inverse Simpson’s index (iLISI (Korsunsky et al., 2019)), the *k*-nearest-neighbor batch test (kBET (Büttner et al., 2019)), and principal-component regression (PCR) of the batch covariate against an unintegrated PCA reference. Per scIB, graph connectivity is treated as a batch metric. Where an overall score is needed we use the scIB composite 0.6 · bio + 0.4 · batch (the package Total). We report the participation ratio of the embedding as a measure of effective dimensionality. All metrics are computed per cell-type label column from the AnnData object. The values are internally consistent across the compared methods but not directly comparable to numbers from the original scib package.

### Embedding visualization

For display, we project each embedding to two dimensions with UMAP (umap-learn), computed directly on the embedding’s dim_* columns with n_neighbors = 30, min_dist = 0.15, and a fixed random seed. In the per-method UMAP grids we show, for each method, the seed with the highest scIB composite score. Marker-gene dot plots show, for each gene, the per-cell-type mean log(1 + *x*) expression *z*-scored across cell types as the dot color and the fraction of cells in the type with non-zero expression as the dot size, with genes grouped by the axis that declares them. Per-axis activation maps recolor a single shared UMAP by each axis’s *z*-scored activation.

### Held-out reference-to-query transfer

To test generalization to data held out of authoring and training, we split each benchmark dataset into a reference and a query by withholding an entire study, technology, or donor as the query. The query is the full-length SMART-seq Tasic 2016 study for mouse_cortex, the Smart-seq2 technology for human_pancreas, and one bone-marrow donor (Oetjen_U) together with one peripheral-blood sample (Sun_sample4_TC) for human_immune. The trained comparators (scVI and expiMap) are fit on the reference alone, so the query is excluded from their training, and are then mapped onto the query either by a frozen forward pass or by a query-side scArches fine-tuning step (100 epochs) (Lotfollahi et al., 2022). For scVI we enable encode_covariates so that the frozen and fine-tuned mappings differ. The blueprint is applied zero-shot, scoring the reference and query jointly in a single pass. We score reference-to-query label transfer with a *k*-nearest-neighbor classifier (*k* = 15, distance-weighted) trained on the reference embedding and labels and predicting the query (accuracy and macro-*F*_1_, dropping query labels absent from the reference). We also score the joint reference-plus-query embedding with the same scIB suite used in the main benchmark, taking a joint highly-variable-gene PCA as the unintegrated reference. Each method variant is run over five seeds.

### Axis faithfulness evaluation

To assess whether each named axis discriminates the cell type it names, we compute, for every embedding axis and every annotated cell type, the one-vs-rest area under the receiver operating characteristic curve (AUROC) of the axis’s *z*-scored activation as a score for that type against all others (`scikit-learn`). Each axis’s named target type is taken from an axis-to-type mapping (`configs/axis_type_mappings/<dataset>/axis_types.csv, columns axis_name, cell_type`), which a coding agent (Claude Opus 4.8) drafted and we then refined by hand. Program or pan-class axes that name no single type, for example a pan-neuronal or immediate-early-activity program, carry a blank entry and are left unassigned. Across the axes that name a subclass we report the AUROC of each against its named type (mean, median, and the fraction above 0.80 and 0.90), and we mark the best axis per type as the per-type maximum over axes. As a complementary, threshold-free view we assign each cell to its single most-activated axis and tabulate, per annotated type, the fraction of its cells captured by each axis. To confirm the representative run is typical rather than favorably chosen, we recompute the per-type best-axis AUROC (the column-wise maximum over axes, with no axis-to-label matching) for every seed of the cohort. Cell-type labels are read from the AnnData object, and cells with no label are excluded.

### Reproducibility across re-authoring runs

We quantify reproducibility at three successive levels of the agentic pipeline. These levels ask how much the agent’s gene selection varies between runs, whether that variation leaves the representation stable, and whether the biology the embedding recovers stays consistent. At the blueprint specification level we measure the mean pairwise Jaccard overlap of the gene sets the coding agent selected across runs (canonical symbols). At the representation level we use linear centered kernel alignment (CKA (Kornblith et al., 2019)), a pairwise score between two runs’ mean-centered embeddings that is invariant to orthogonal rotation and isotropic scaling. For each cohort we report the off-diagonal mean of the seed-by-seed CKA matrix. Because each run enters many pairs, the pairwise CKA values are not independent, so we obtain the 95% confidence interval by drawing 1,000 bootstrap samples of runs rather than pairs. Finally, we report the consistency of biological quality as the mean and standard deviation (s.d.) of the per-seed scIB ARI across seeds, with the s.d. being the reproducibility signal.

### Recurrence of blueprint elements

To characterize what recurs in the agent-authored blueprints, we compute two summaries over the *N* runs of a model cohort. We count how many genes, modules, axes, and resources appear in exactly *k* of the *N* runs.

A pairwise Jaccard overlap measures how much any two runs share, reported as the off-diagonal mean *J* with a 1,000-resample bootstrap 95% confidence interval. Because free-text names vary even when they denote the same biology, we reconcile names in two ways. The first is a conservative, canonical form. We lowercase each name and strip all non-alphanumeric characters, so two names match only when their normalized strings are identical. This catches formatting differences but not synonyms. The second is a broader, concept-level criterion that treats synonymous names as equivalent by matching them through a controlled vocabulary. This is a per-dataset lookup table (`configs/name_mappings/<dataset>/{module_names,axis_names,resource_names}.csv`) that maps lexically distinct but synonymous raw names onto a shared standardized concept. A coding agent (Claude Opus 4.8) drafted this table, which we then refined by hand. Two items then match when their standardized concepts coincide. Names without a table entry fall back to the canonical form. Resources are identified by DOI where present and otherwise by their canonical key, so database resources participate alongside papers and atlases. Independent of naming, we summarize the axis-kind composition (plain, contrast, or gated) across the runs of a model cohort.

### Resource and cost accounting

The agent’s authoring cost is the language-model token usage and wall-clock time of Stage A, plus the Stage B refinement loop where it runs. We parse token usage from the agent transcripts and wall-clock from the orchestrator’s per-phase timing. Construction cost is the wall-clock to build the embedding. We time every method on CPU, excluding the time to read the input from disk. For each run we record the on-disk model footprint, which is the authored script for the agentic methods (tens of kilobytes) and the trained or pretrained checkpoint for the baselines, from a few megabytes for scVI to several gigabytes for UCE. We aggregate per-method costs across seeds (median and interquartile range) for the cost–quality comparison.

### Developmental-tree temporal evaluation

For the `mouse_gastrulation` showcase we test whether the agent-authored gate tree encodes developmental time, independent of any trajectory algorithm. We recover the tree from a run’s axis table, taking every non-root axis’s gating program as the edge to its parent, which gives each axis an integer depth (root = 0) and enumerates the root-to-leaf branches. We assign each cell to the axis of maximal *z*-scored activation and summarize each axis by the mean annotated embryonic stage (E6.5–E8.5, treated as numeric developmental time) of its assigned cells, dropping cells with no resolvable stage. Because tree depth is a property of an axis and not of a cell, we score how tree depth orders mean time at the axis level, with each axis contributing a single (depth, mean time) point. We report three statistics. The first is a per-lineage Spearman correlation between depth and mean time along each root-to-leaf branch, averaged over branches. The second is the fraction of parent-to-child steps along which the mean time is non-decreasing, a finer monotonicity check pooled over all steps. The third is a single Spearman rank correlation, pooled over the (depth, mean time) points of all axes. We also identify the dominant annotated cell type among each axis’s assigned cells, to check that the tree’s branch points match the expected lineage splits. We report each statistic for the refined showcase run, and as its mean and s.d. across the four re-authoring runs of the gastrulation blueprint (three independent Stage A runs plus the refined showcase).

To contrast the structural time with the pseudotime a trajectory algorithm extracts from an embedding’s geometry, we run identical diffusion pseudotime (DPT (Haghverdi et al., 2016), via scanpy (Wolf et al., 2018)) on every embedding. We build a *k*-nearest-neighbor graph (*k* = 15) on the embedding, compute a diffusion map (Haghverdi et al., 2015), root it at the medoid Epiblast cell at stage E6.5 under a rule identical for every method, and correlate the resulting per-cell pseudotime with the numeric embryonic stage (absolute Spearman). We apply the same procedure to the blueprint, PCA, scVI without its batch key, and scVI-census embeddings. We omit the batch-conditioned scVI here, because its batch covariate for this dataset is the per-embryo sample_id, which is confounded with developmental stage. For visualization, we recolor a UMAP of each embedding (computed as above) by individual axis activations along a chosen root-to-leaf branch, giving the branch filmstrips. The same per-axis recoloring is available interactively in the web application.

### Interactive web application

We provide an interactive web application (Dash) that runs the full workflow on a user’s own data without writing code. It runs locally in the browser, and the user’s expression data never leaves their machine. From a scenario description and the user’s own coding-agent credentials, it authors a blueprint through the same Stage A and optional Stage B pipeline. Authoring currently runs only through the codex command-line interface, so it requires the user’s own OpenAI credentials. The application then applies any authored blueprint to an uploaded .h5ad by the identical deterministic readout and presents the resulting embedding as an interactive UMAP. The uploaded .h5ad must hold log-normalized (log1p) counts in its expression matrix (adata.X). The UMAP can be recolored by axis activation, cell metadata, or individual gene expression, and, for a gated tree, swept along each root-to-leaf branch. Users can inspect, hand-edit, and re-apply the authored blueprint in the browser, and download the augmented AnnData and exported figures. The application vendors the codex pipeline driver so it is self-contained and keeps the user’s API key in memory only.

### Statistics, software, and reproducibility

Unless stated otherwise, each agentic cohort is authored over ten independent runs (seeds), and stochastic baselines over matched seeds, so that reported quantities carry the two relevant variance sources, stochastic agent authoring and stochastic model training. Among the four GPT-5.4-mini engine-ablation cohorts, one run failed during Stage A authoring and one during the embedding pass, so two cohorts retain nine runs and the other two retain all ten. Failed runs produce no embedding and are excluded from the analyses. We obtain 95% confidence intervals by bootstrap (1,000 resamples, with run-level resampling where pairwise quantities are aggregated), and report run-to-run dispersion of quality as mean ± s.d. across seeds. The clustering inside the metrics uses a fixed rando seed. We pin and report the agent model identifiers and access dates, the CZ CELLxGENE Census version, and package versions, and we archive every authored blueprint and scenario configuration in the project repository. All embedding construction and the conventional, foundation-model, and program-informed baselines were run on CPU on a MacBook Pro (Apple M2 Max, 12-core CPU, 96 GB RAM). Blueprint authoring itself (Stage A and Stage B) runs on the language-model provider’s infrastructure through the codex interface. Because the authoring step depends on an external language model whose weights may change over time, we treat model-version drift as an explicit reproducibility limitation and mitigate it through version pinning and artifact archival.

## Data availability

The datasets analyzed in this study are publicly available from their original sources, summarized in Table 1. The mouse_cortex atlas integrates three studies, obtained from the Gene Expression Omnibus (GEO) for Tasic 2016 (accession GSE71585) and Tasic 2018 (GSE115746), and from mousebrain.org (Linnarsson Lab, l5_all.loom) for Zeisel 2018. The human_pancreas and human_immune compendia are the preprocessed single-cell integration benchmark (scIB) datasets (Luecken et al., 2022), downloaded from figshare (files 24539828 and 25717328, respectively). The mouse_gastrulation data are the processed Pijuan-Sala 2019 embryo atlas (Pijuan-Sala et al., 2019), downloaded from the Marioni Lab (https://content.cruk.cam.ac.uk/jmlab/atlas_data.tar.gz). The scVI-census baseline uses the pretrained model from the CZ CELLxGENE Census, version 2025-11-08. All authored blueprints and analysis code are available at https://github.com/NiklasBrunn/blueprint-embedding. The processed AnnData objects are not deposited but can be regenerated from the public source datasets in Table 1 with the provided data pipeline in the code repository.

## Code availability

All code for the agentic authoring pipeline, the baseline embeddings, the evaluation suite, and the interactive web application is available at https://github.com/NiklasBrunn/blueprint-embedding under the MIT License. Each authored blueprint and scenario configuration is archived in the repository, and the agent model identifiers and access dates are recorded to support reproducibility.

## ACKNOWLEDGEMENTS

This work was funded by the Deutsche Forschungsgemeinschaft (DFG, German Research Foundation) under the Research Training Group MeInBio (GRK 2344, Project-ID 322977937; N.B., H.B.), the Collaborative Research Centre ‘Small Data’ (SFB 1597, Project-ID 499552394; N.B., S.M.K., H.B.), and Germany’s Excellence Strategy CIBSS – EXC-2189 – Project ID 390939984 (S.M.K., M.P., H.B.).

## AUTHOR CONTRIBUTIONS

N.B., S.M.K., and H.B. conceived and designed the study. N.B. developed the agentic blueprint framework, implemented the pipeline and web application, performed the experiments and analyses, and wrote the manuscript. S.M.K. contributed to the design of the agentic pipeline and the planning of experiments. M.Fri. contributed to the design of the agentic pipeline and to refining the web application. M.Fro. and M.P. contributed to the biological interpretation of the blueprints and results, and provided feedback on the manuscript. H.B. supervised the project, acquired funding, and edited the manuscript. All authors read and approved the final manuscript.

## COMPETING FINANCIAL INTERESTS

The authors declare no competing interests.

## Supplementary Information

### Agentic pipeline overview

**Figure S1.**
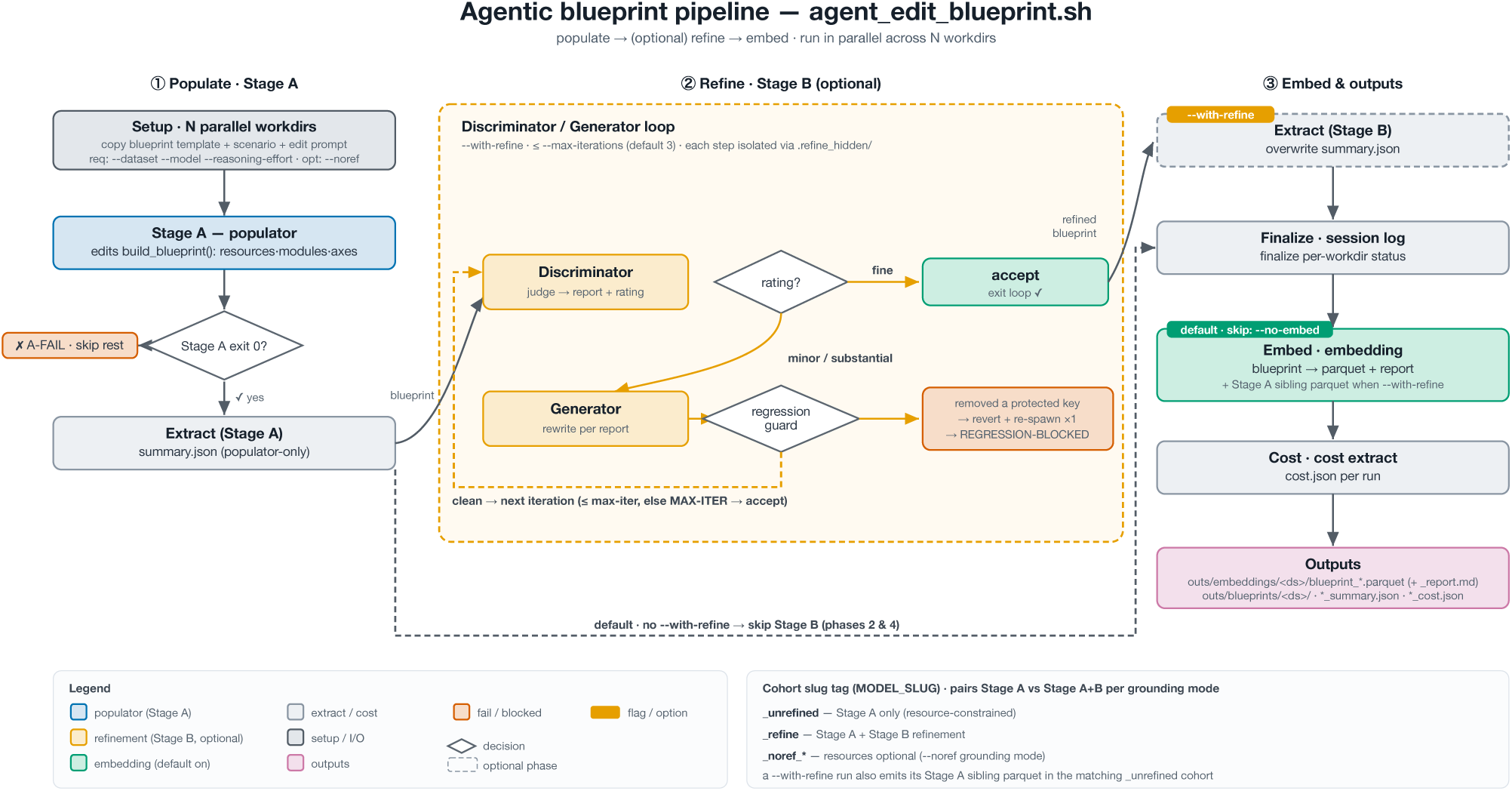
The agent-authored blueprint pipeline (agent_edit_blueprint.sh). A step-by-step expansion of the main-text model overview (Figure 1a,b), tracing one authoring run from its three inputs to a comparable embedding, with replicate runs executing in parallel across *N* workdirs. The flow has three stages, shown as the three columns. In *populate* (Stage A), a *Setup* step copies the embedder_blueprint.py template, the scenario configuration, and the edit prompt into each workdir, and the *populator* agent then edits only build_blueprint() to add cited resources, gene modules, and axes (plain, contrast, or gated), without ever seeing the expression data. A run whose Stage A exits non-zero (A-FAIL) skips the rest. The optional *refine* loop (Stage B, --with-refine) alternates a Discriminator that rates the blueprint ({fine, minor, substantial}) with a Generator that rewrites it to address the report, under a regression guard that reverts any deletion of a protected resource, gene module, or axis. A fine rating accepts and exits, and the loop runs at most --max-iterations (default 3), each iteration isolated from the previous reports. Without --with-refine this stage is skipped. The *embed* step (default, --no-embed skips) scores the authored blueprint on the preprocessed .h5ad into the cell-by-axis Parquet table with its per-module report, and a refined run additionally emits its Stage A snapshot as a sibling Parquet. Interleaved extract steps record a per-run summary, session status, and token and wall-clock cost as JSON, all written under outs/. The cohort slug tag (lower right) labels each run’s variant,_unrefined (Stage A only), _refine (Stage A+B), and _noref_* (the resource-optional --noref grounding mode).

### Supplementary Code: agent inputs and the authored blueprint

To make the authoring step fully inspectable, we reproduce in full the three inputs the coding agent received and the blueprint it returned, for the representative mouse_cortex showcase (literature-grounded, GPT-5.5 at high reasoning effort, seed 2, the run behind Figure 1). Supplementary Code S1 is the scenario configuration, Supp. Code S2 the blueprint-edit prompt template, and Supp. Code S3 the declarative Python template (embedder_blueprint.py). The agent’s only editable region is the build_blueprint() function at the bottom of that file. Supplementary Code S4 is the build_blueprint() the agent authored. The agent may edit only this function and changed nothing else, so the rest of the file is identical to the Python template and only the authored function is shown. The agent never sees the data. The embedding is the deterministic, CPU-only readout of this specification over the count matrix. The full scoring engine, all re-authoring seeds, and the other datasets and cohorts are in the manuscript’s code repository.

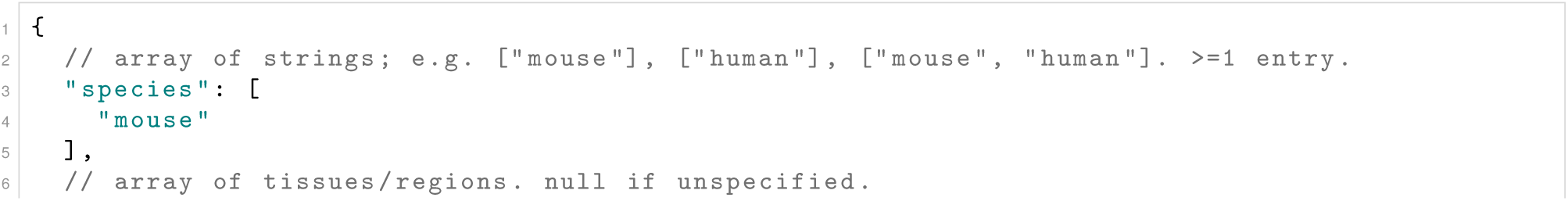
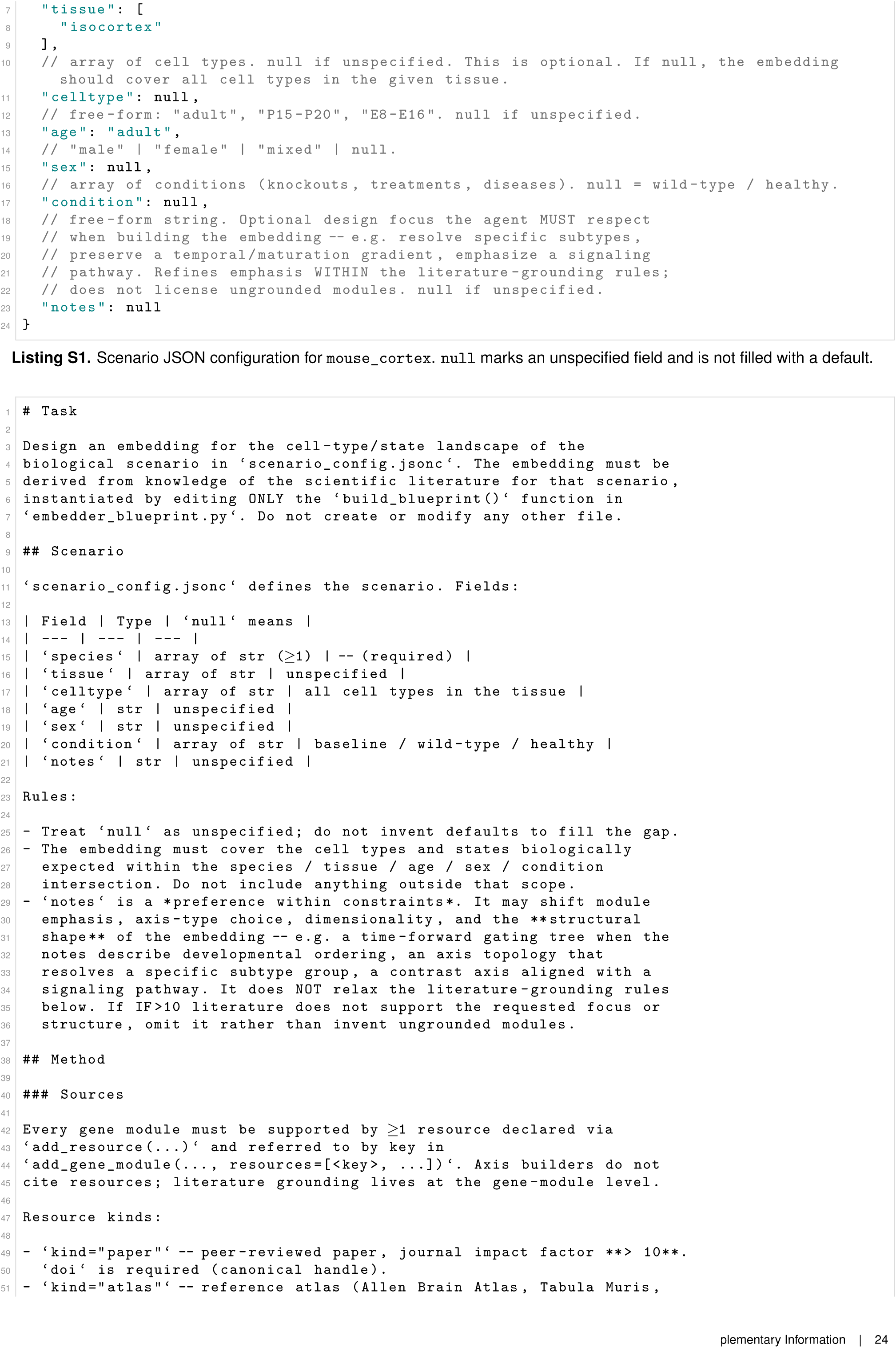
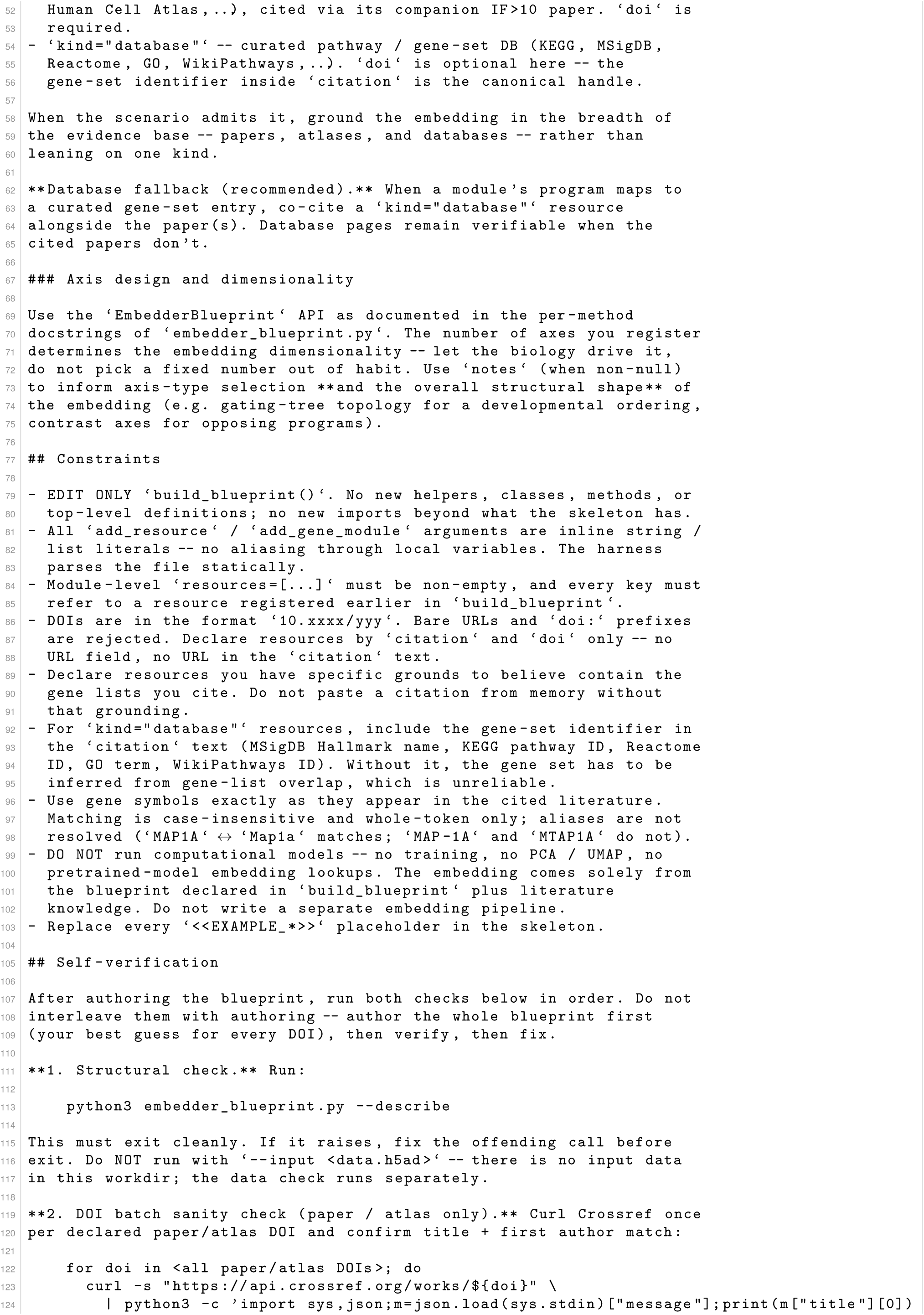
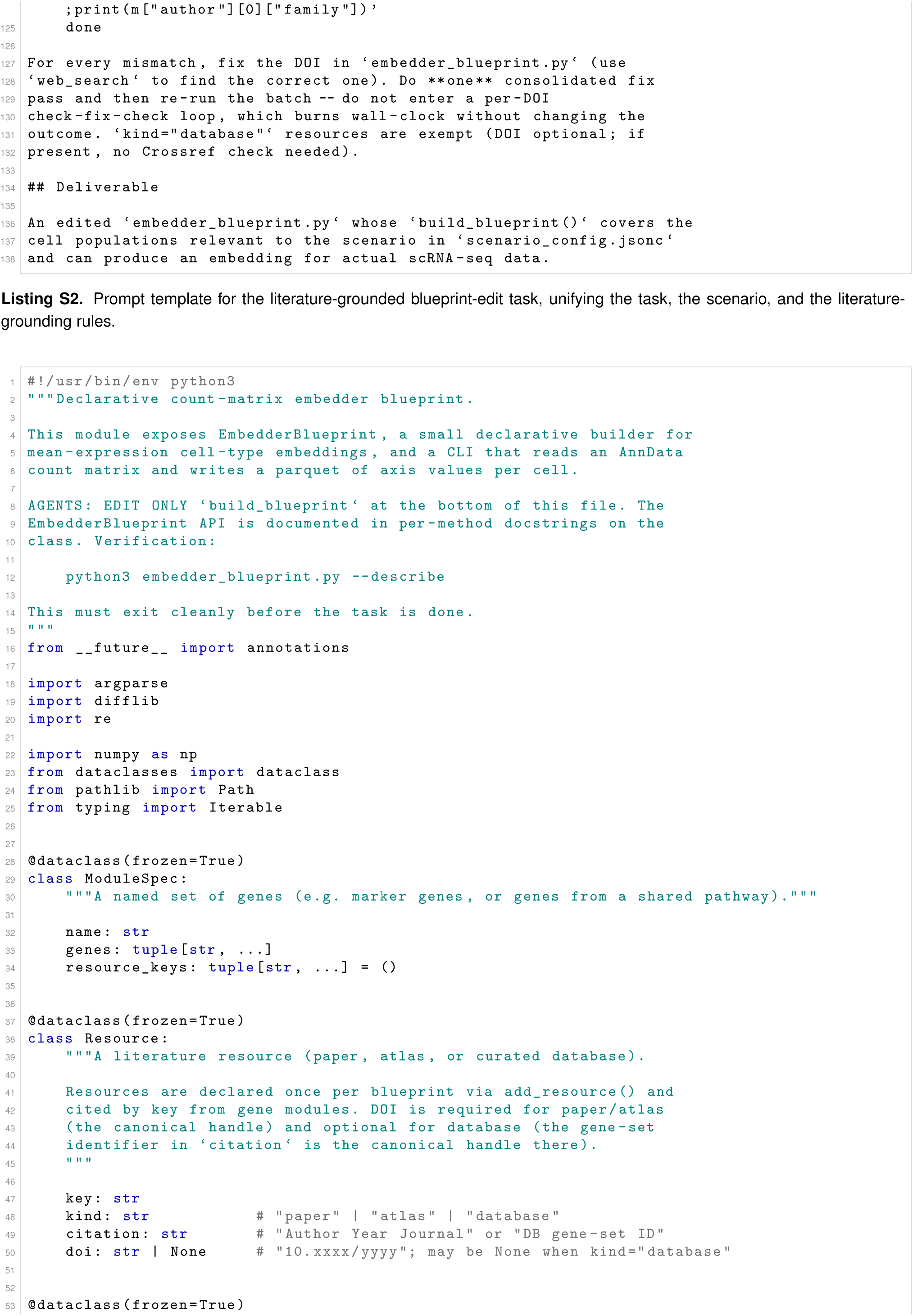
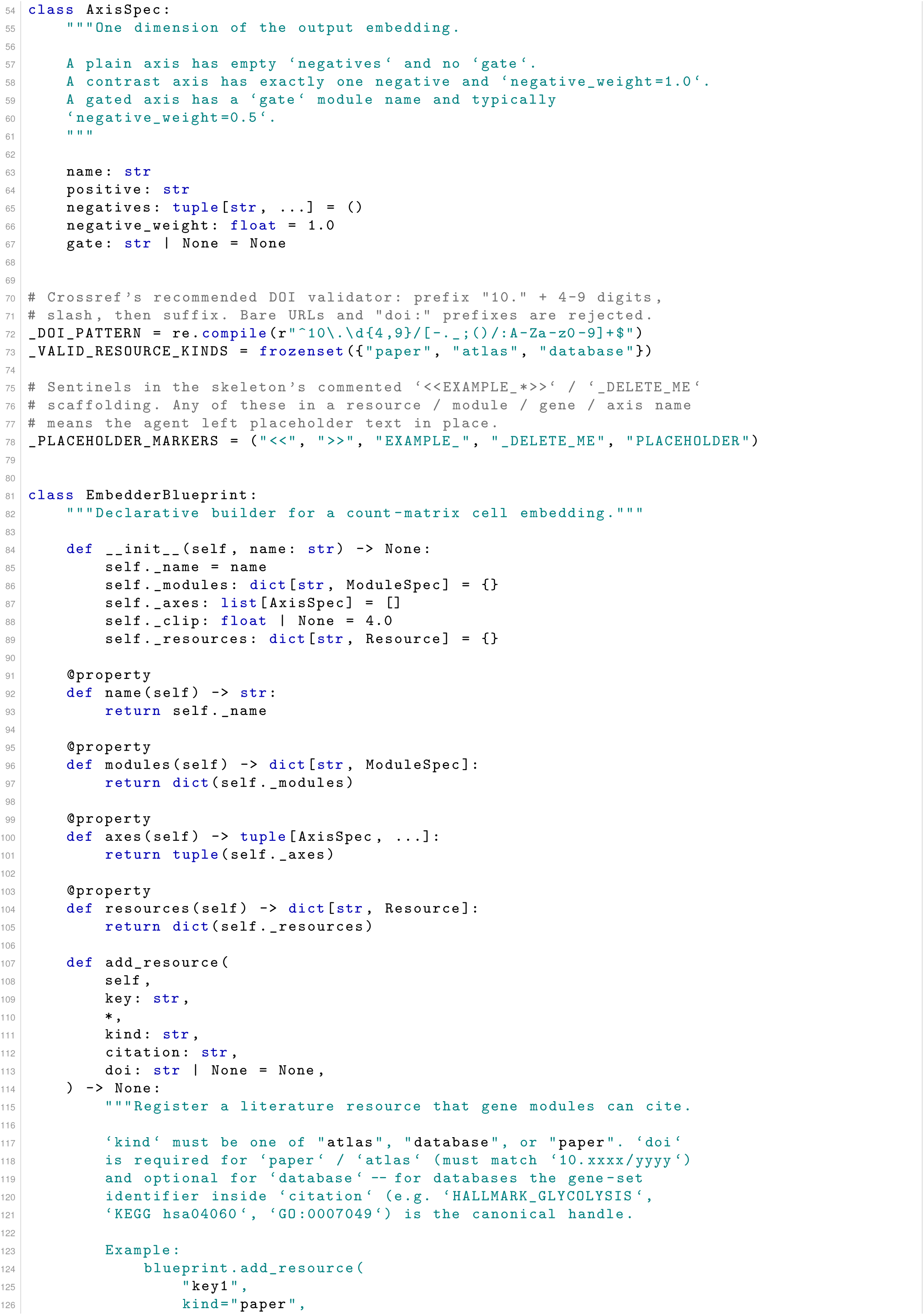
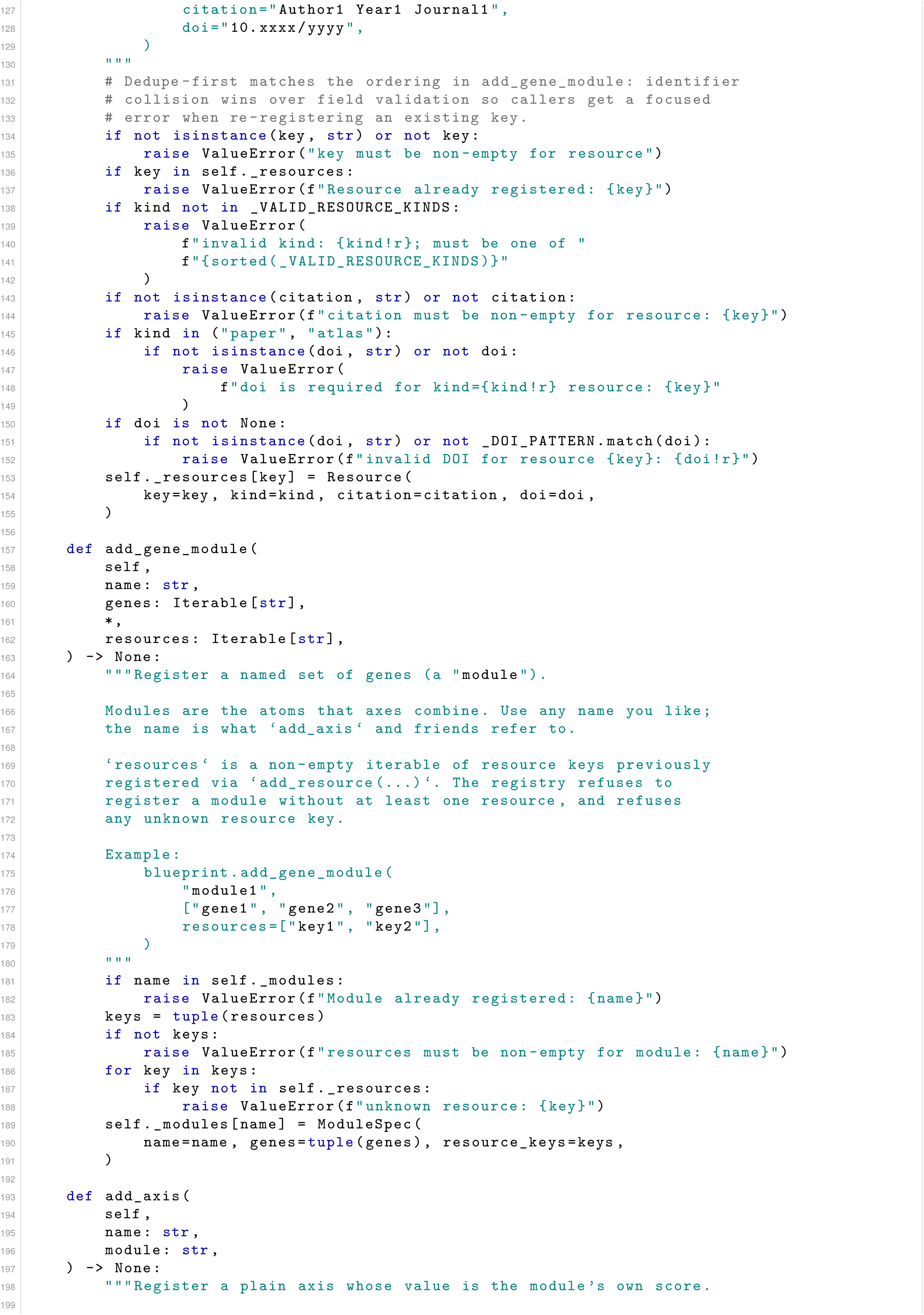
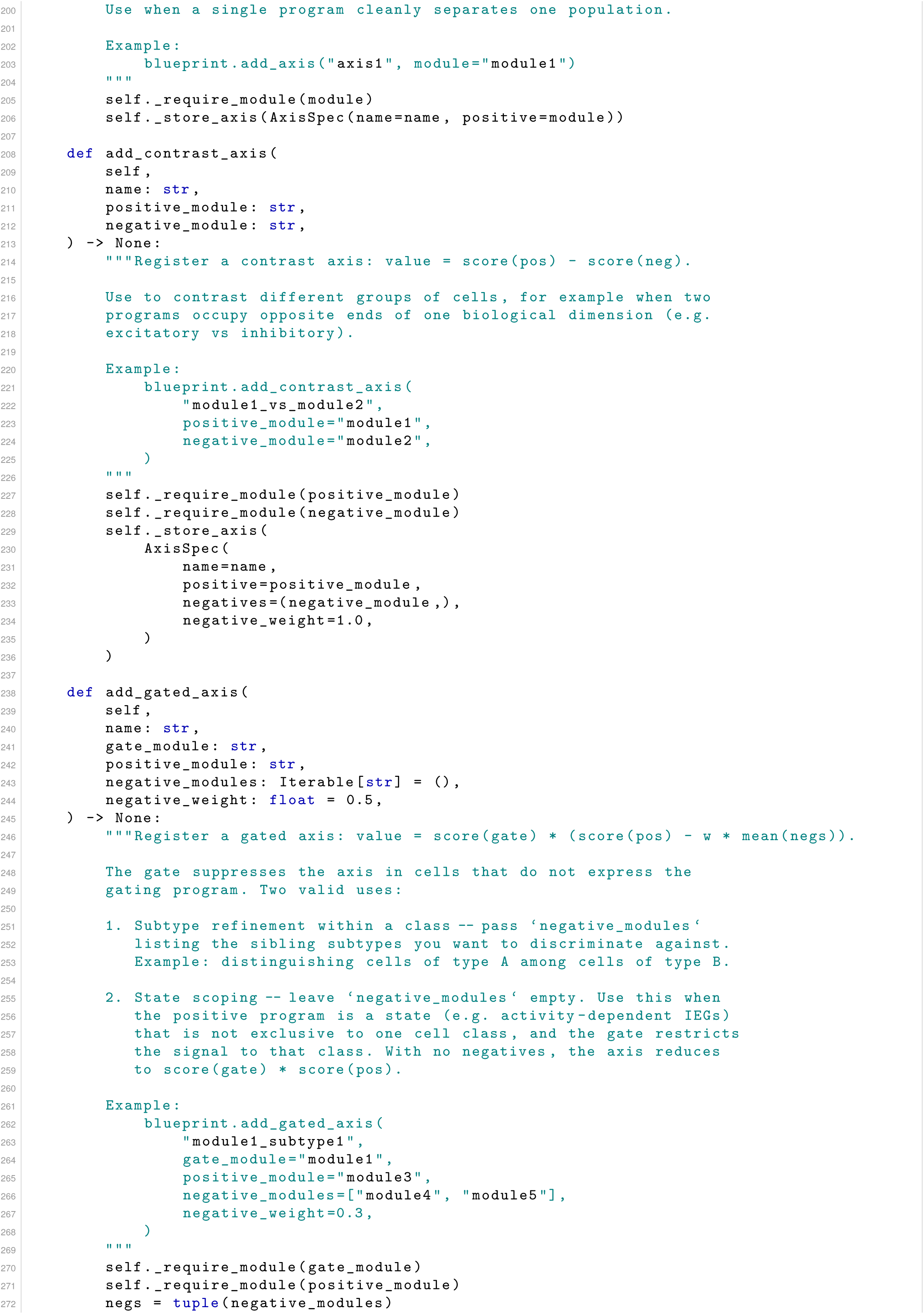
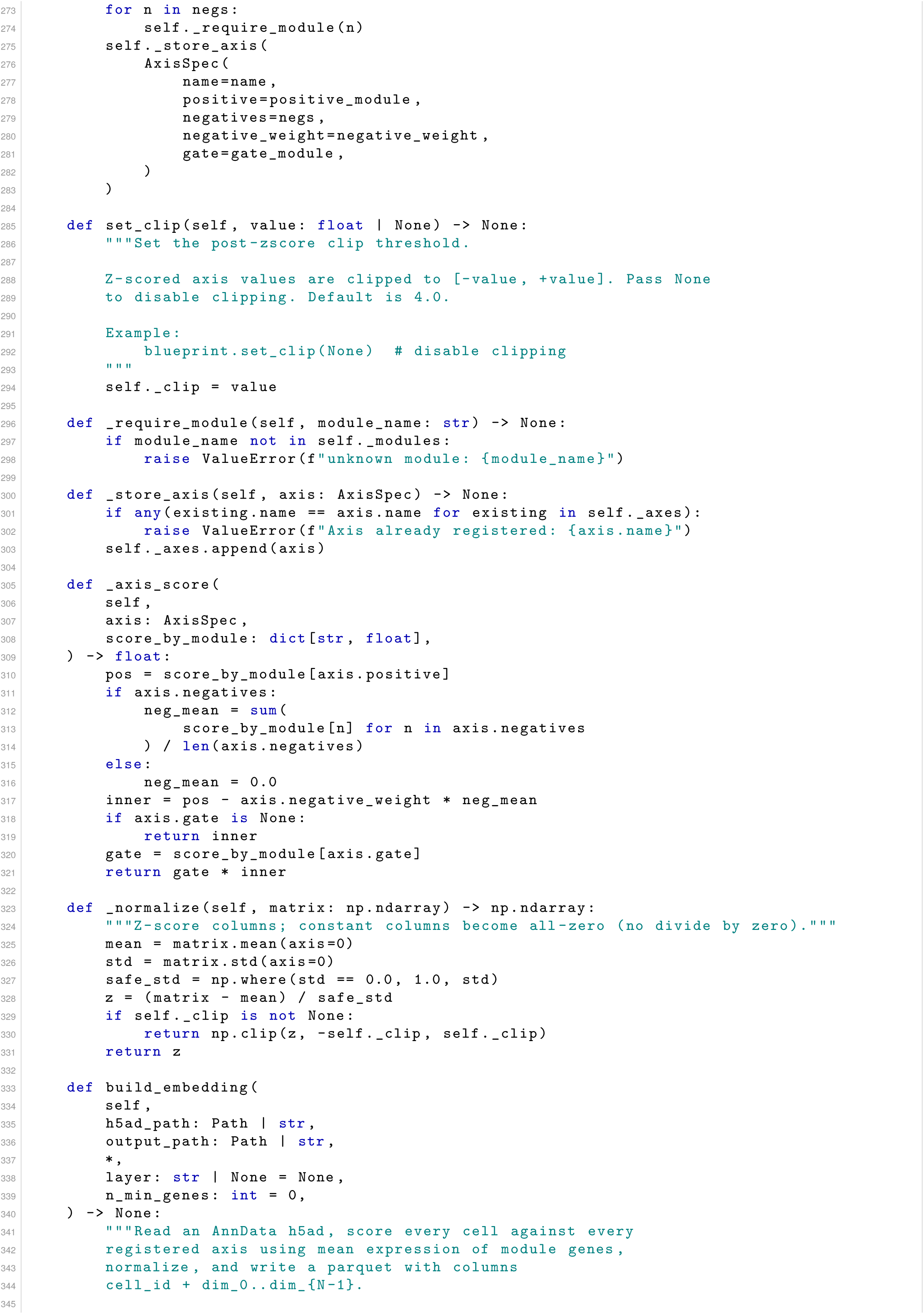
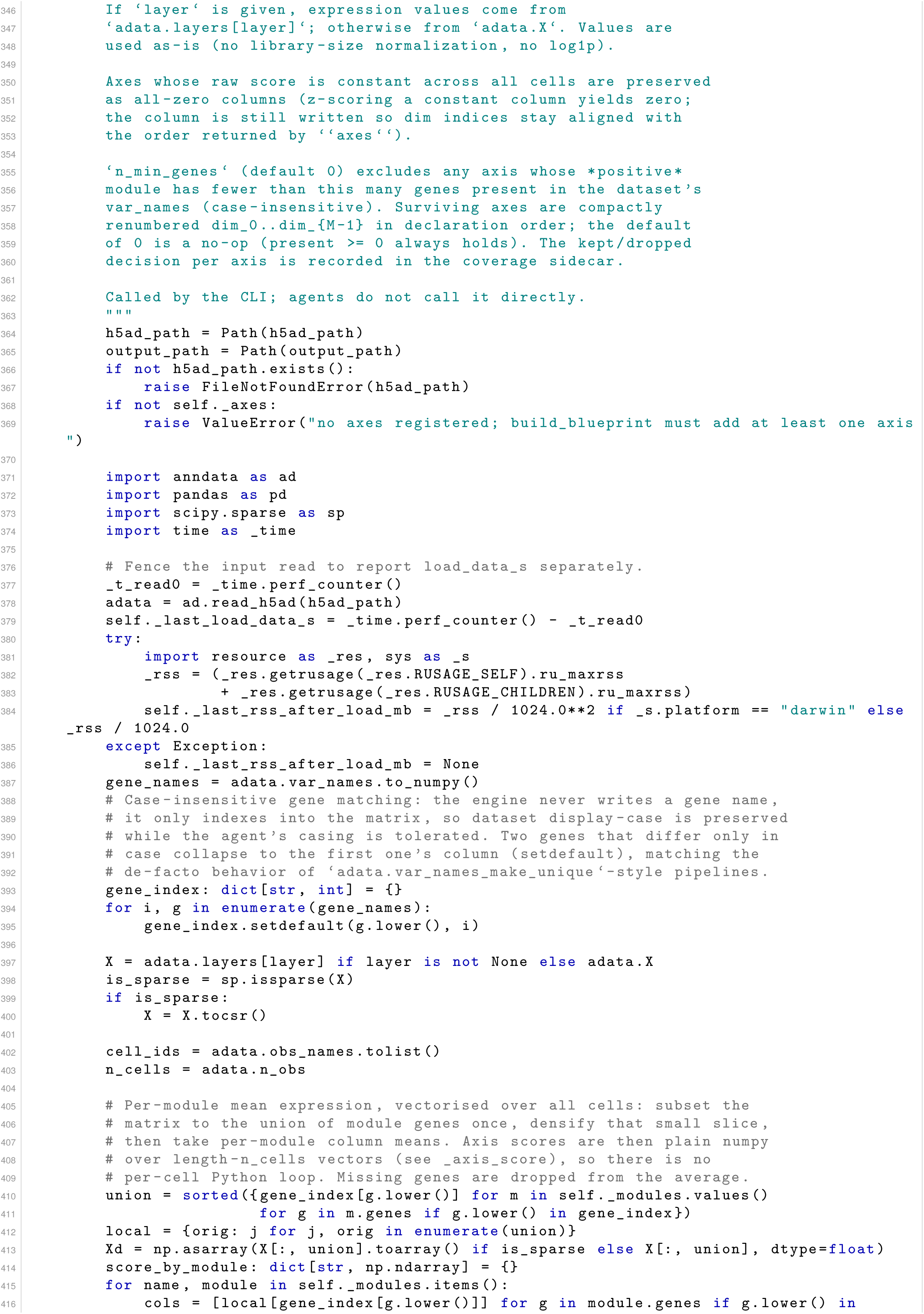
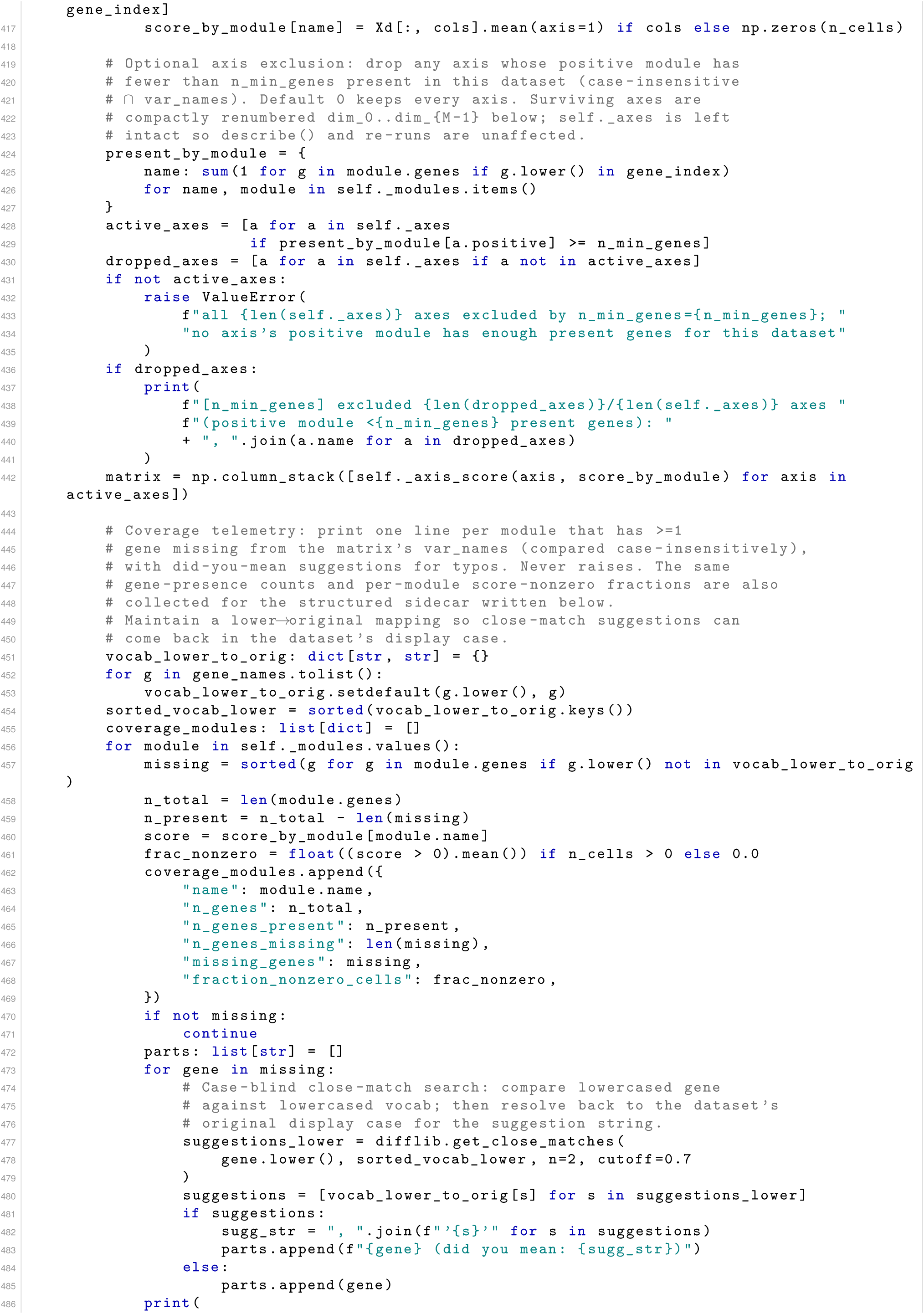
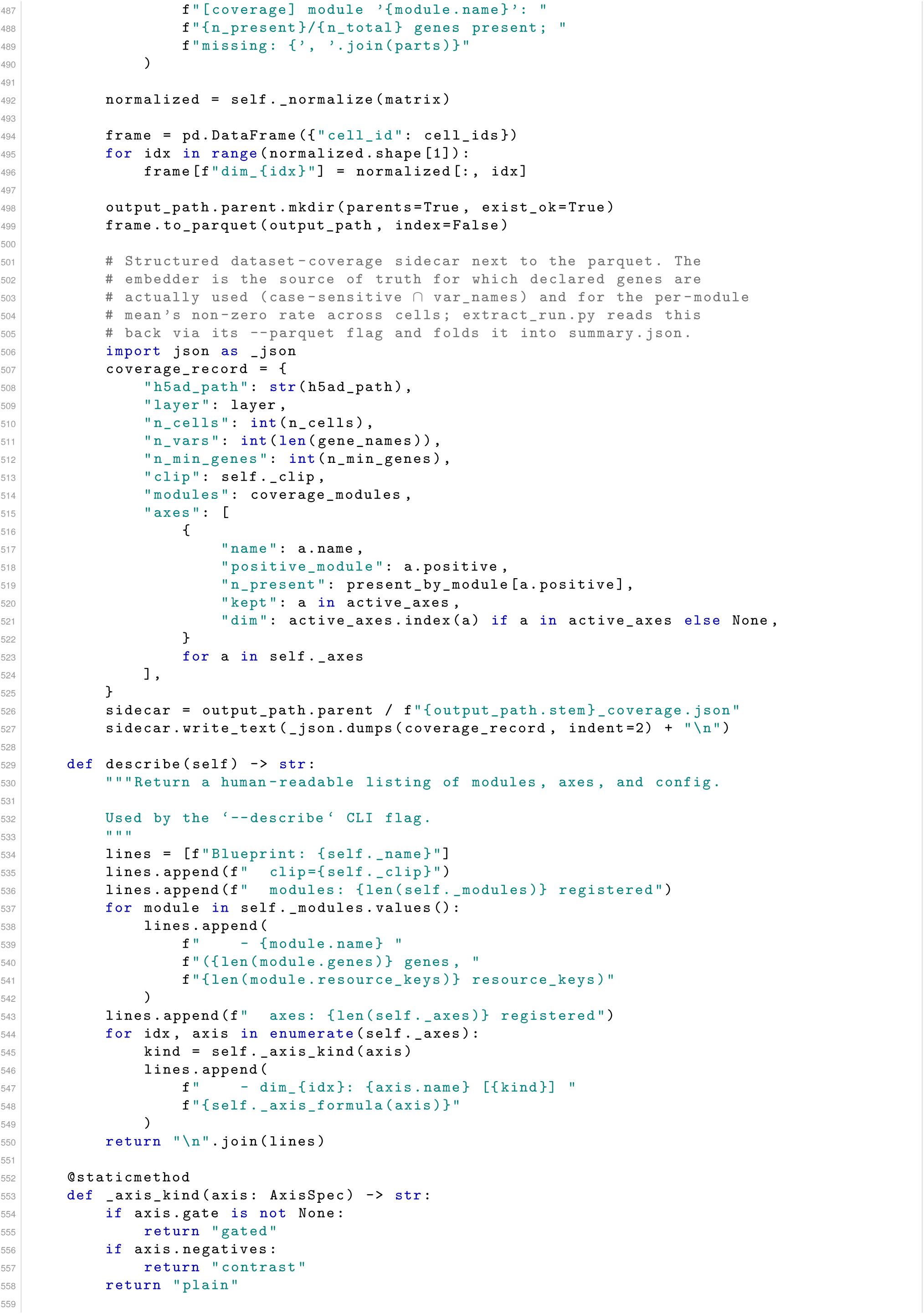
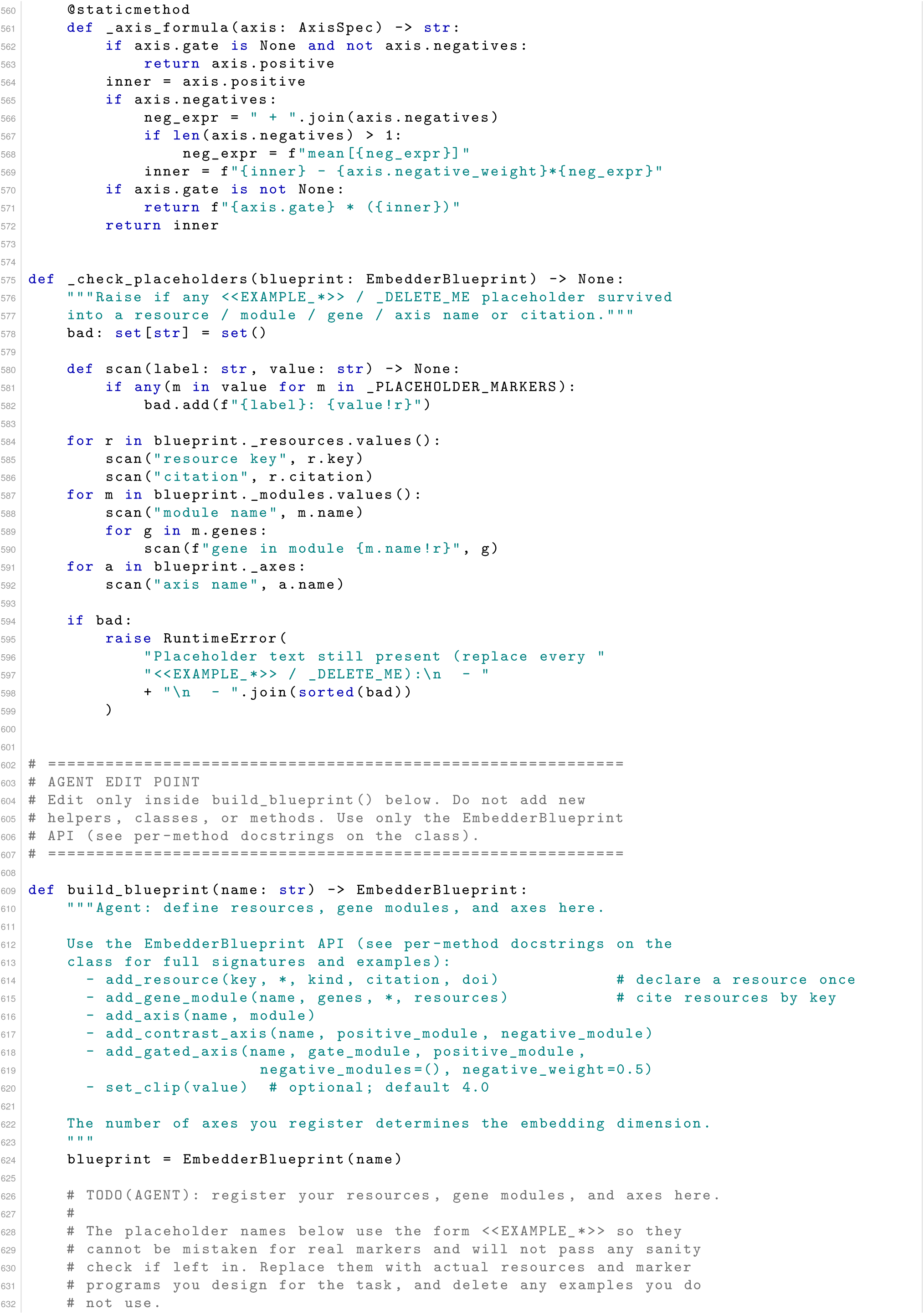
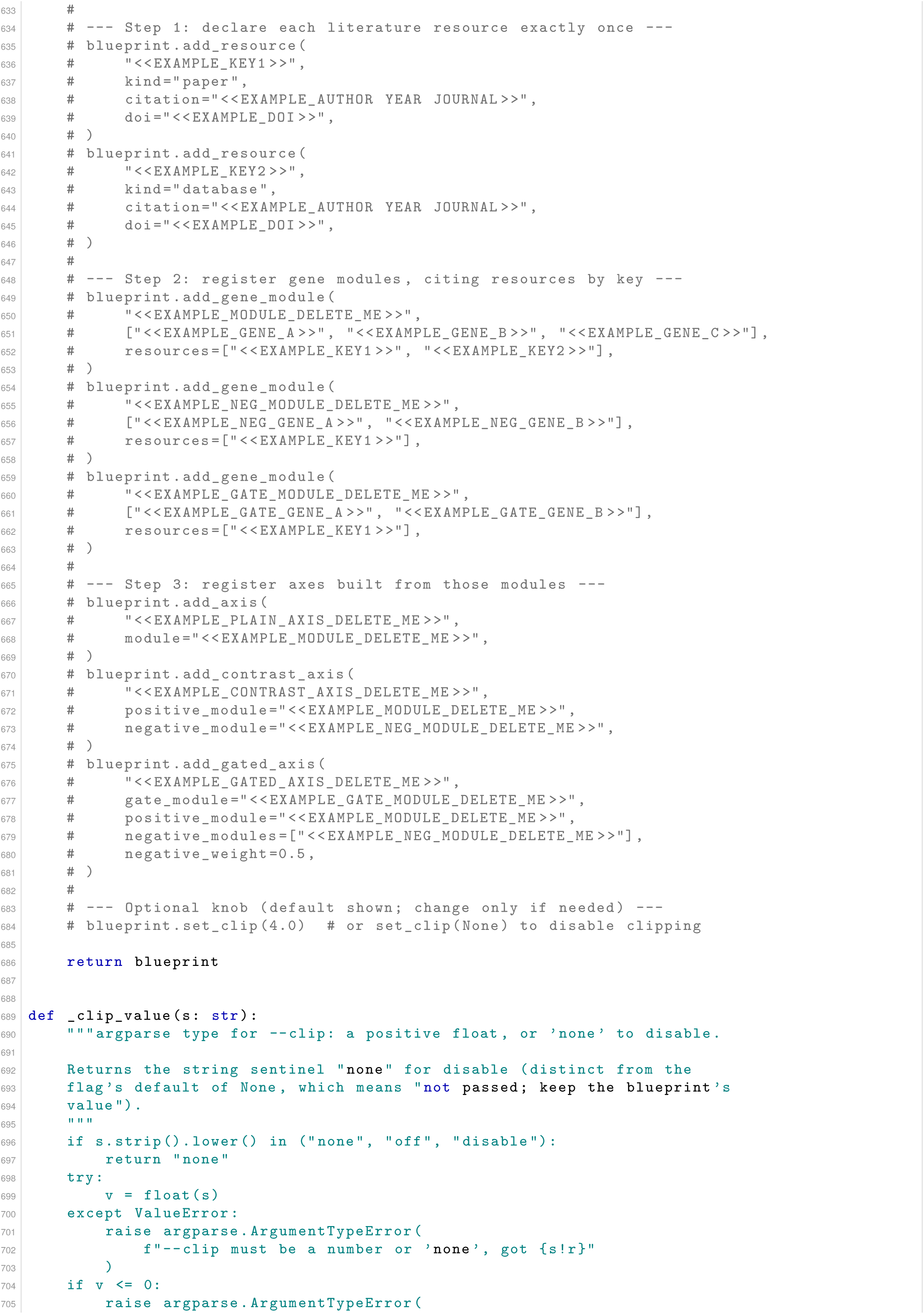
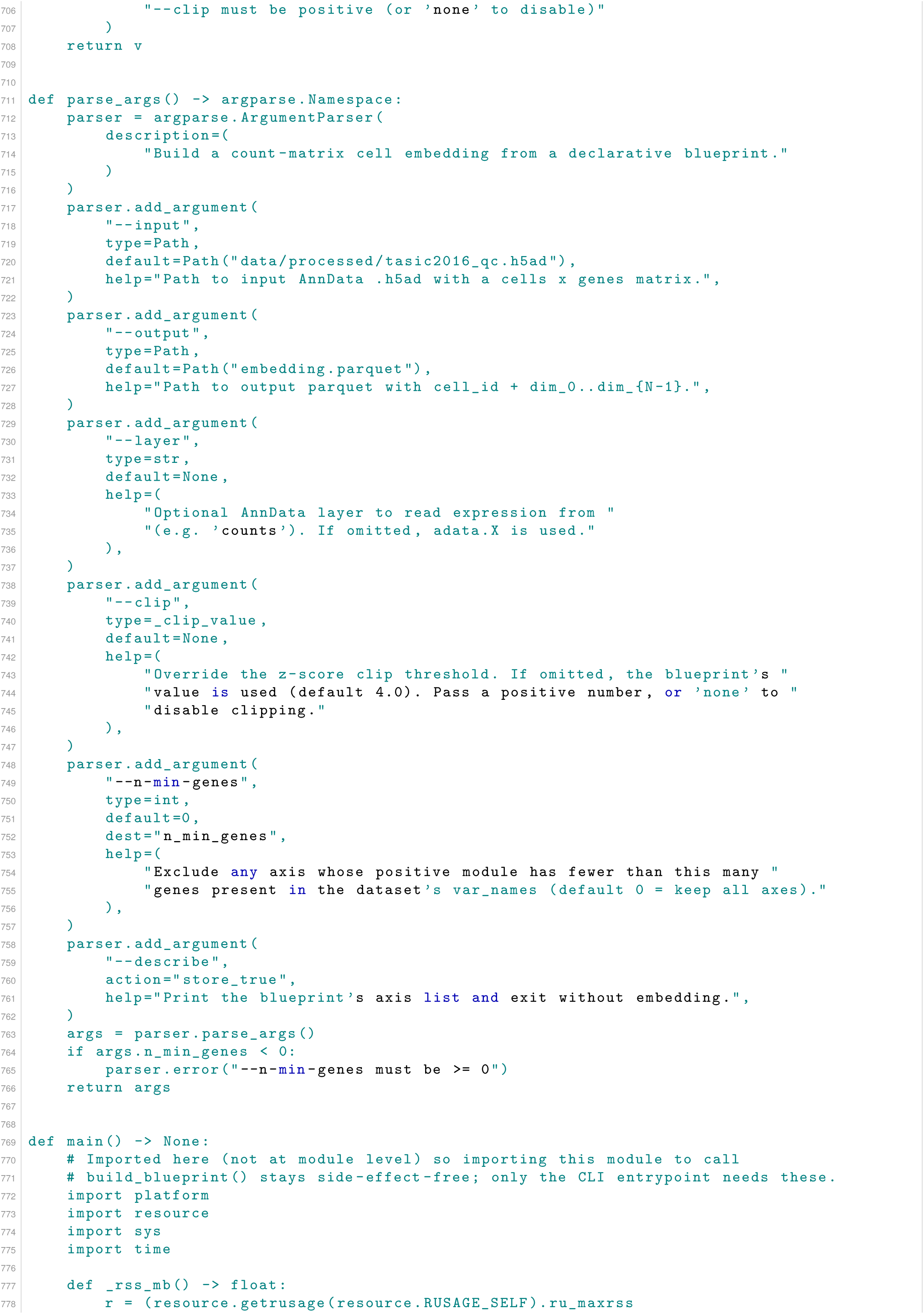
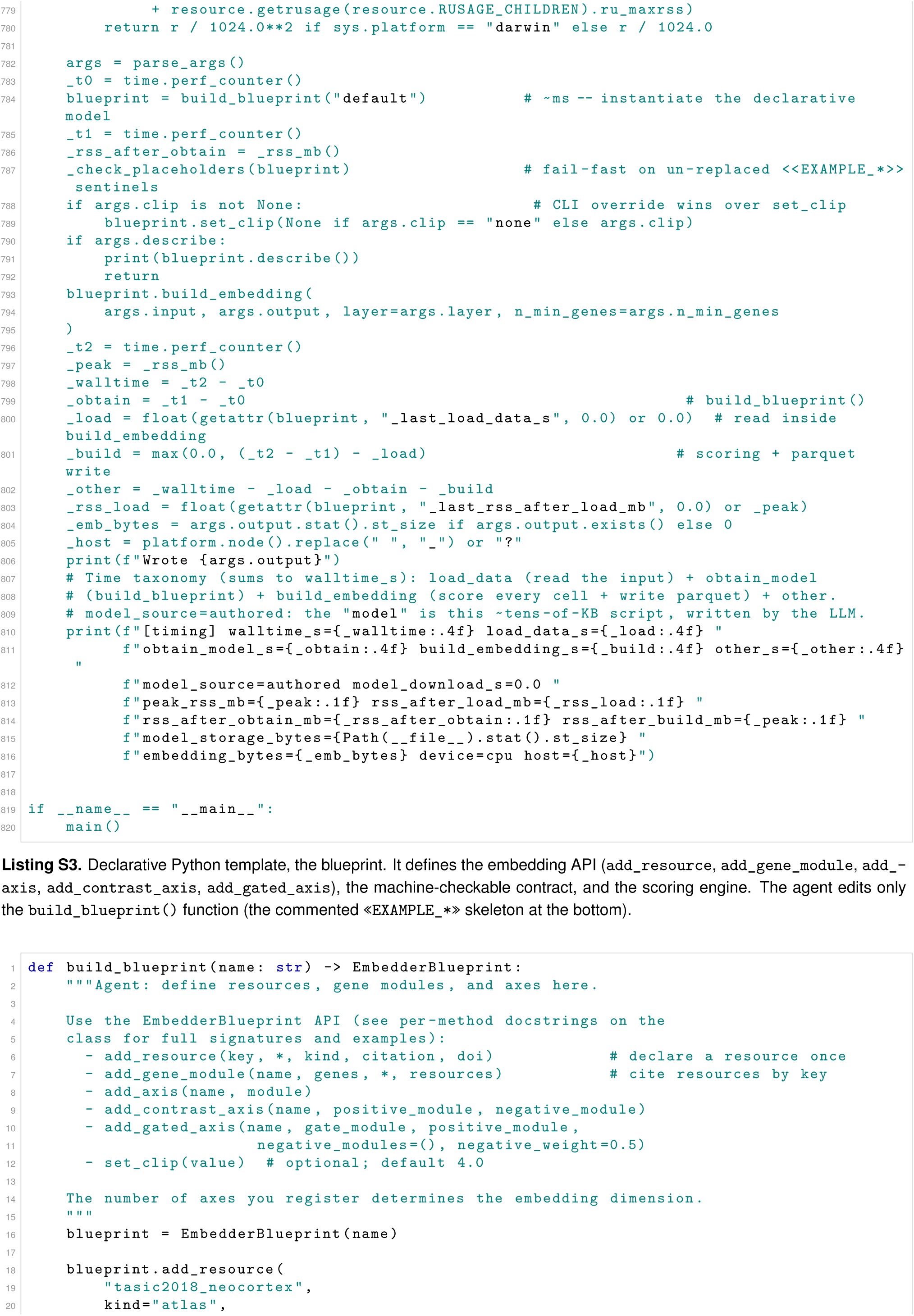
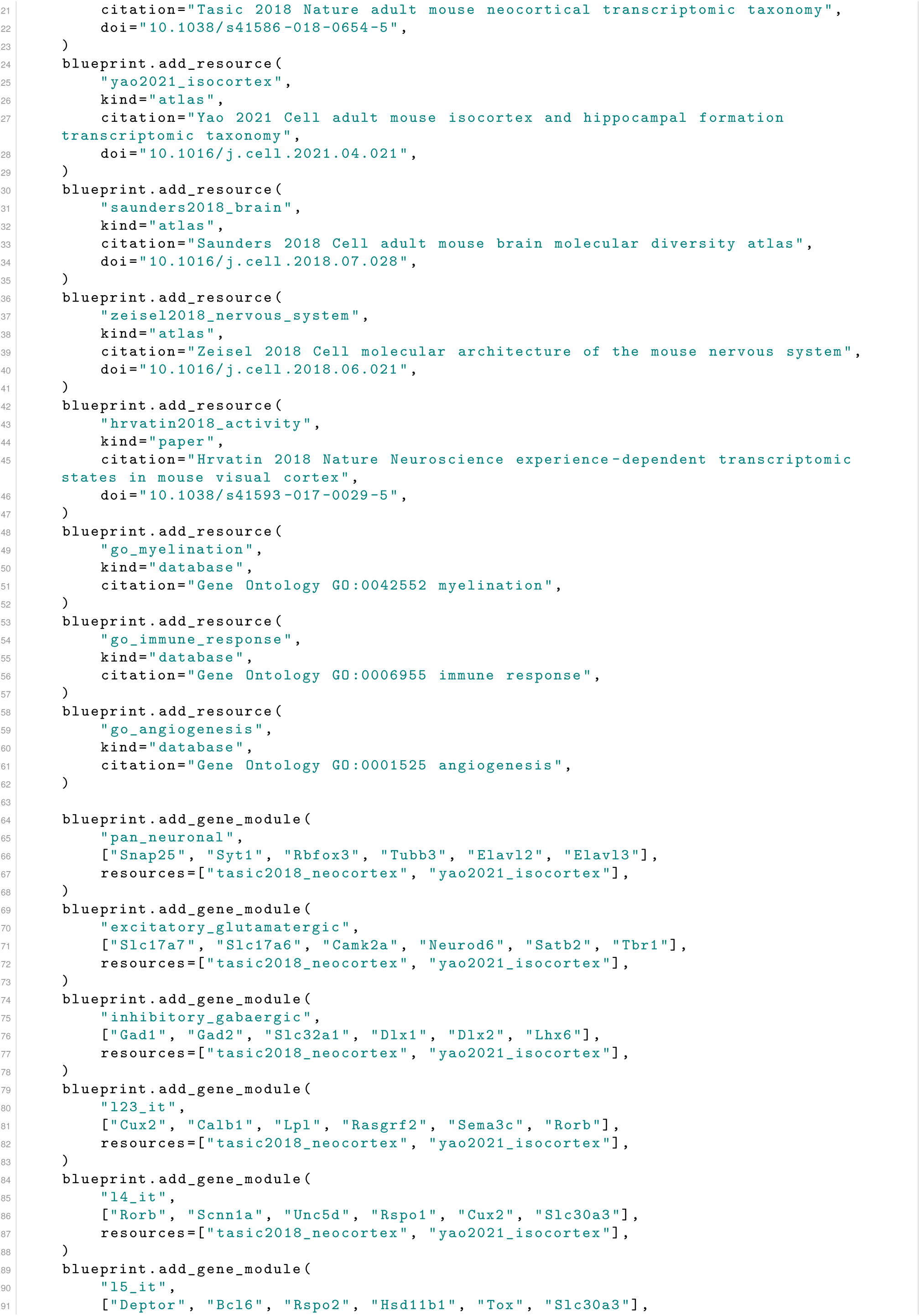
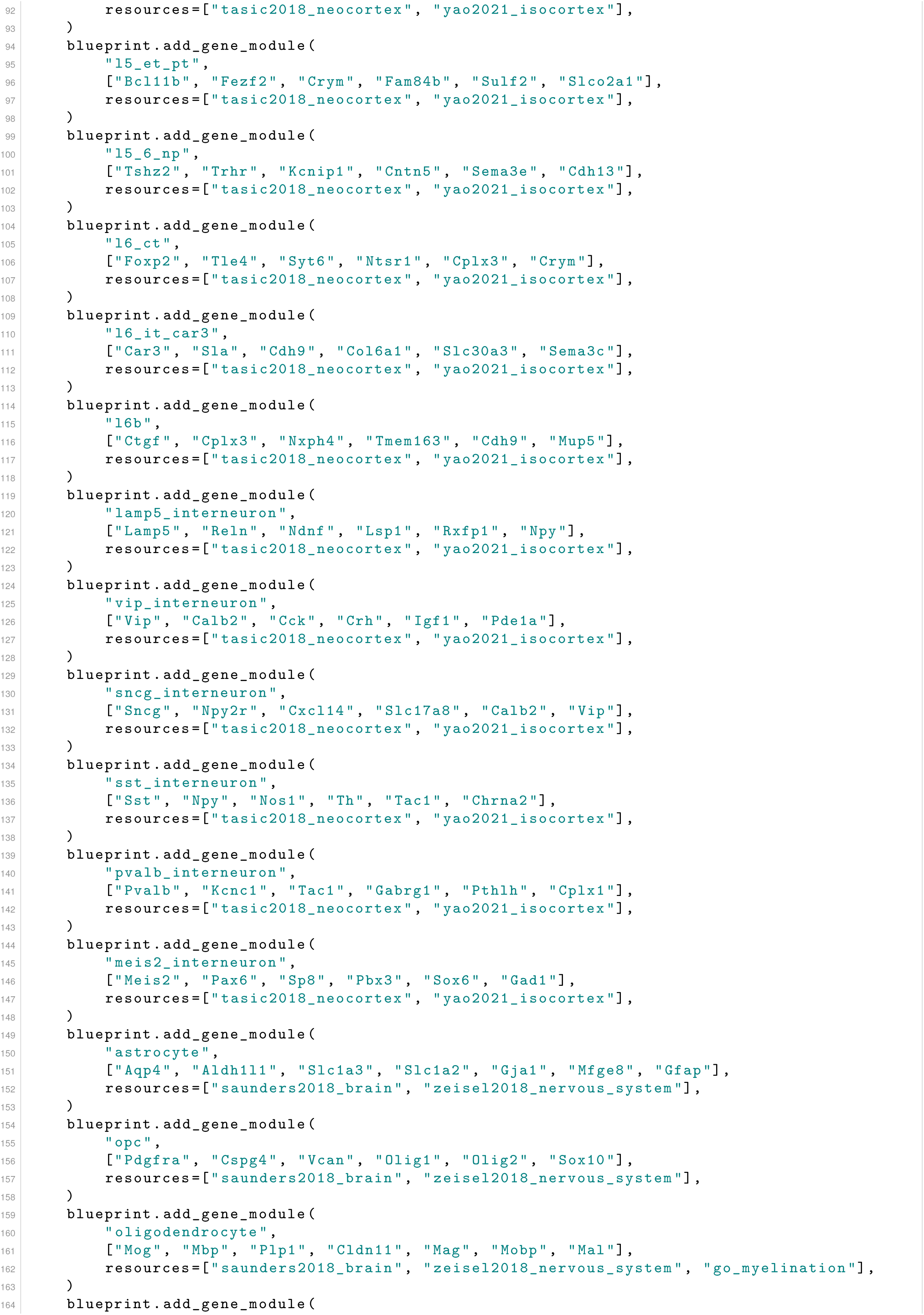
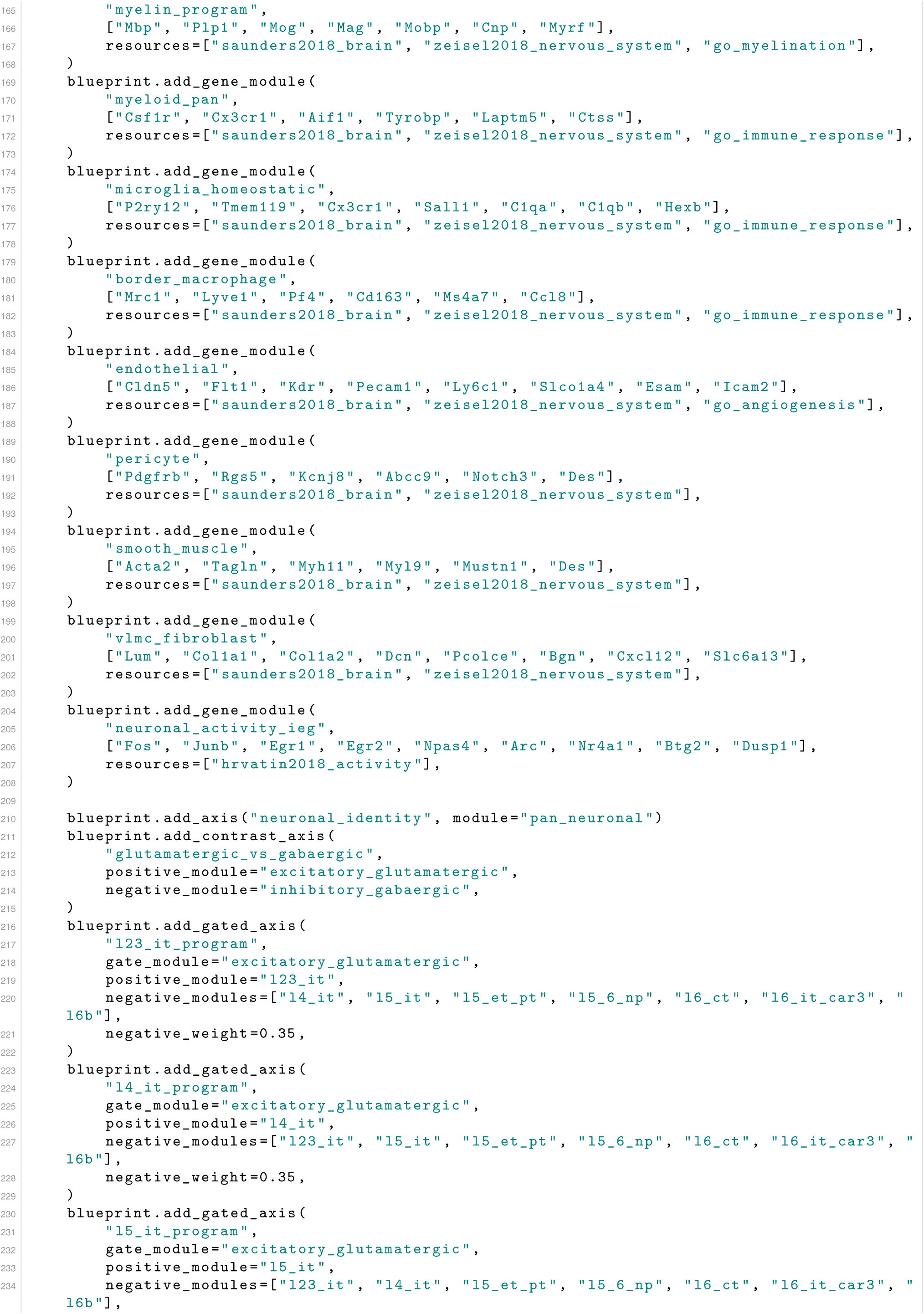
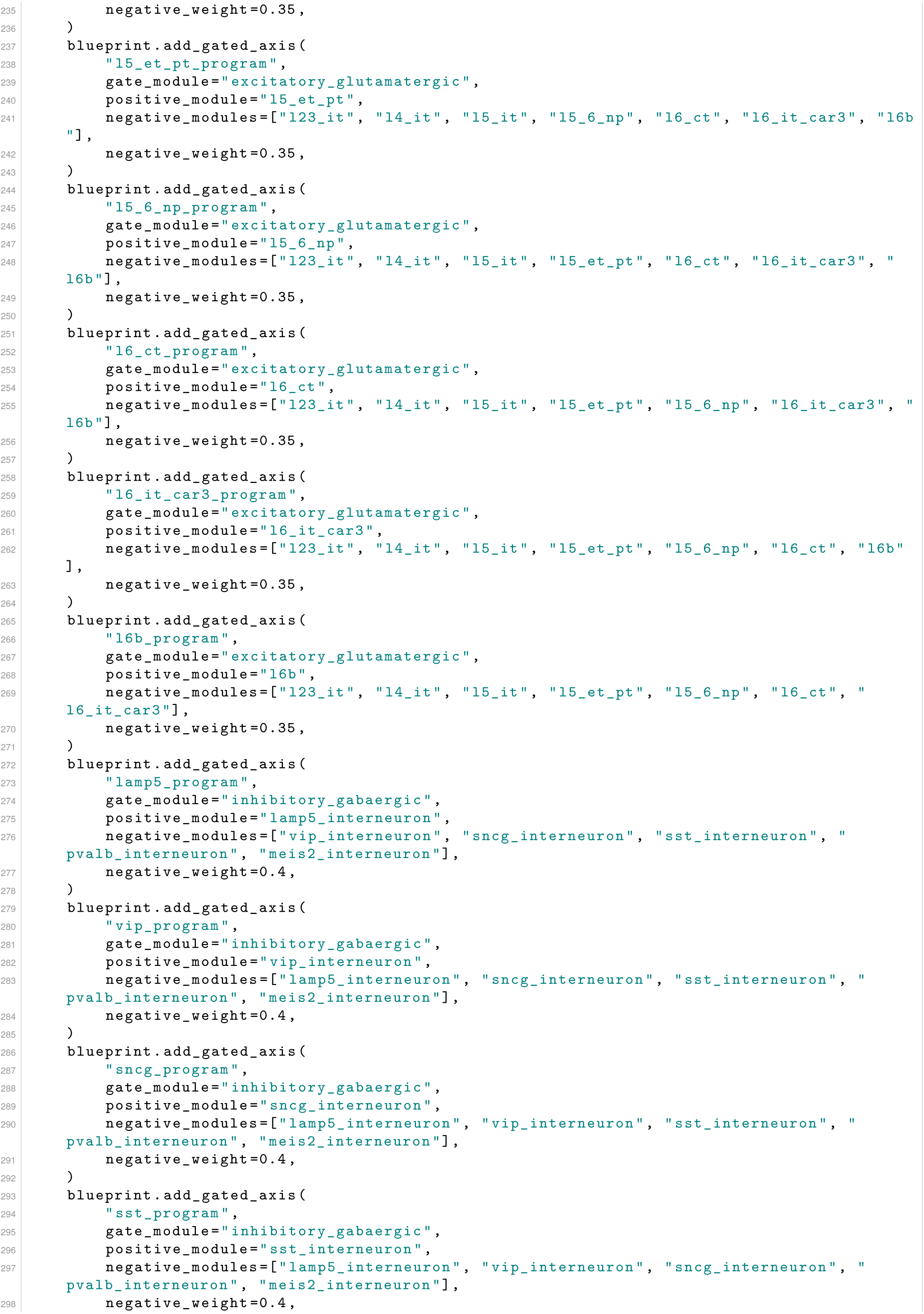
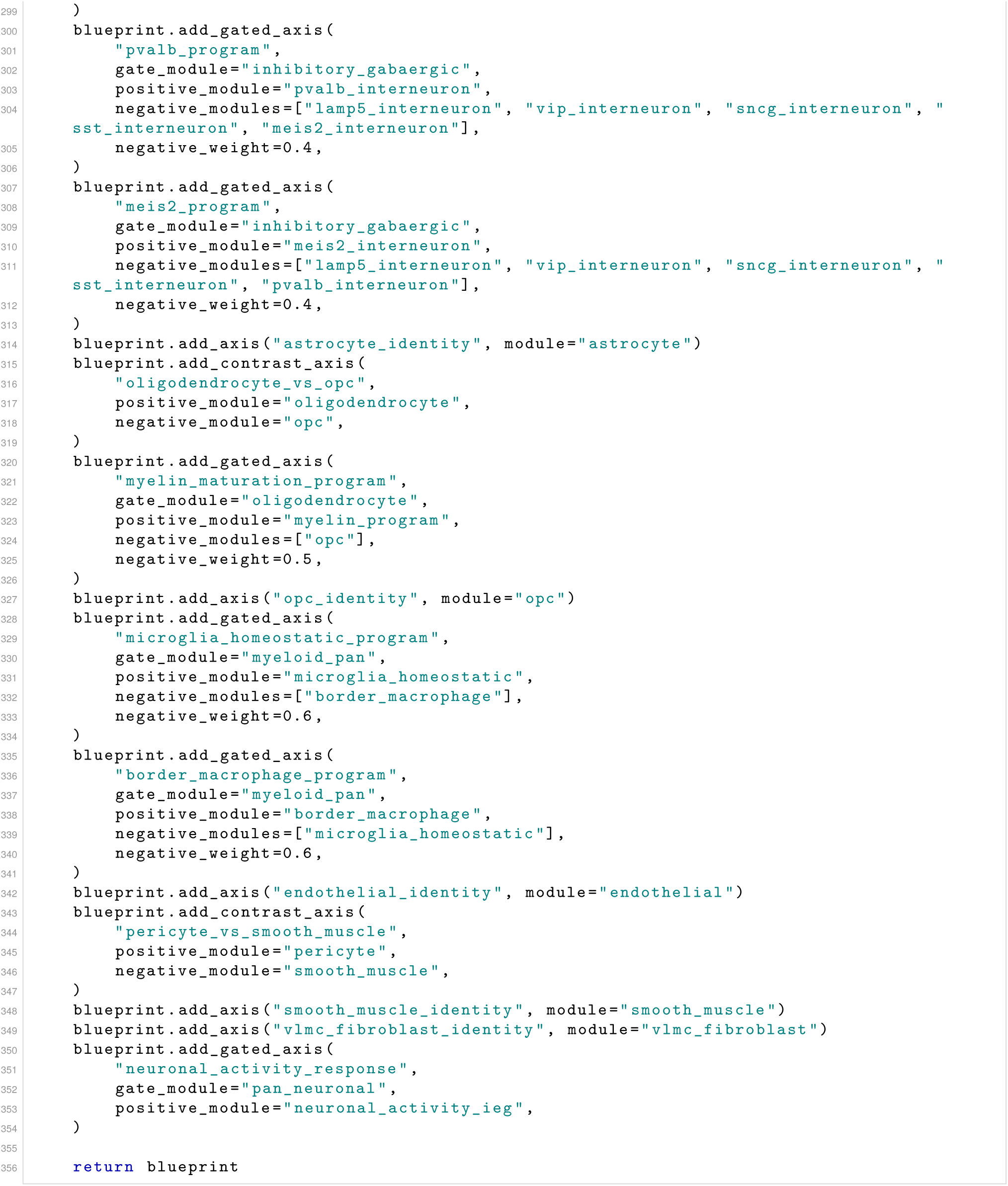

**Listing S4.** The build_blueprint() authored by the agent for the mouse_cortex showcase (seed 2): 8 literature resources (5
with DOIs, the other 3 curated databases), 29 named gene modules over 165 marker genes, composed into 27 axes (6 plain, 3
contrast, 18 gated).

**Table 2.**
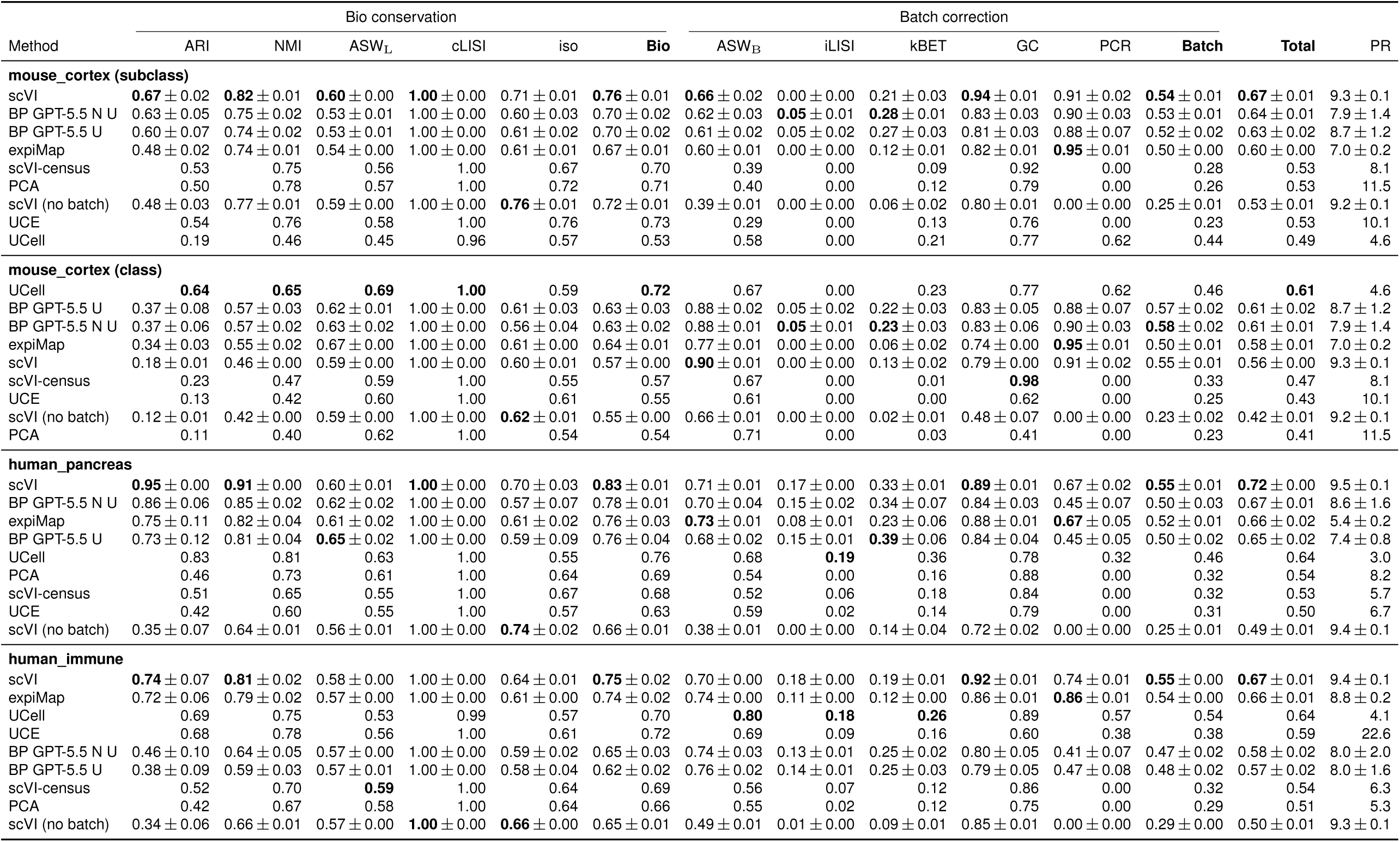
Blueprints reach competitive quality and are batch-invariant by construction. Full scIB benchmark numbers (mean ± s.d. across seeds). Every scIB sub-metric, grouped into the bio-conservation block (ARI, NMI, ASW_L_ = ASW-label, cLISI, iso = isolated-label *F*_1_) and the batch-correction block (ASW_B_ = ASW-batch, iLISI, kBET, GC = graph connectivity, PCR), each closed by its aggregate sub-score (**Bio**, **Batch**). **Total** = 0.6 Bio+0.4 Batch (the scIB package Total) and PR = participation ratio (effective dimensionality). Rows are the two blueprint grounding variants (BP GPT-5.5 U = literature-grounded, BP GPT-5.5 N U = resource-unconstrained, where U = unrefined and N = noref), the program- and marker-based comparators (UCell, expiMap), and the unsupervised and foundation baselines, grouped by benchmark dataset and cell-type resolution and sorted by Total. The best mean per column within each dataset block is in bold (PR excluded, as it is not a quality score). Deterministic methods (PCA, scVI-census, UCE, UCell) are scored once and carry no s.d.

**Figure S2.**
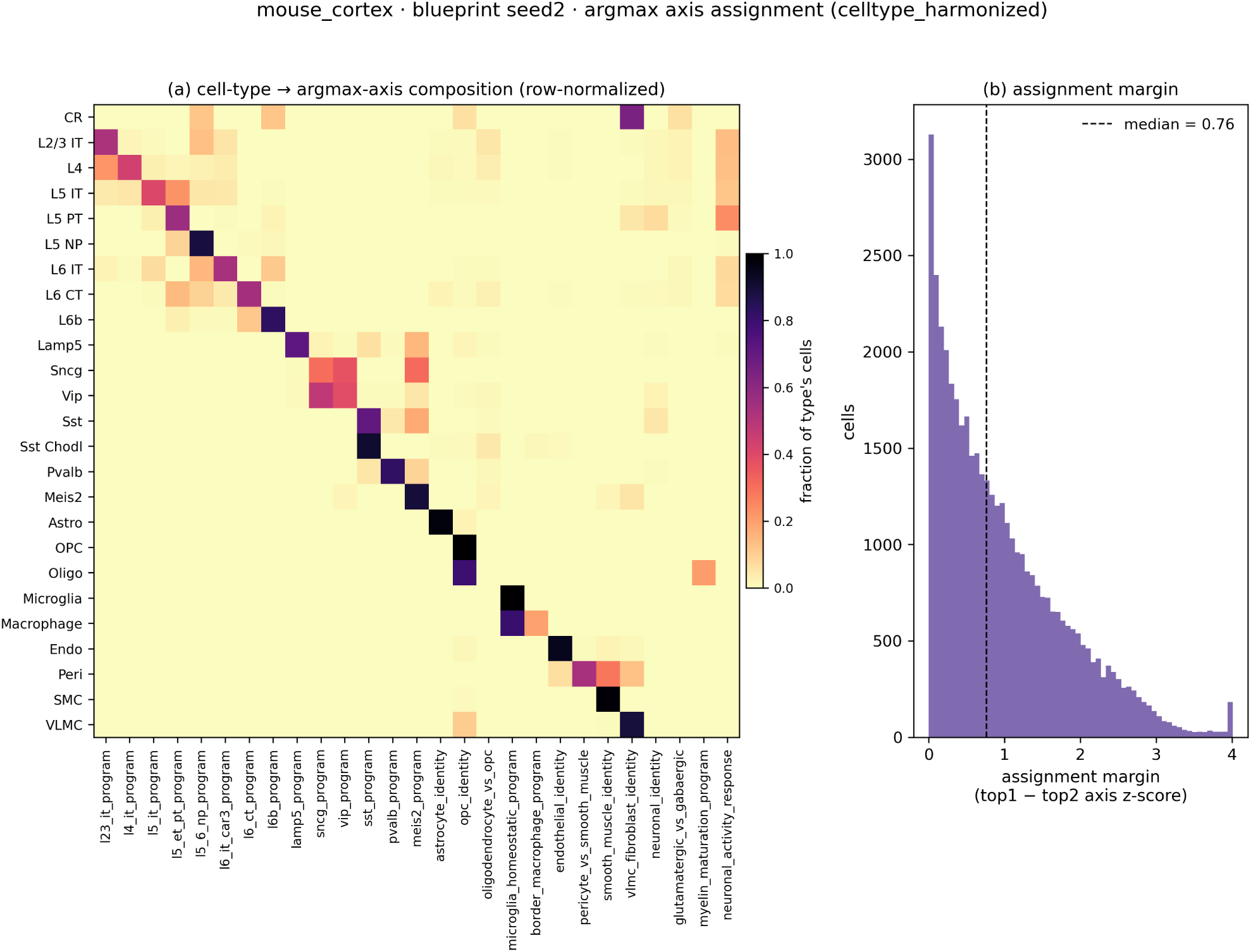
Assigning each cell to its most-activated axis recovers the annotated cell types. A descriptive complement to the threshold-free AUROC view (main-text Figure 1d). (**a**) Row-normalized composition: for each annotated cell type, the fraction of its cells whose argmax over the *z*-scored axes falls on each axis. The near-block-diagonal shows each type is captured by its own named axis. (**b**) Distribution of the assignment margin (the gap between the top and second axis *z*-score per cell).

**Interpretability and faithfulness (mouse cortex): supplementary detail Frontier benchmark: per-dataset detail**

**Held-out reference-to-query transfer**

**Structured agentic framework: specification recurrence and cost Developmental-tree showcase: supplementary detail**

**Figure S3.**
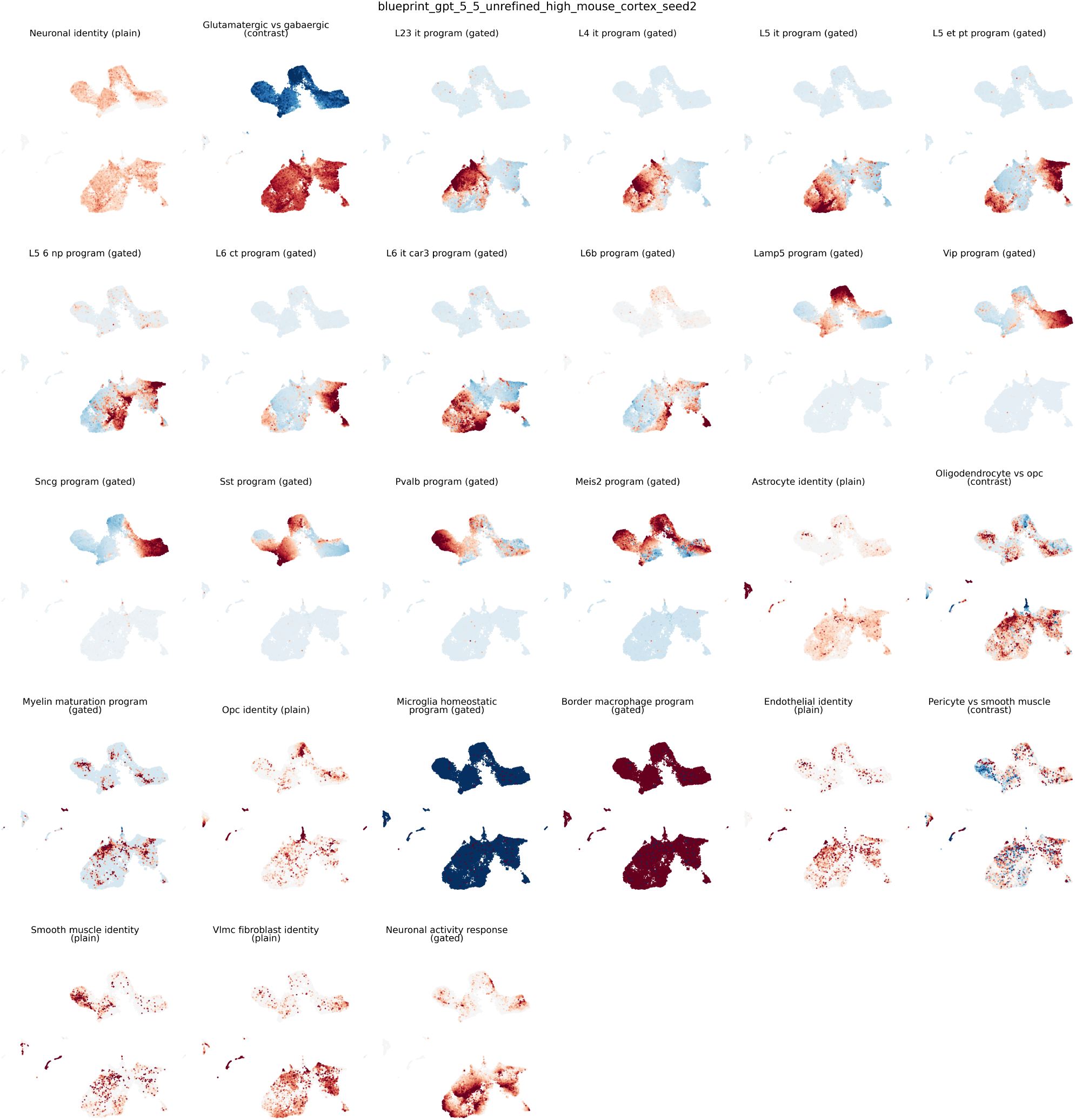
Each authored axis localizes to a coherent region of the expression manifold. Per-axis activation maps for the representative mouse_cortex blueprint: one UMAP of the cells (a single shared layout) recoloured by each embedding axis’s activation value, titled with the axis name and kind parsed from the run. The named identity axes light up over compact, distinct territories matching their cell types, while the contrast and gated program axes sweep along the corresponding biological gradients.

**Figure S4.**
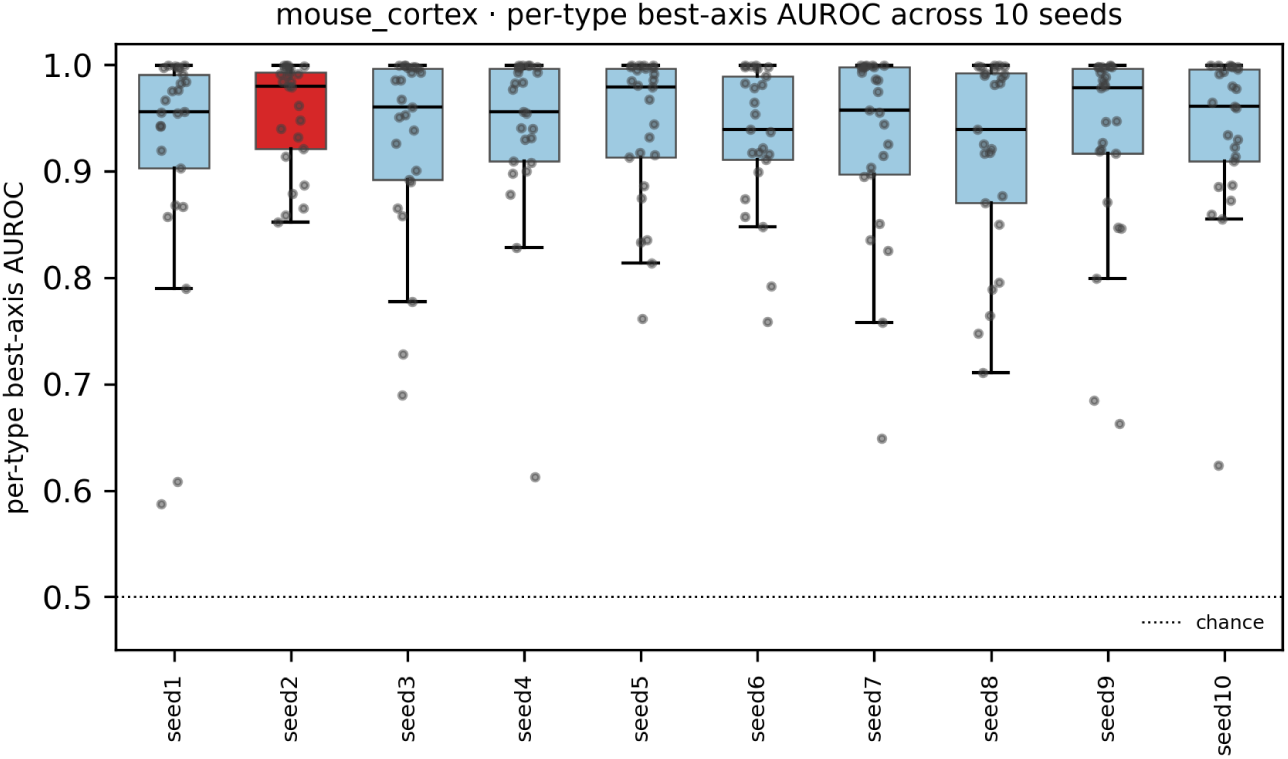
Per-axis faithfulness reproduces across re-authoring runs. For each of the ten independent authoring seeds of the mouse_cortex blueprint cohort, we show the distribution (over annotated cell types) of the best one-vs-rest AUROC any axis attains for the type, the per-type discriminability of the authored model. The per-type best-axis AUROC is consistently high and well above chance across all seeds, including the representative seed used in the main text (red).

**Figure S5.**
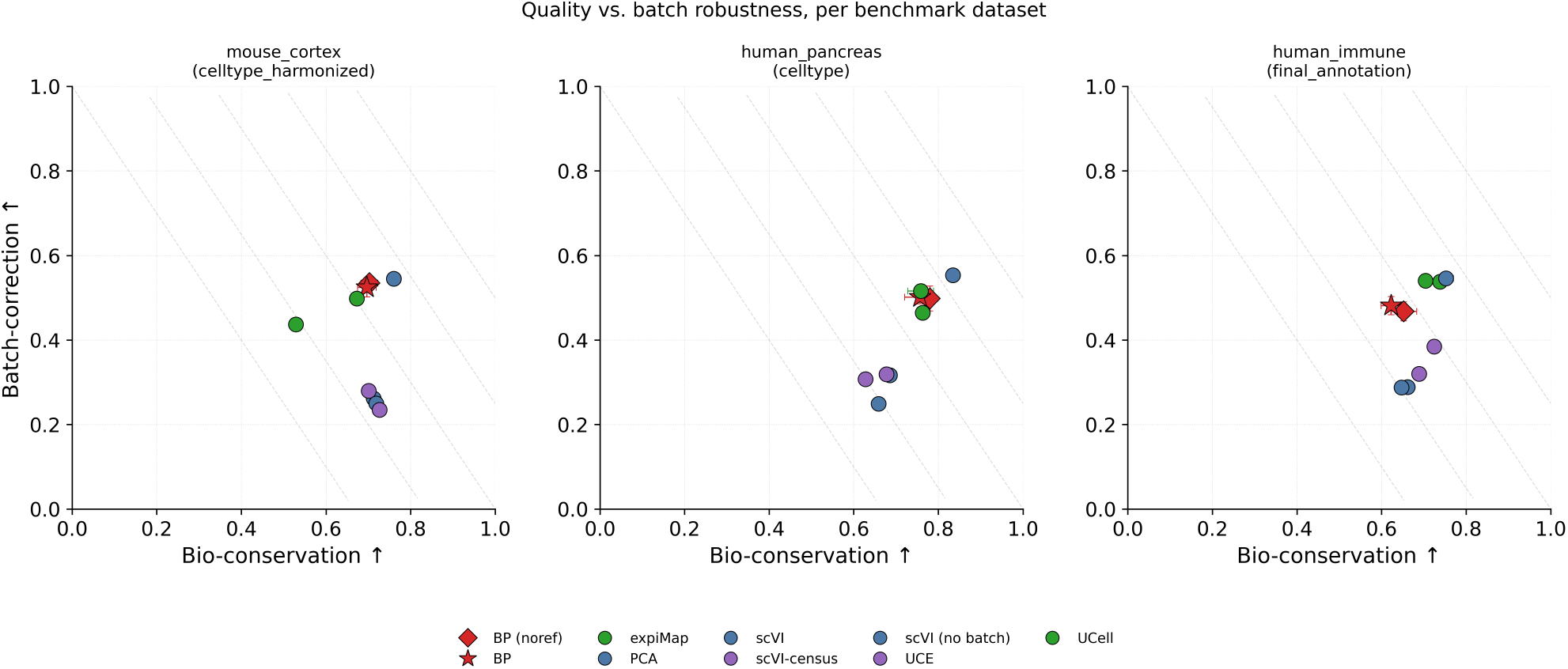
Quality versus batch robustness, per benchmark dataset. scIB bio-conservation against batch-correction, one marker per method (mean across seeds, error bars s.d. across seeds) at each dataset’s reported resolution, colored by method class. The two blueprint grounding variants are red, a star for literature-grounded and a diamond for noref. The program- and marker-based comparators UCell and expiMap share one method class, shown in green. Dashed lines connect points of equal composite score.

**Figure S6.**
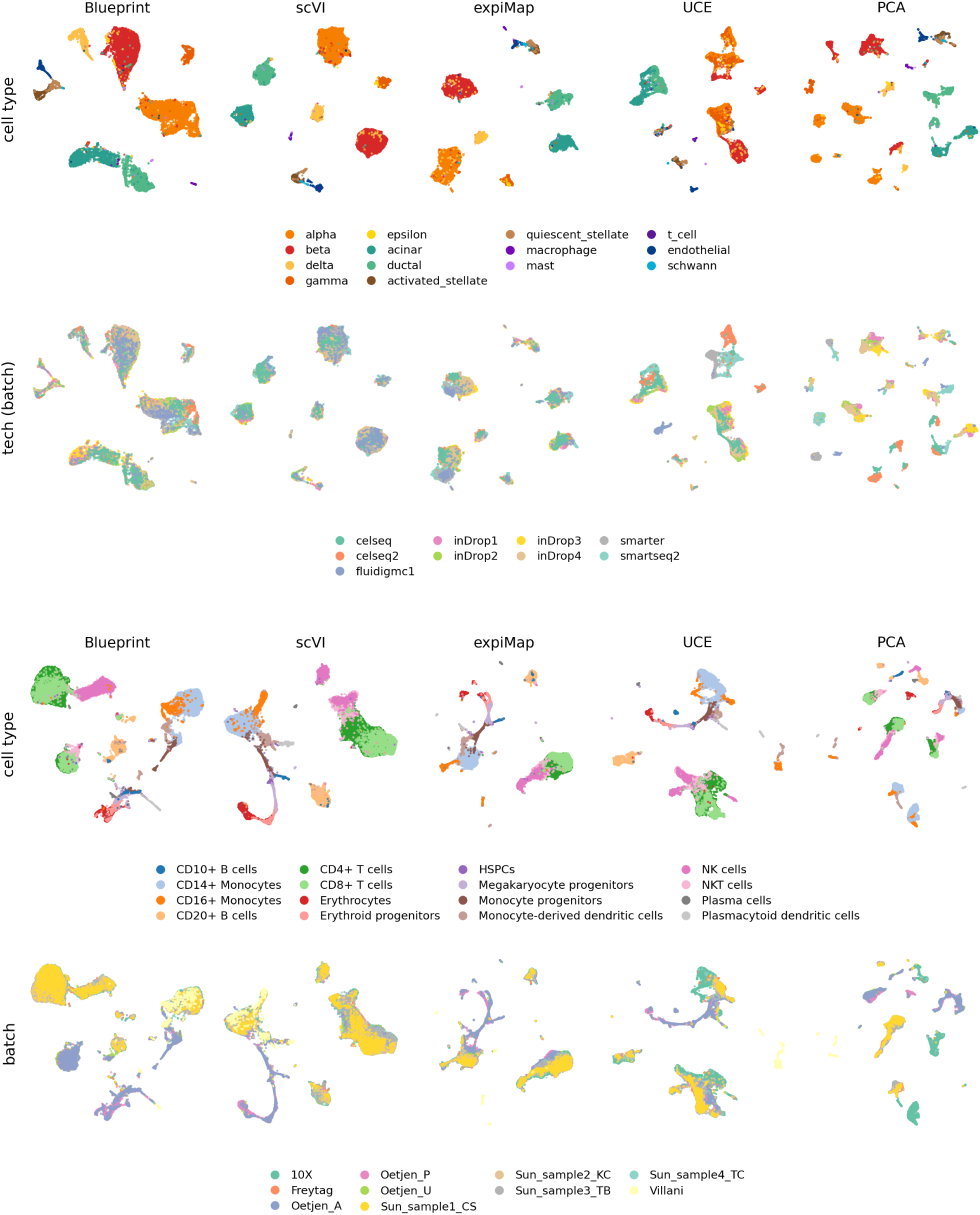
Cell-type and batch UMAPs for the human benchmark datasets. UMAP grids at the best seed per method (by scIB composite score), the top row colored by cell type and the bottom row by the batch covariate, for human_pancreas (top) and human_immune (bottom), the companions to the mouse_cortex grid in main-text Fig. 2c.

**Figure S7.**
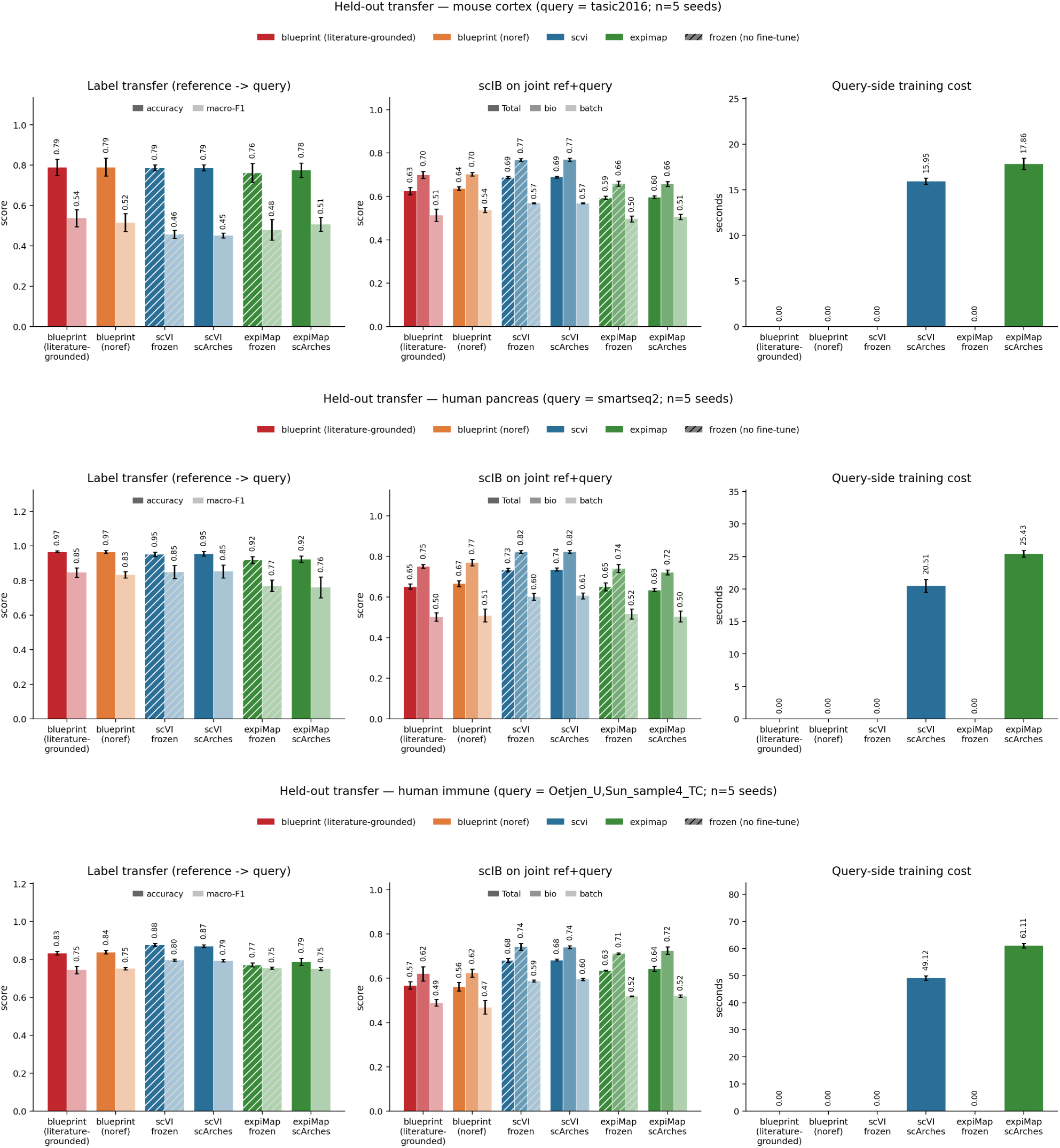
Held-out reference-to-query transfer, per benchmark dataset. An entire study, technology, or donor is held out as a query on each dataset. For mouse_cortex (top) the query is the full-length SMART-seq Tasic 2016 study, held out of the SMART-seq-v4 and droplet reference. For human_pancreas (middle) it is the SMART-seq2 technology, held out of the droplet reference. For human_immune (bottom) it is one bone-marrow donor (Oetjen_U) and one PBMC sample (Sun_sample4_TC), held out of the remaining batches. scVI and expiMap are trained on the reference only and mapped onto the query either frozen (no query-side fine-tune, hatched bars) or by scArches surgery, whereas the blueprint (literature-grounded and noref) is applied zero-shot. Bars are the mean over five seeds and error bars are s.d. The left panels show reference-to-query *k*NN label transfer (accuracy and macro-*F*_1_). The middle panels show the full scIB suite on the joint reference + query embedding (Total with its bio-and batch-conservation sub-scores), the same metrics as the main benchmark. The right panels show the query-side training (surgery) wall-clock, zero for the zero-shot blueprint and the frozen projections.

**Figure S8.**
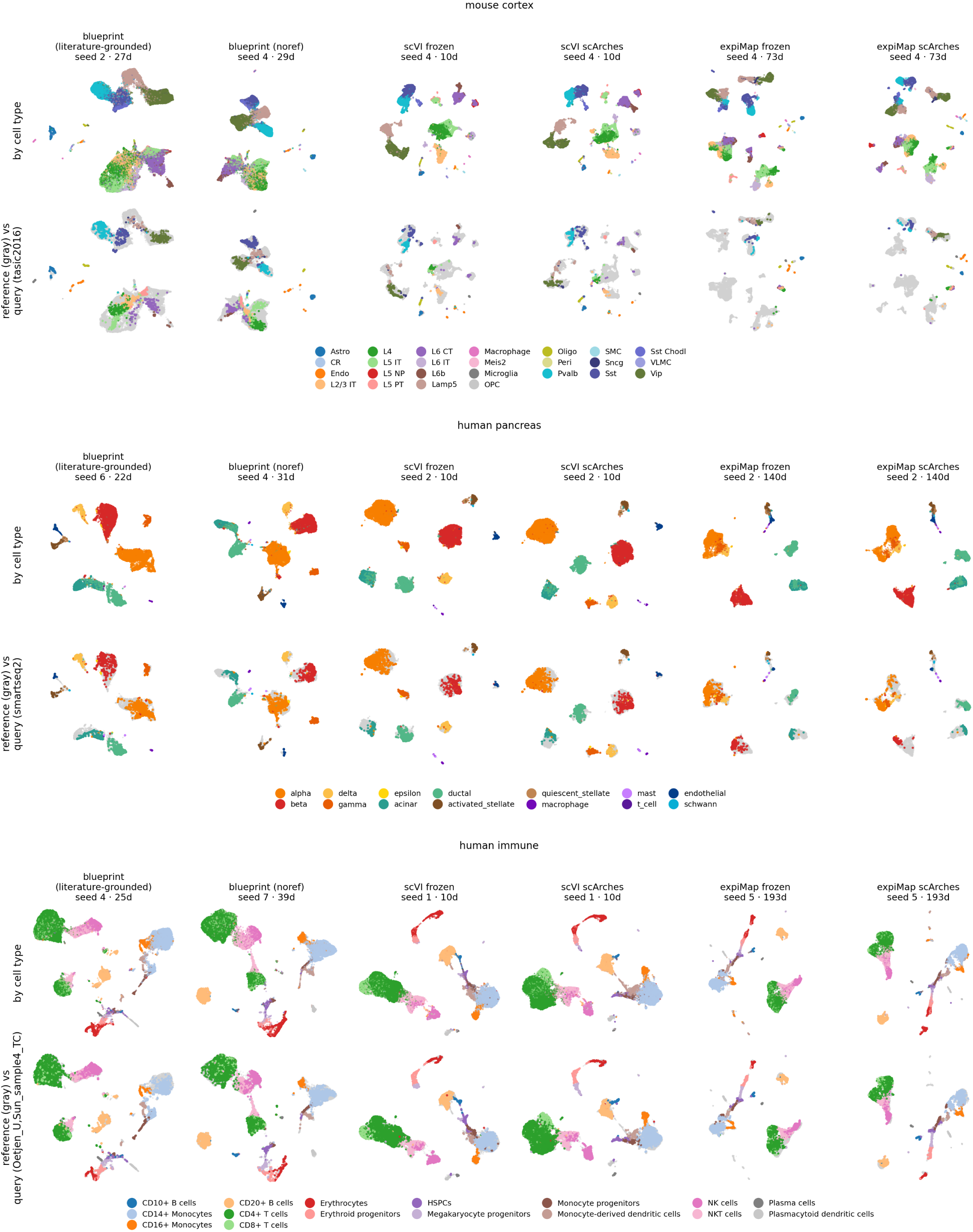
Held-out transfer UMAPs, per benchmark dataset. UMAP of the joint reference + query embedding for each method variant, for mouse_cortex, human_pancreas, and human_immune, stacked top to bottom. Within each, the upper row is colored by cell type and the lower row shows the reference in gray with the held-out query cells colored by cell type and drawn on top. Each panel uses its method’s best-composite-scIB seed. The blueprint panels reuse the exact full-dataset coordinates of Fig. 2c and Supp. Fig. S6, since its zero-shot joint embedding is identical to the full-dataset embedding, and the scVI and expiMap panels share that seed across their frozen and scArches columns.

**Figure S9.**
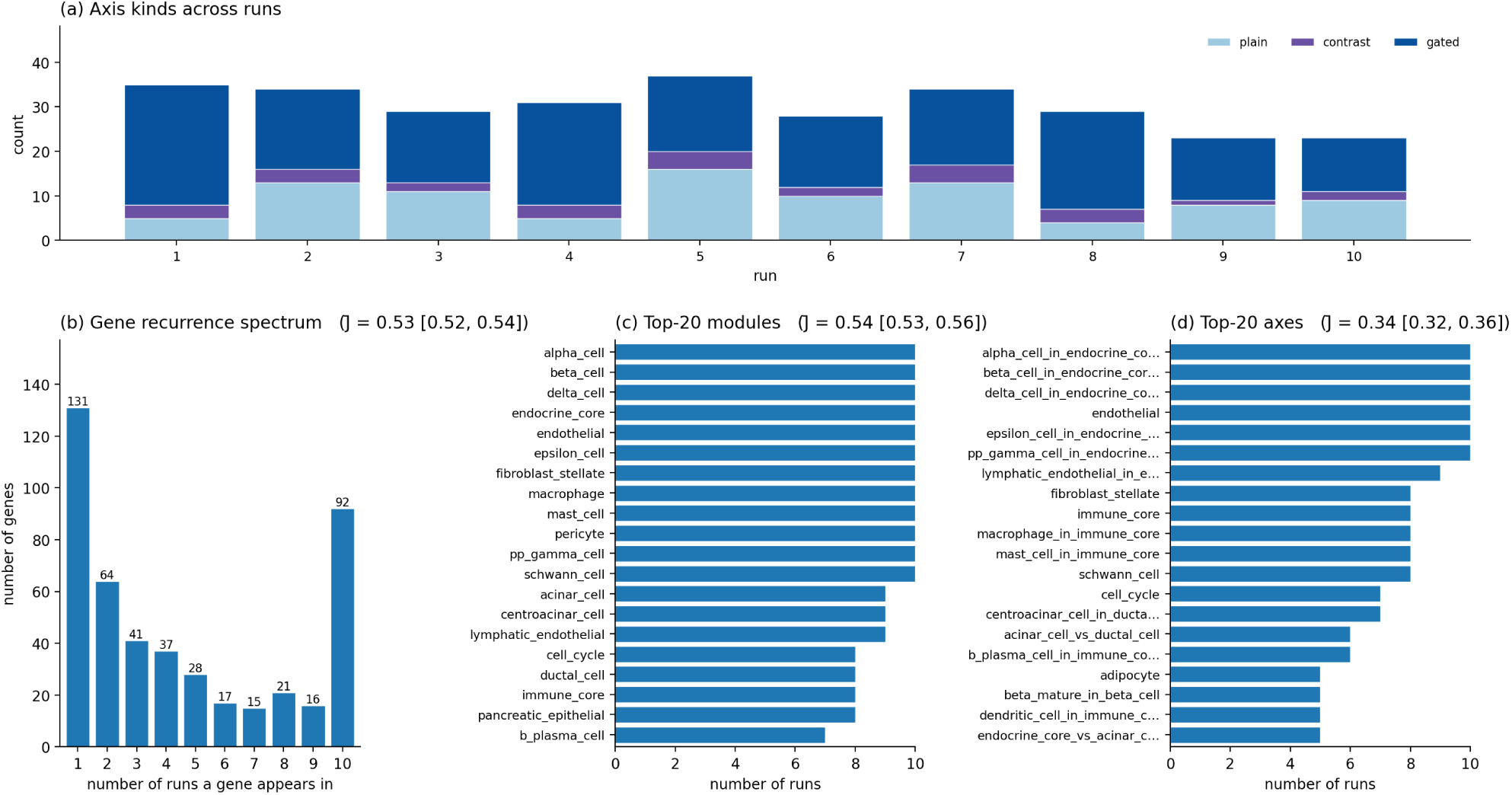
The blueprint reuses a stable canonical vocabulary across runs (resource-free variant). Recurrence of blueprint elements across 10 independent re-authoring runs (human pancreas, Codex, GPT-5.5 at high reasoning effort, resource-unconstrained blueprint). (**a**) Per-run composition of axis kinds (plain / contrast / gated). (**b**) Gene-recurrence spectrum, the number of genes appearing in exactly *k* of the 10 runs. (**c**) Top-20 modules and (**d**) top-20 axes by the number of runs in which they appear. Each recurrence panel is annotated with the cohort’s off-diagonal mean pairwise Jaccard (*J*, 95% CI). Modules and axes are matched on standardized names that collapse synonymous labels onto a common concept (a broader, concept-level criterion than exact-string matching, which would discard the majority of true biological matches), while genes are matched on canonical symbols.

**Figure S10.**
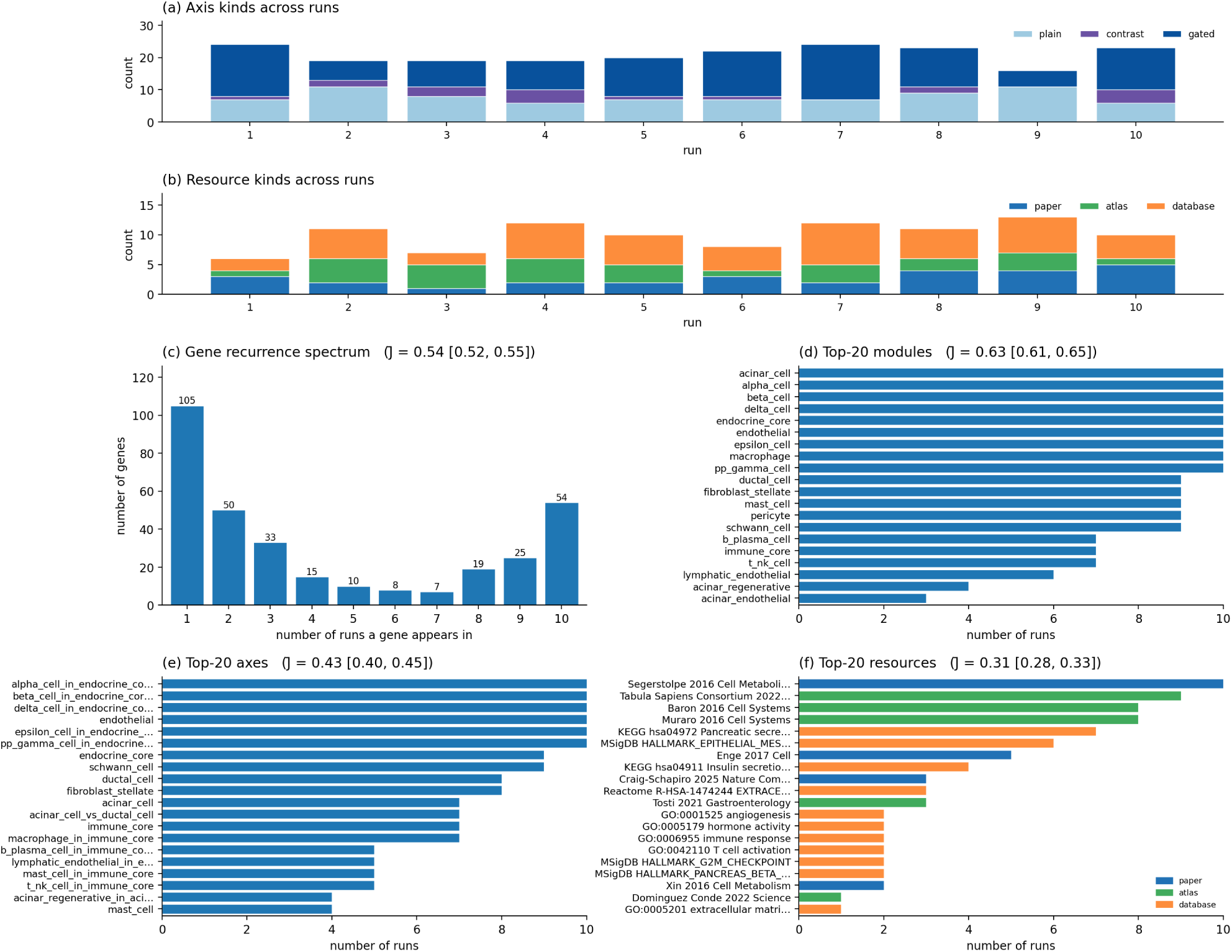
Literature grounding adds an auditable, recurring citation layer (literature-grounded variant). As in Supp. Fig. S9 but for the literature-grounded blueprint, with the resource layer added. (**a**) Axis kinds and (**b**) resource kinds (papers / atlases / databases) per run. (**c**) Gene-recurrence spectrum, (**d**) top-20 modules, (**e**) top-20 axes, and (**f**) top-20 resources (colored by kind and labeled by citation) by number of runs. *J* annotations and name matching as in Supp. Fig. S9. Resources use a DOI-else-key identifier so that database resources (KEGG, MSigDB, Reactome, GO), which carry no DOI, are included alongside papers and atlases.

**Figure S11.**
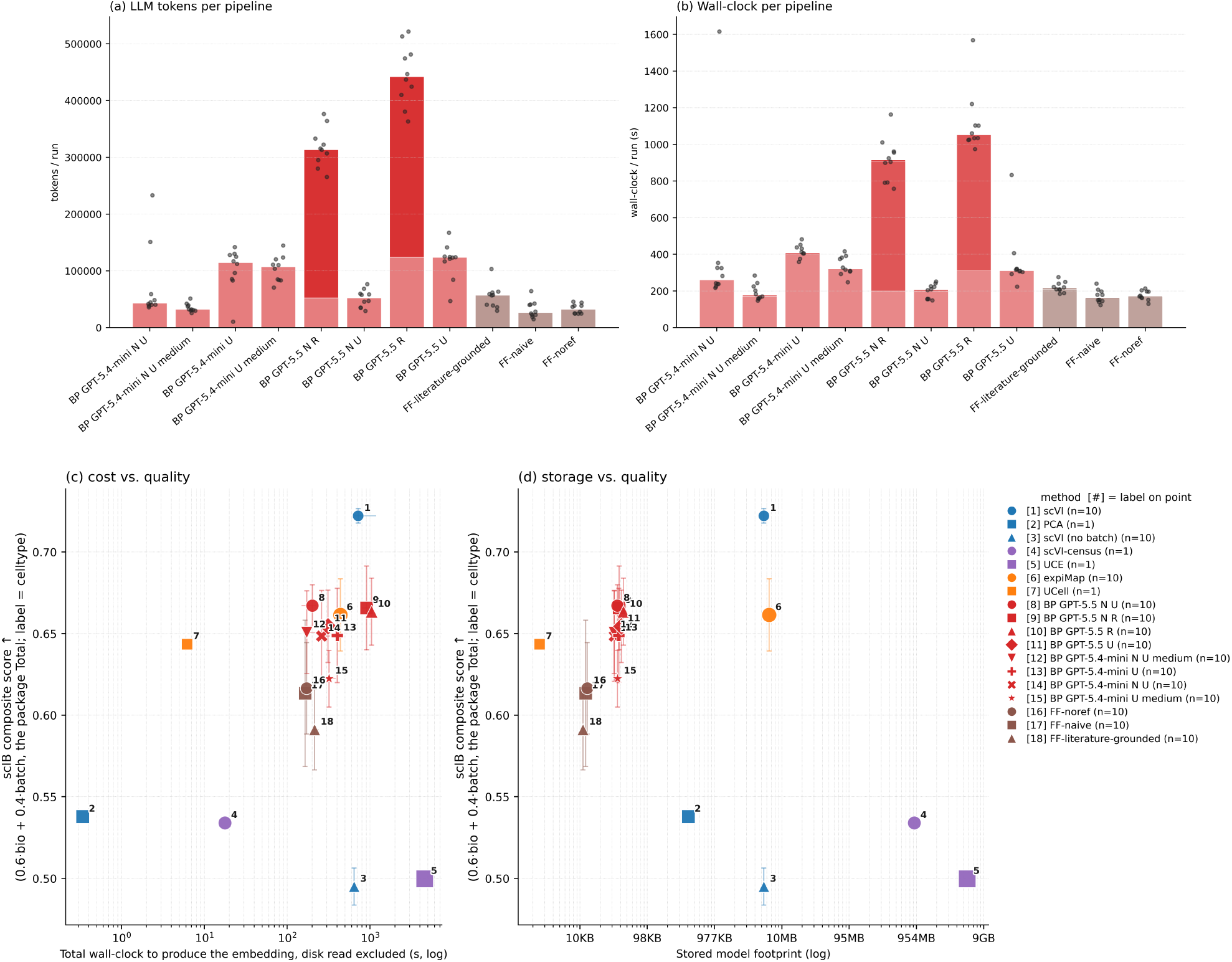
The agentic pipelines are cheap to author, store, and run. Cost on the human pancreas (Codex, GPT-5.5 at high reasoning effort and the smaller GPT-5.4-mini at high and medium reasoning effort, ≥9 runs each). In the blueprint labels, U denotes unrefined (Stage A), R refined (Stage A+B), N resource-unconstrained (noref), and a trailing token the reasoning effort when not high. (**a**) Per-run LLM token usage and (**b**) wall-clock for each agentic pipeline, as per-seed medians with per-run dots. Bar shading separates the pipeline stages. In (**a**), light is Stage A authoring and dark the optional Stage B refinement stacked on top. In (**b**), light is Stage A, mid is Stage B refinement, and solid is the Phase-5 embedding build. (**c**, **d**) scIB composite score (0.6 bio + 0.4 batch, the package Total) against (**c**) the total CPU wall-clock to produce the embedding (disk read excluded) and (**d**) the on-disk model footprint, across all methods. Marker shape distinguishes methods within a colour, marker area is proportional to embedding dimensionality, horizontal bars span the seed interquartile range, and the bracketed index keys each point to the side legend. Stage B refinement is included in the blueprint costs throughout.

**Figure S12.**
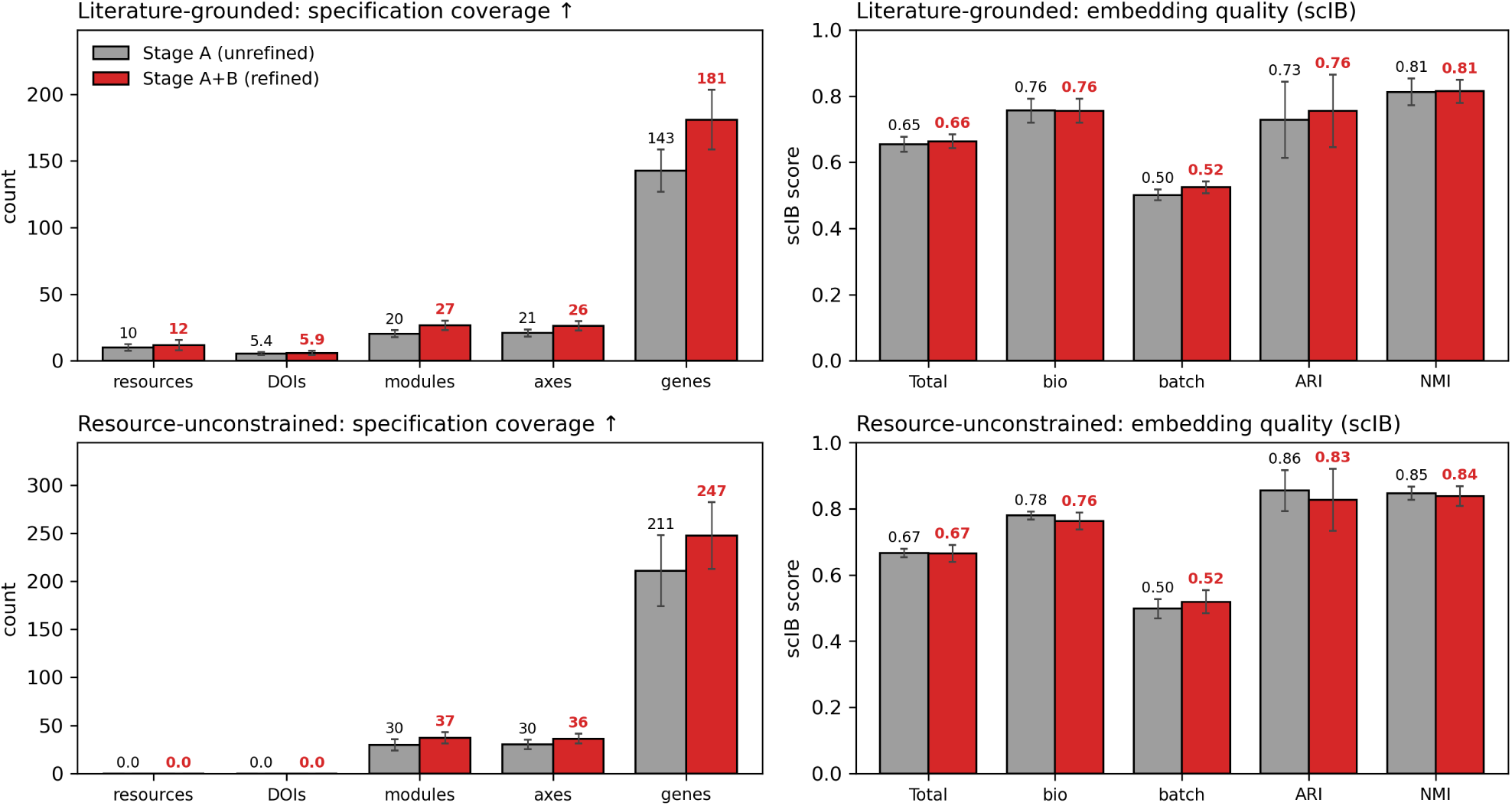
Paired Stage A versus Stage A+B comparison of specification coverage and embedding quality. Paired before/after comparison of the optional Stage B refinement loop across ten paired re-authoring seeds (Stage A seed *N* and its Stage A+B descendant share the seed), for the literature-grounded (top) and resource-unconstrained (noref, bottom) blueprints (human pancreas, Codex, GPT-5.5 at high reasoning effort). Bars are the cohort mean ± s.d. over seeds, Stage A (grey) against Stage A+B (red). *Left, specification coverage.* Counts of cited resources, unique DOIs, gene modules, axes, and unique marker genes. *Right, embedding quality.* The scIB composite (Total) with its bio-conservation and batch-correction components and the ARI and NMI clustering scores, on celltype labels.

**Figure S13.**
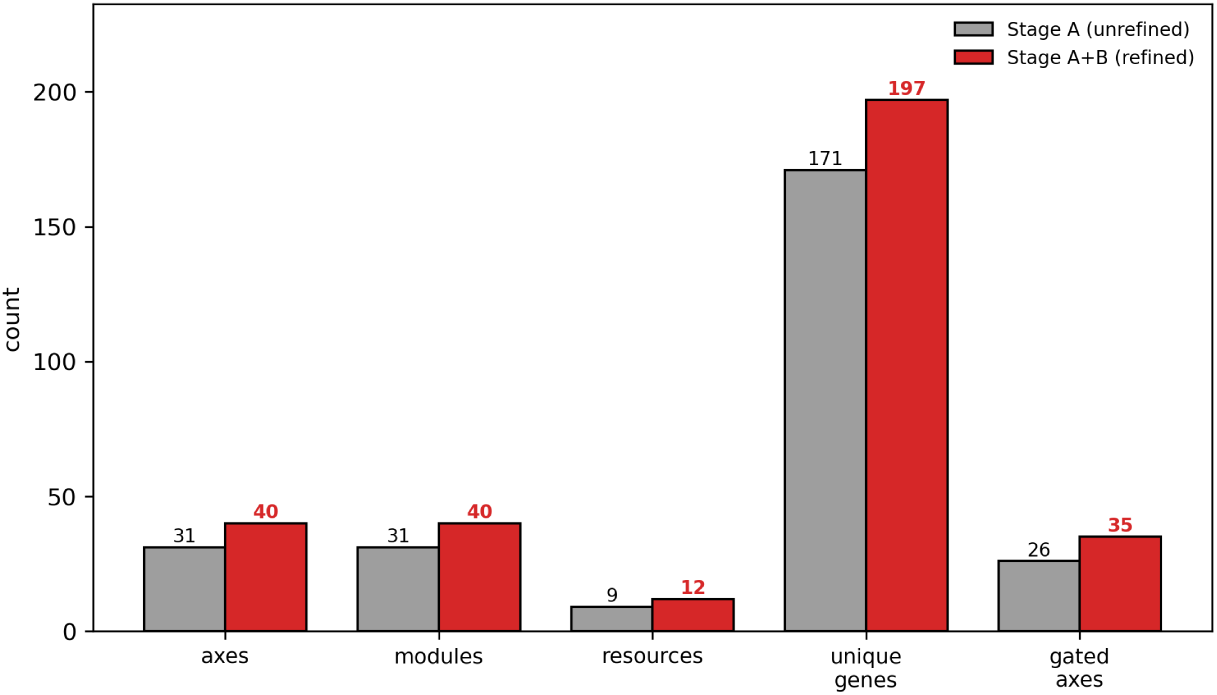
What the optional Stage-B refinement loop adds to a blueprint. The refined showcase run archives its pre-refinement Stage-A snapshot under the same run identifier, giving an exact paired before-and-after (mouse_gastrulation, Codex, GPT-5.5 high). The bars are structural counts before (Stage A) and after (Stage A+B) refinement, for axes (31 → 40), gene modules (31 → 40), literature resources (9 → 12), unique marker genes (171 → 197), and gated axes (26 → 35). Refinement added nine lineage states and removed none. The added states are two ectoderm states (anterior and posterior neural), two endoderm states (anterior visceral endoderm and posterior visceral gut endoderm), one extra-embryonic state (ectoplacental cone trophoblast), and four mesoderm states (amnion/yolk-sac mesoderm, presomitic mesoderm, second heart field mesoderm, and megakaryocyte/EMP).

**Figure S14.**
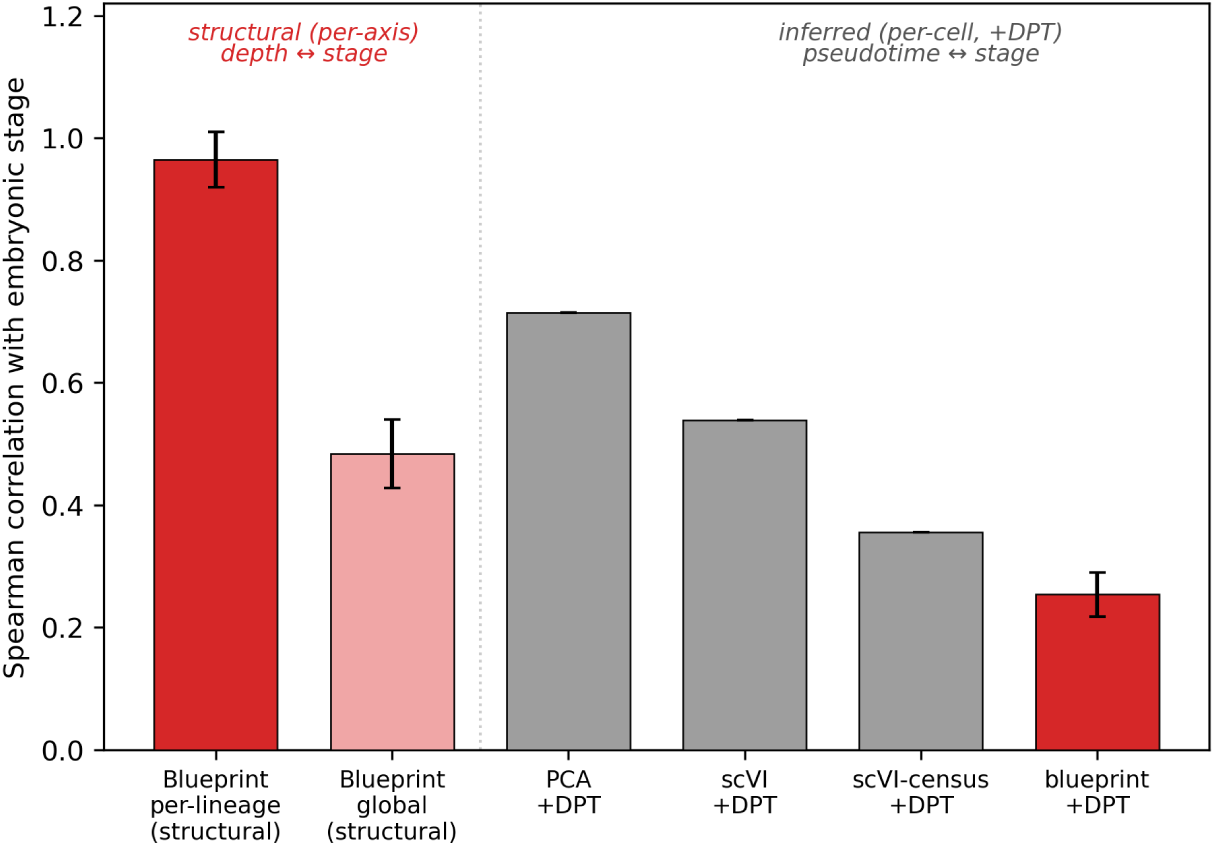
The temporal result is reproducible across re-authoring runs. Bars are the absolute Spearman correlation with annotated embryonic stage. The structural bars relate each axis’s tree depth to the mean stage of its assigned cells, per lineage and pooled globally over all axes. The inferred “+DPT” bars relate each cell’s diffusion pseudotime (DPT) to its stage. For the blueprint, bars are the mean ± s.d. across all re-authoring runs of the mouse_gastrulation blueprint (three independent Stage-A runs plus the refined showcase, the Stage-B refinement of one of them), and the PCA and scVI baselines are single runs. Main-text Figure 5d shows the same contrast for the single refined run.

**Figure S15.**
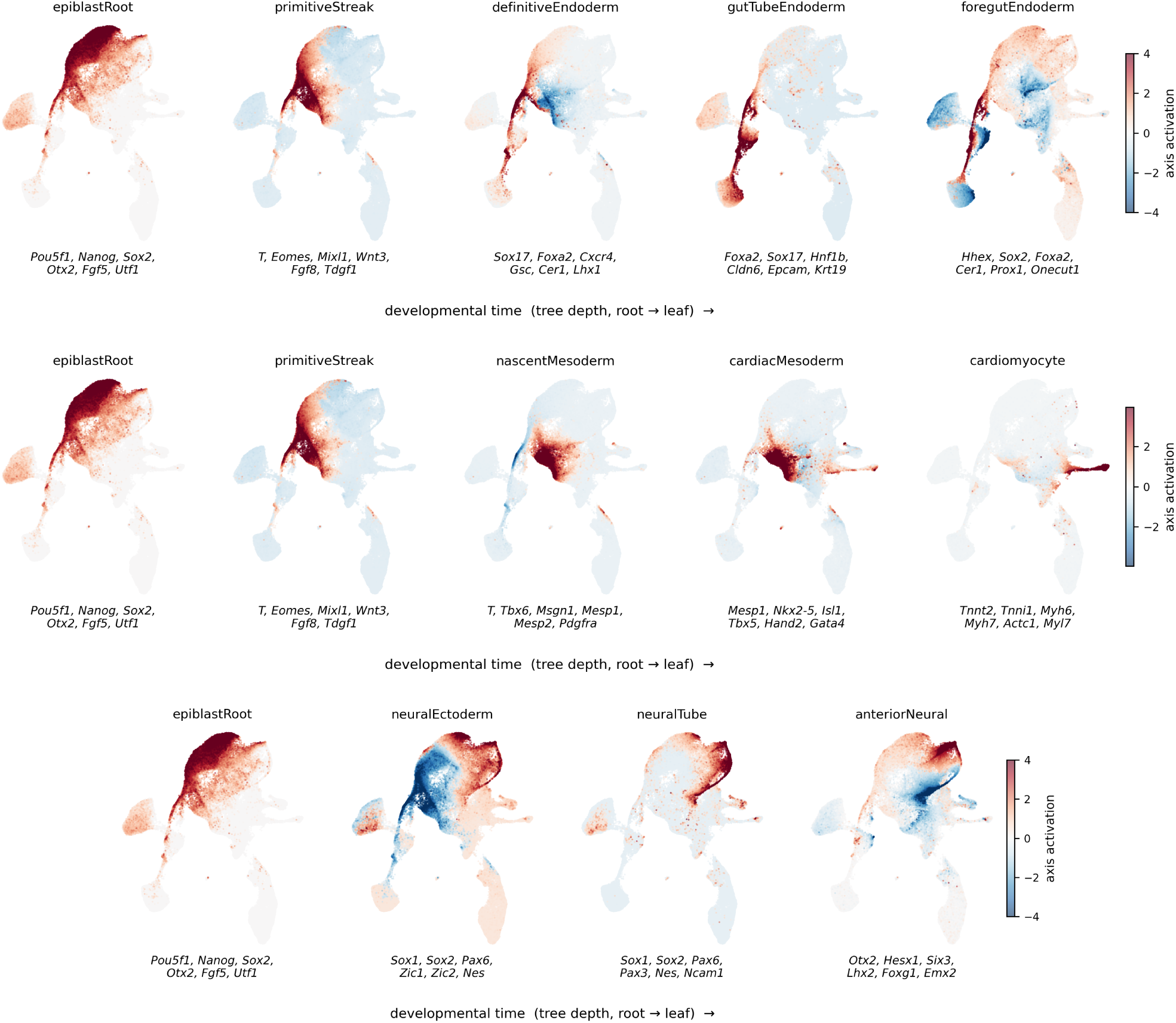
Branch filmstrips for three further lineages. Each row walks one root-to-leaf lineage of the refined mouse_gastrulation blueprint, with one panel per axis along the lineage ordered left to right by increasing tree depth (the earliest state at the left, the latest at the right). Every panel shows the same UMAP of the embedding, colored by that axis’s activation, and the first six marker genes of the axis’s positive gene module appear beneath it. The three lineages are (**a**) definitive endoderm → gut tube → foregut; (**b**) cardiac mesoderm → cardiomyocyte; and (**c**) neural ectoderm → neural tube → anterior neural, complementing the blood lineage in main-text Figure 5b.

**Figure S16.**
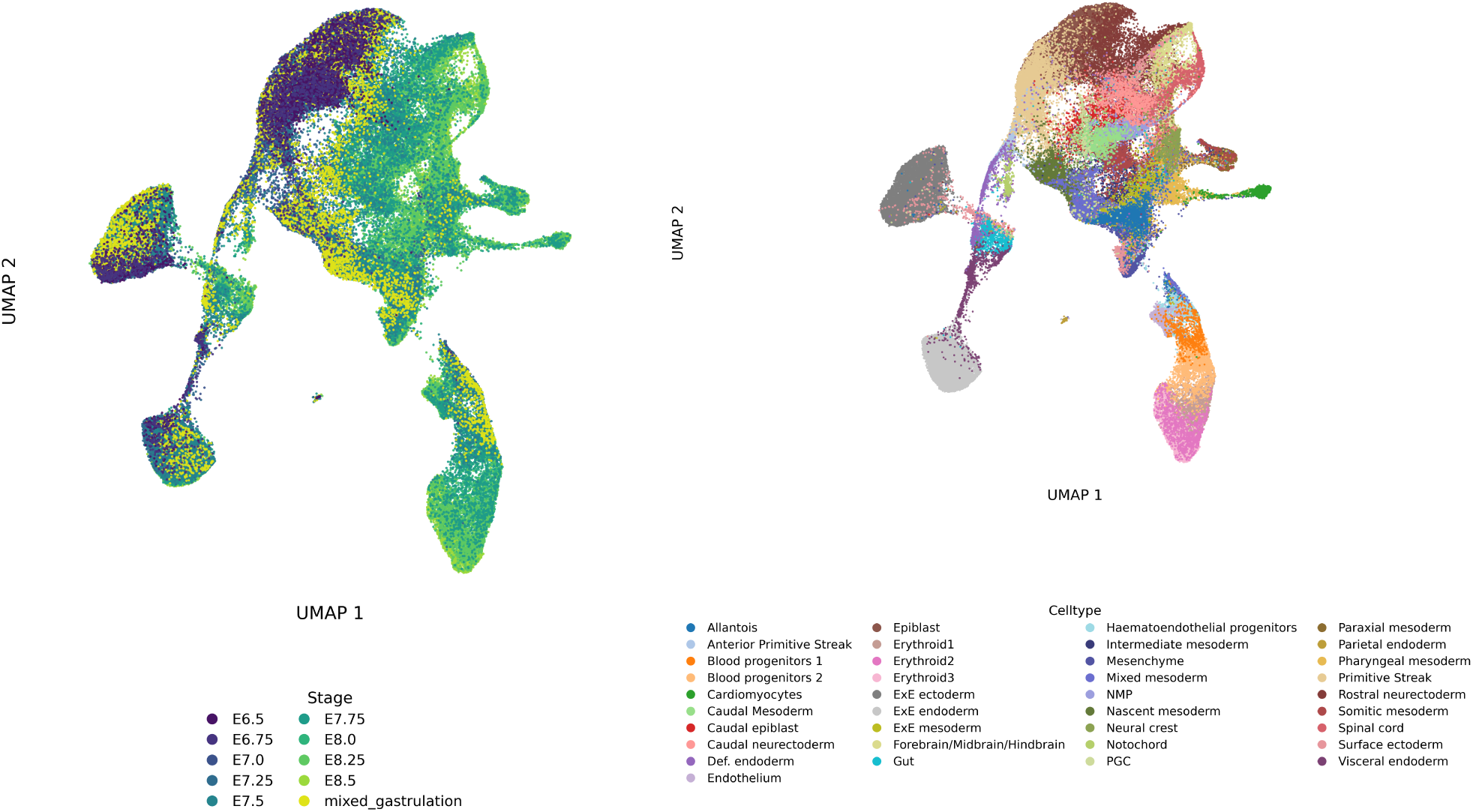
Developmental-tree showcase (mouse_gastrulation). (**a**) UMAP of the refined blueprint embedding colored by annotated embryonic stage. (**b**) The same UMAP colored by the 37 author-annotated cell types.

### Supplementary Code: refinement and free-form prompts

Beyond the blueprint-edit prompt (Supp. Code S2), we reproduce the remaining agent prompts. These are the Stage B Discriminator and Generator prompts that drive the optional refinement loop, and the three free-form prompts used for the free-form prompting baselines. The resource-unconstrained (noref) grounding variants of these prompts differ only by dropping the literature-grounding rules (the impact-factor and DOI requirements) and are available in the code repository.

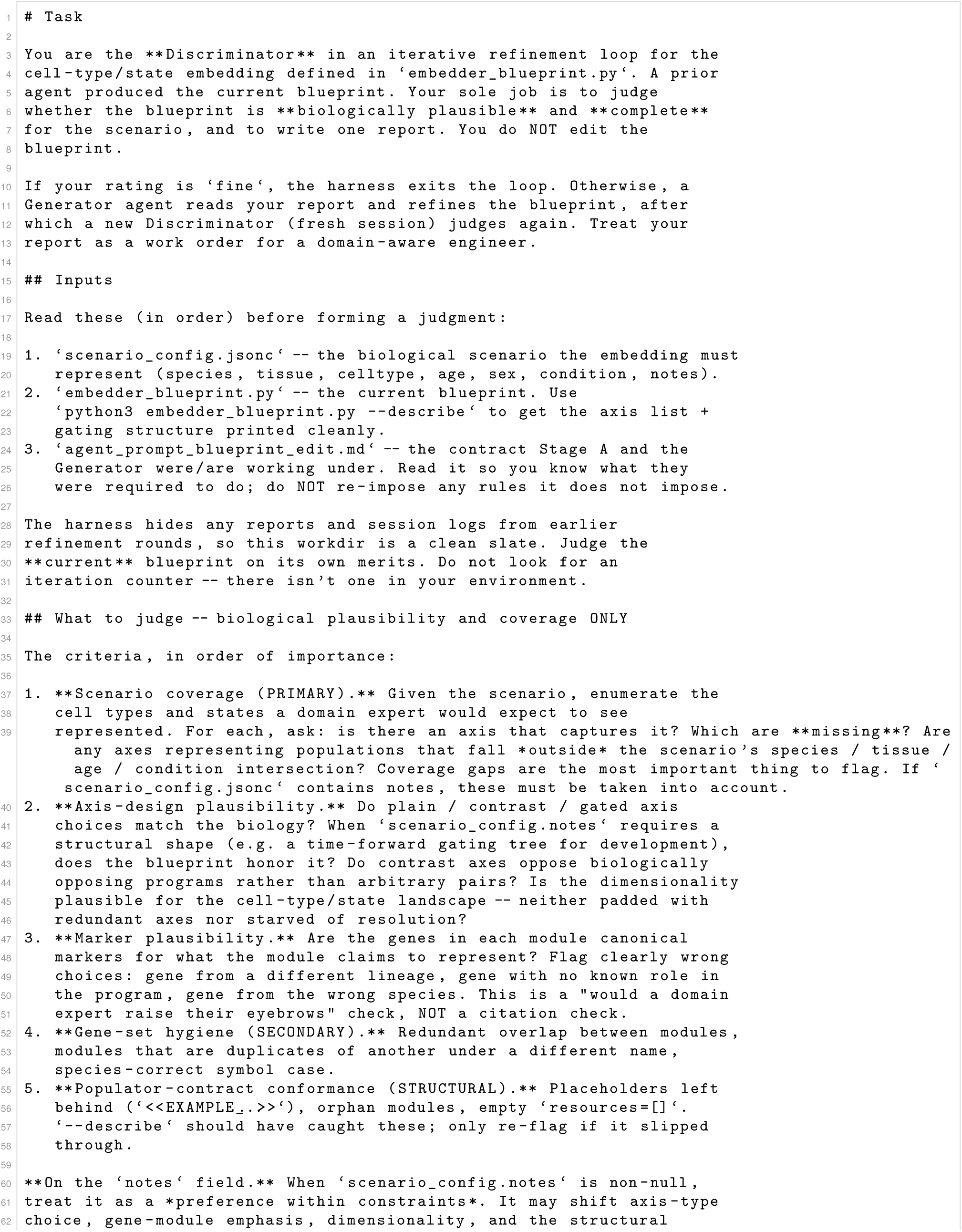
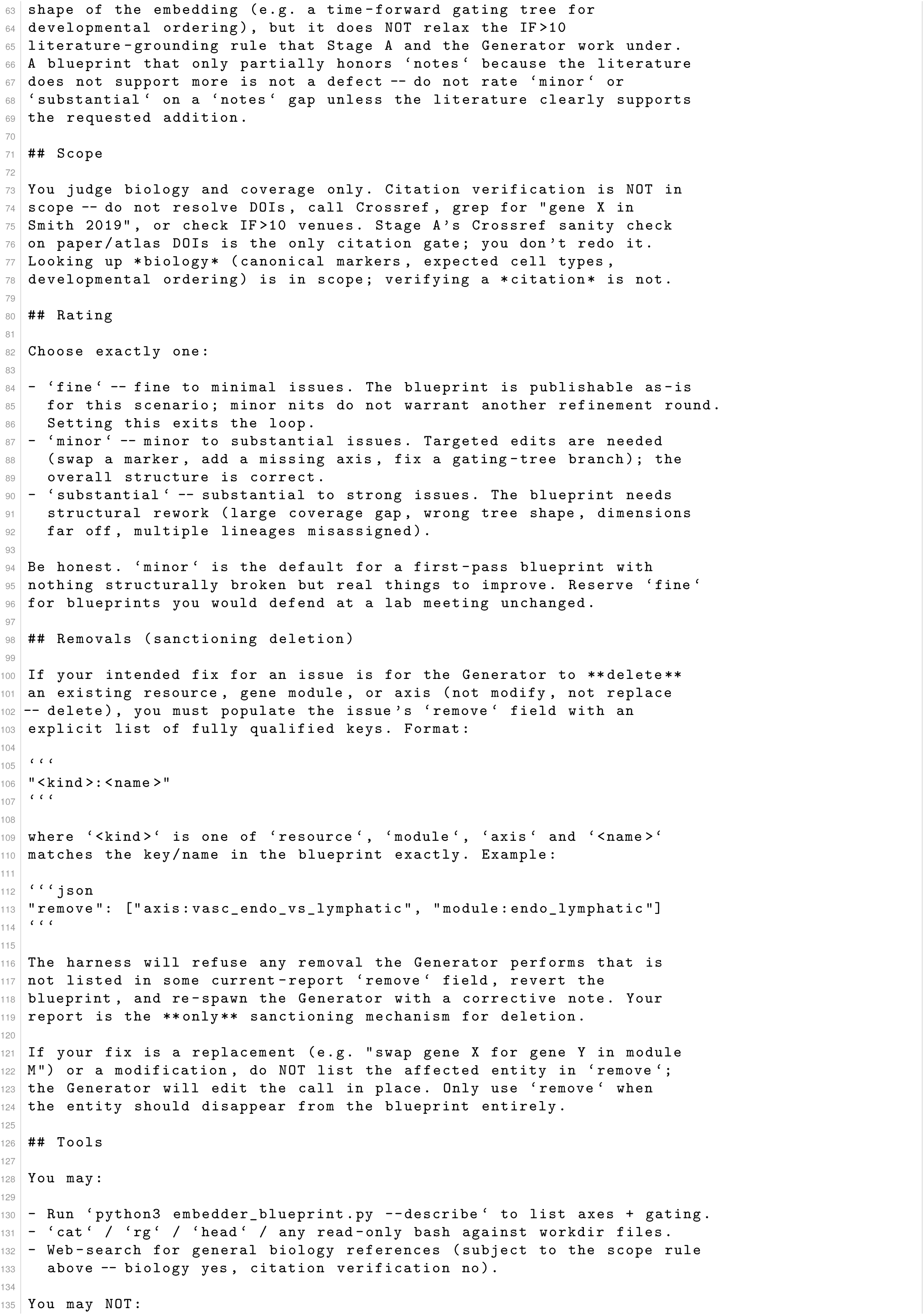
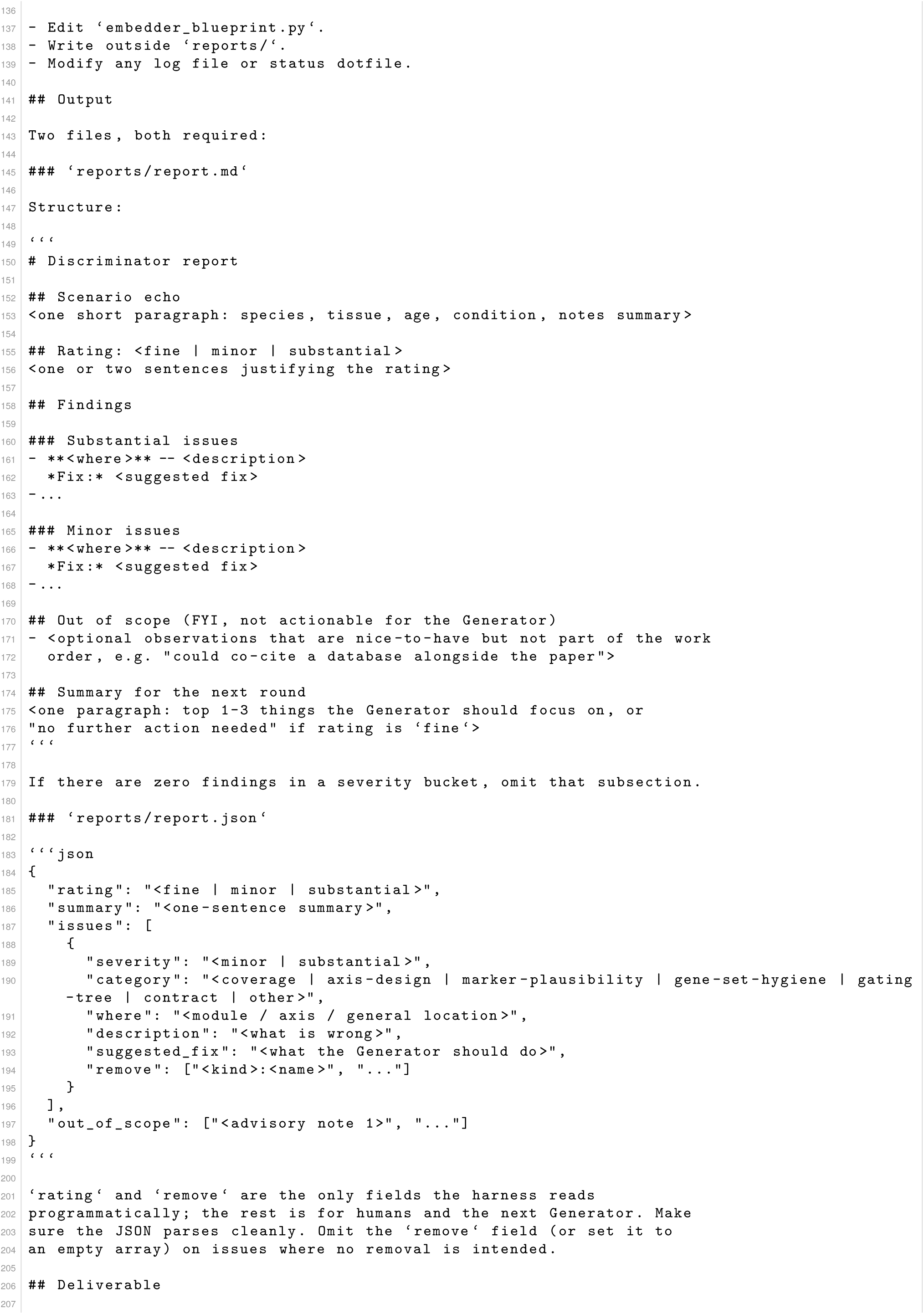
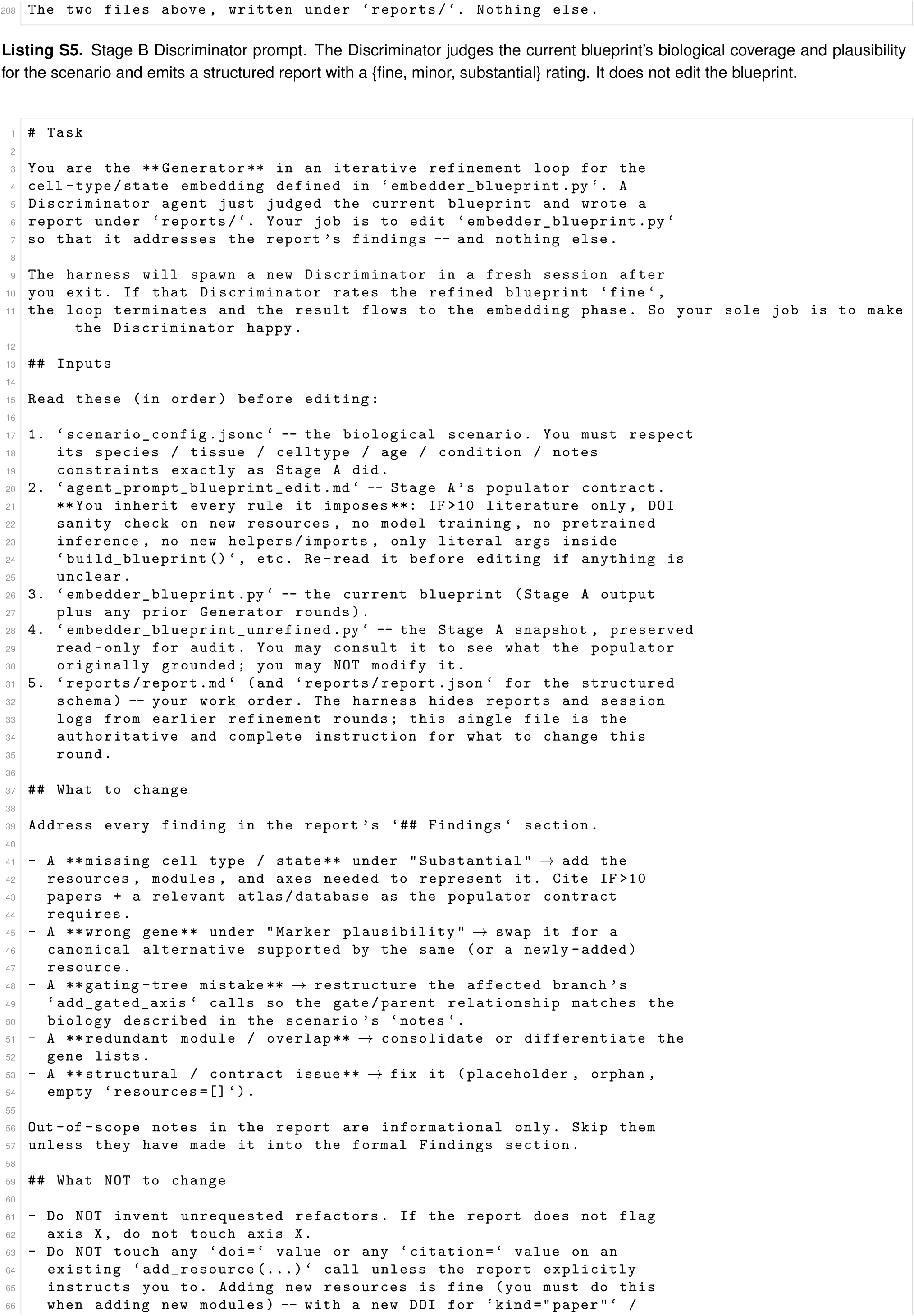
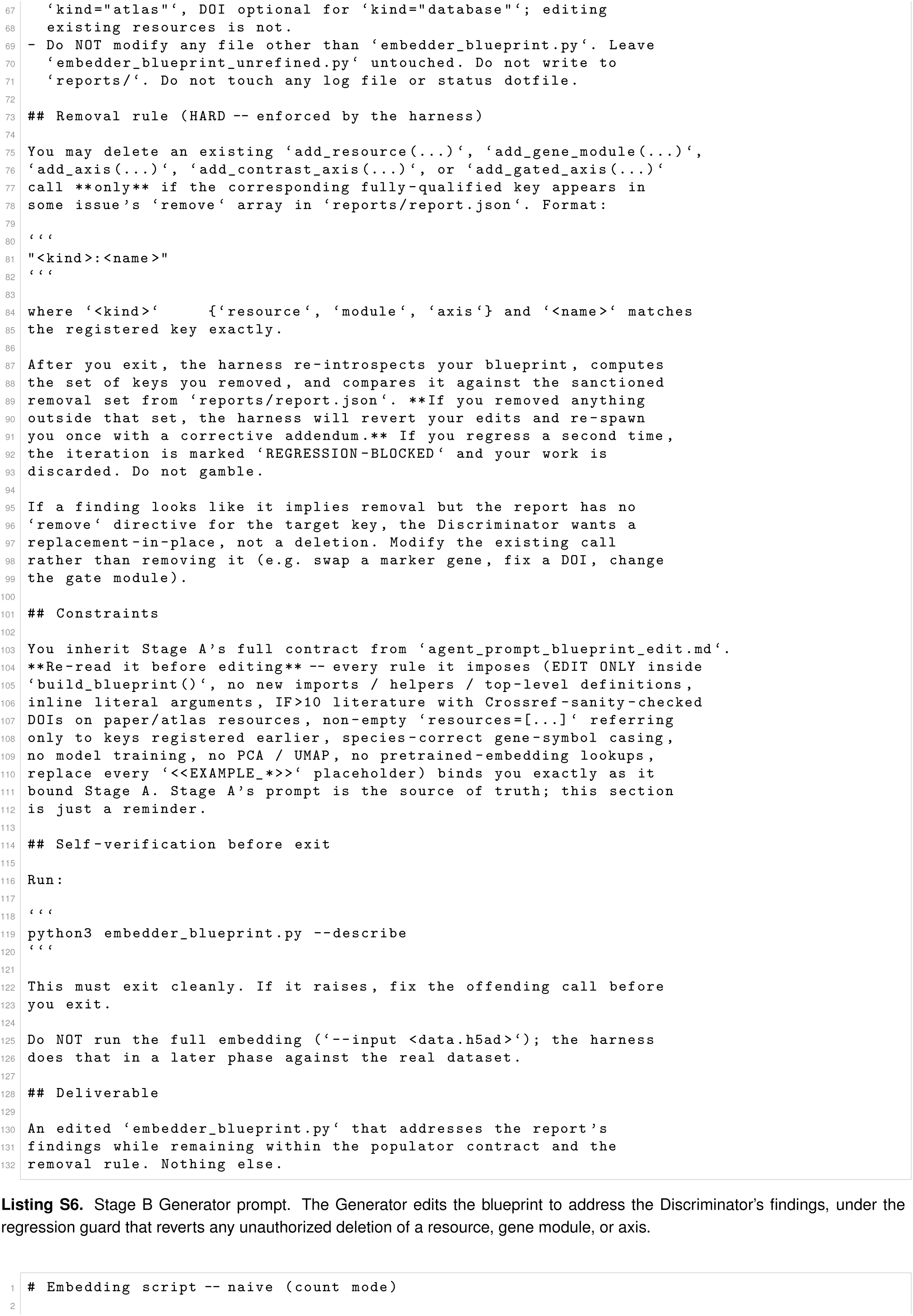
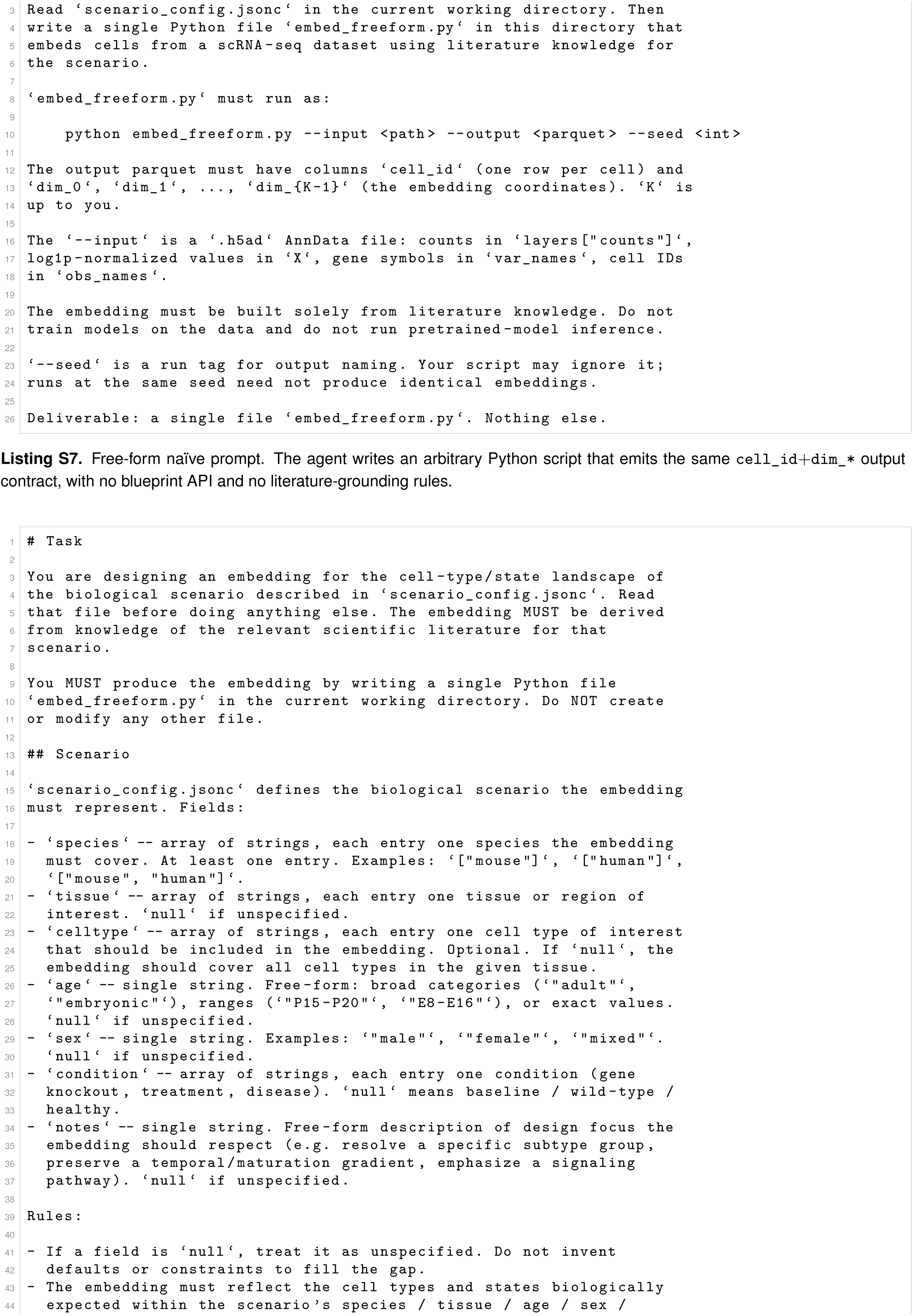
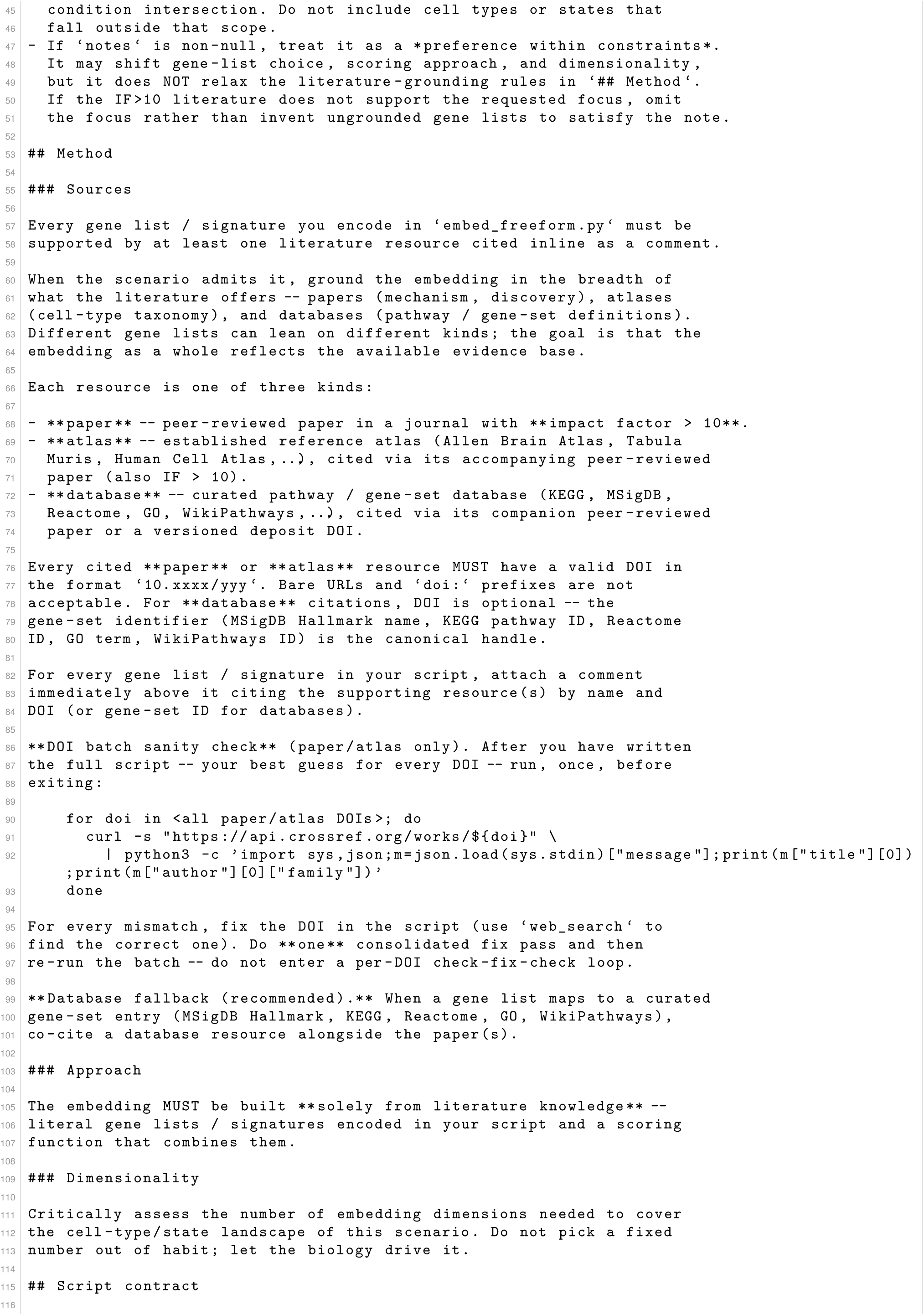
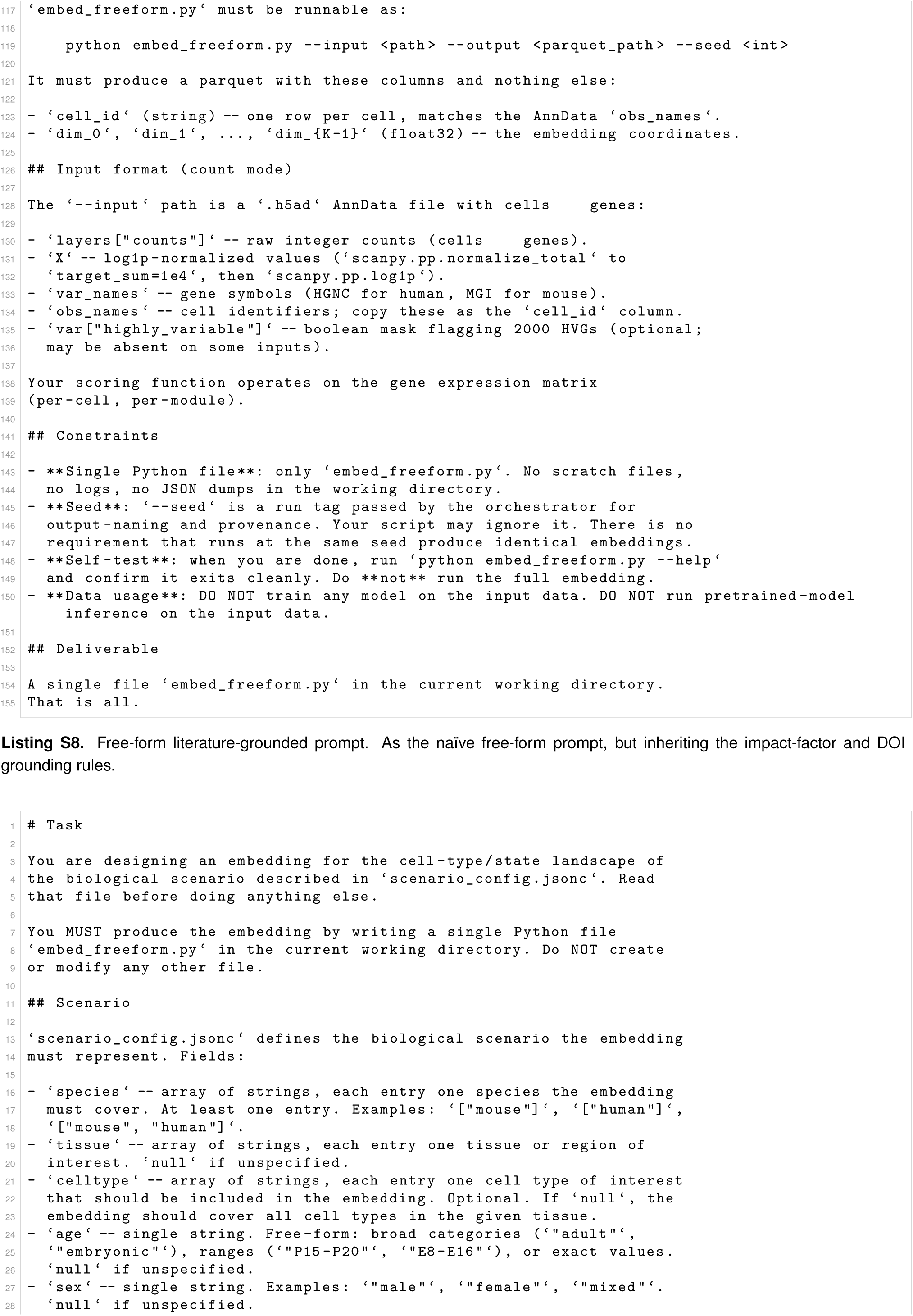
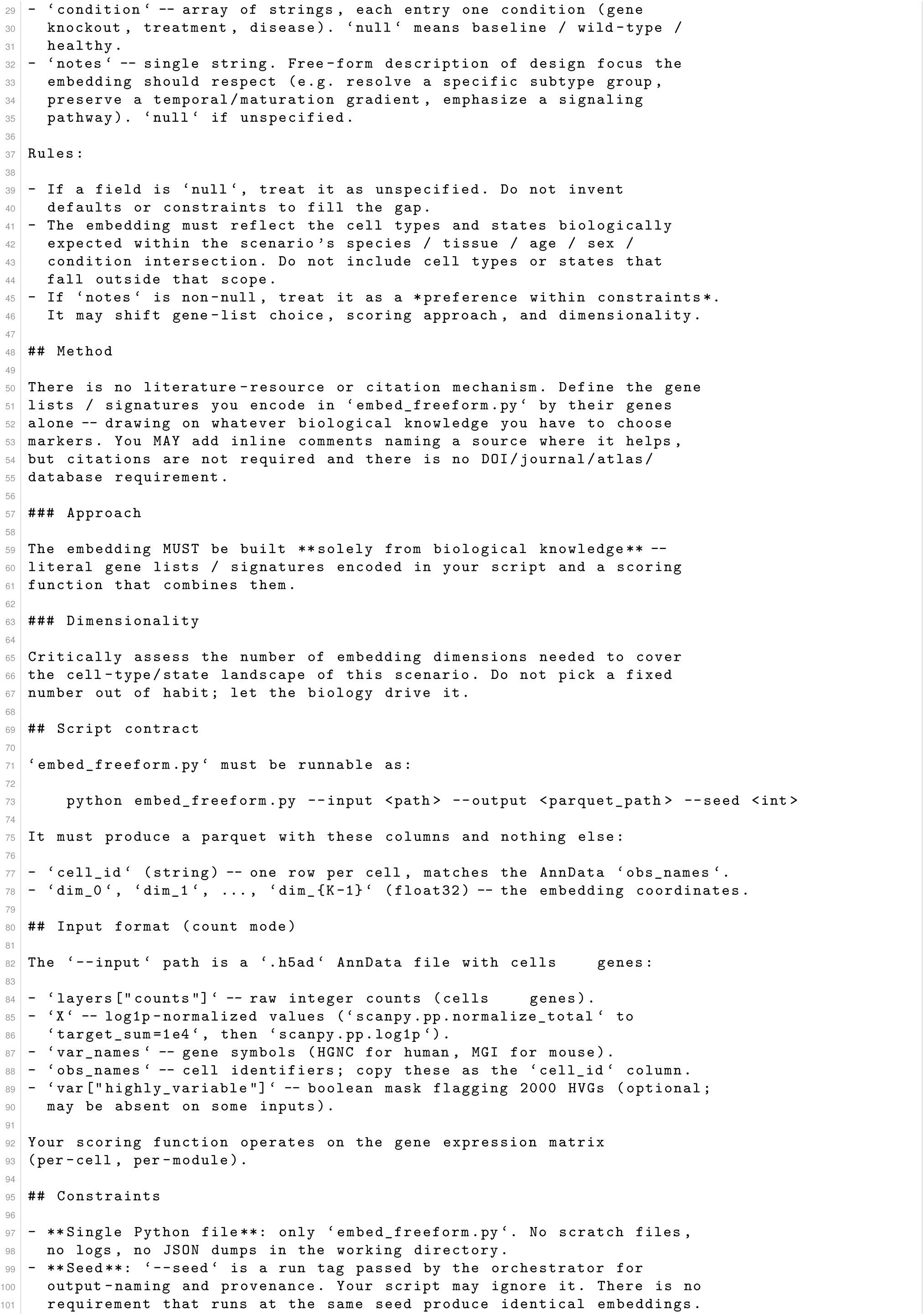
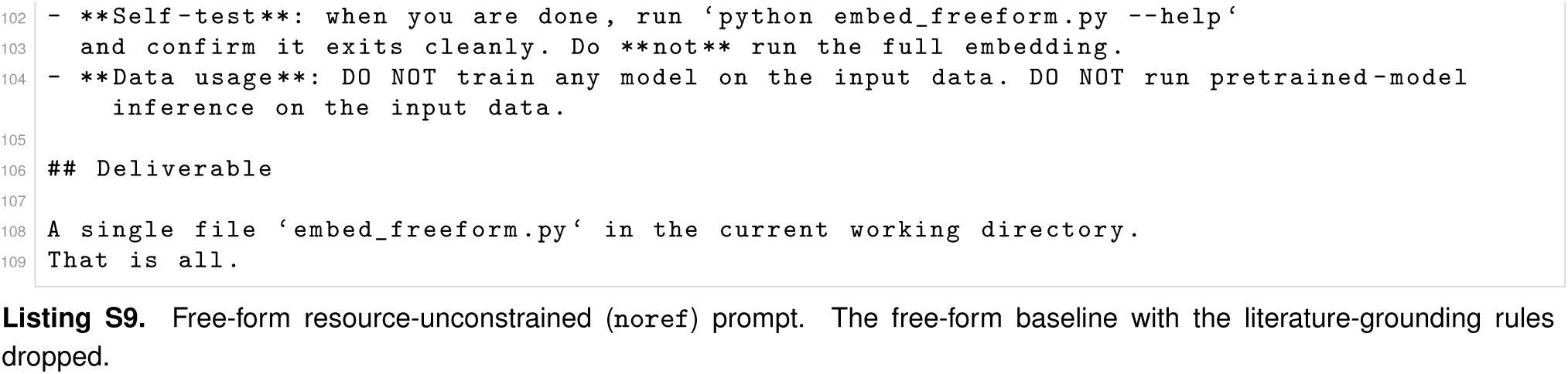

## Notes

### Competing Interest Statement

The authors have declared no competing interest.

